# Early cellular and molecular signatures correlate with severity of West Nile Virus infection

**DOI:** 10.1101/2023.05.14.540642

**Authors:** Ho-Joon Lee, Yujiao Zhao, Ira Fleming, Sameet Mehta, Xiaomei Wang, Brent Vander Wyk, Shannon E. Ronca, Heather Kang, Chih-Hung Chou, Benoit Fatou, Kinga K. Smolen, Ofer Levy, Clary B. Clish, Ramnik J. Xavier, Hanno Steen, David A. Hafler, J. Christopher Love, Alex K. Shalek, Leying Guan, Kristy O. Murray, Steven H. Kleinstein, Ruth R. Montgomery

## Abstract

Infection with West Nile Virus (WNV) can drive a wide range of responses, from asymptomatic to flu-like symptoms/fever or severe cases of encephalitis and death. To identify cellular and molecular signatures distinguishing WNV severity, we employed systems profiling of peripheral blood from asymptomatic and severely ill individuals infected with WNV. We interrogated immune responses longitudinally from acute infection through convalescence at 3 months and 1 year employing multiplexed single cell protein and transcriptional profiling (CyTOF and Seq-Well) complemented with matched serum proteomics and metabolomics. At the acute time point, we detected both an elevated proportion of pro-inflammatory markers in innate immune cell types and reduced frequency of regulatory T cell activity in participants with severe infection compared to those with asymptomatic infection. Single-cell transcriptomics of paired samples revealed that asymptomatic donors had higher expression of genes associated with innate immune pathways, in particular anti-inflammatory CD16^+^ monocytes at the acute time point. A multi-omics analysis identified factors--beyond those from individual analyses--that distinguished immune state trajectory between severity groups. Here we highlighted the potential of systems immunology using multiple cell-type and cell-state-specific analyses to identify correlates of infection severity and host cellular activity contributing to an effective anti-viral response.

## INTRODUCTION

West Nile virus (WNV) is a global mosquito-borne pathogen which can cause severe neurological illness, especially in immunocompromised and older individuals (1, 2). Since being introduced to the United States in 1999, the virus has spread through North and South America (3) and more than 51,000 cases have been reported, including 2,390 fatalities (4, 5). By 2016, infections in the USA were estimated at 7 million people (6). While about 80% of infected people have an asymptomatic infection, approximately 20% experience mild illness with fever symptoms, and a minority of cases (∼1%) experience severe, neuroinvasive disease (7).

To understand the broad range of clinical symptoms and determinants of susceptibility to severe WNV disease, we can consider elements of the pathogen, vector, and host. The dominant circulating strain of WNV in North America since 2001 (WN2002) and the proportion of severe cases reported across the country have remained fairly constant over two decades (3–5). Thus even though environmental variation in climate, mosquito, and bird populations are relevant for the frequency of exposures (8), variations in host responses contribute significantly to the range of disease severity in human cases.

A number of host factors are relevant for susceptibility to severe WNV. These include genetic analyses that identified variations in several immune-related genes in individuals at increased risk for severe disease (9–11) including those involved in type I interferon (IFN) pathways which are critical for control of WNV infection (12). The critical role of type I IFN pathway has been confirmed in experimental studies (13, 14) as have immune pathways involved in recognition and clearance of WNV, including RIG-I mediated immune responses (1, 7, 15–17). Notably, age is a critical risk factor for WNV and the majority of severe cases are over age 60 (5, 8). Aging is associated with a well-documented decline in immune recognition and signaling by pathogen recognition receptors such as Toll-like receptors (TLRs), RIG-I (retinoic acid-inducible gene I), and RLR (RIG-like receptors), highly relevant for responses to infection with WNV and type I IFN responses to viral infections such as WNV and influenza and responses to vaccination (18–23). But while symptomatic WNV infection occurs predominantly in older people (24), there is no significant difference with age among asymptomatic participants, underscoring the critical role of individual host response mechanisms (25).

Investigating transcriptional and functional immune responses in blood of recovered WNV patients, we previously identified a susceptibility signature in innate immune cells including interferon pathways, production of the chemokine CXCL10, and a key role for reduced induction of pro-inflammatory cytokine interleukin (IL)-1β in severe infection (26). IL-1β has also been shown to be critically important for control of WNV in the brain (27). In addition, symptomatic patients have been reported with reduced numbers of T cell subsets (Th1/Th17 and Tregs) and elevated expression of Tim-3^+^ and atypical T cells (28–30). Notably, antibodies against WNV were durable and were not different between asymptomatic and symptomatic adults for titer or epitopes targeted (31). However, studying samples collected after infection resolves, we cannot distinguish whether features may be a cause or a consequence of clinical severity. Taking advantage of advances in systems immunology, and an unusual recruitment strategy, we investigated immune responses during acute infection from asymptomatic and symptomatic participants using multiple omics profiling platforms and integrating high resolution methods of cellular, transcriptomic, proteomic and metabolomic analyses. Comparing stratified participants longitudinally at acute and convalescent timepoints, we have taken advantage of stable characteristics of immune cell frequencies and gene expression in individuals (32–34) to identify molecular signatures and cell states and integrated temporal responses associated with resistance to severe WNV infection. These signatures may provide relevant guidance for timely therapeutic responses in severe infections.

## RESULTS

### WNV study population for longitudinal immunophenotyping profiles

To investigate immune-related differences in infection with WNV contributing to the divergence in the clinical course and identify responses related to severe infection, we enrolled two groups of WNV participants in Houston, TX. Hospitalized patients had acute CDC-defined WNV infection including clinical criteria of mild symptomatic response or severe neurologic disease, with no specific treatments for WNV (35). We enrolled acutely infected asymptomatic participants found to be viremic while volunteering to donate blood. Our unusual recruitment criteria allowed us to enroll both severity groups at comparable duration of viral exposure (∼ 7 days). Immune signatures from asymptomatic (WNA) and symptomatic (WNF fever and WNE neurologic) study participants were assessed at enrollment (Day 1) and at follow-up time points of Month 3 and Year 1 of convalescence (n=11; **Table 1**).

**Table 1.** Demographics of study participants.

| Parameter | Asymptomatic<br>(n=4) | Symptomatic<br>(n=7) | Total<br>(n=11) | P value* |
| --- | --- | --- | --- | --- |
| **Mean age (yr) (SD)** | 67.8 (7.4) | 52.9(14.0) | 58.3 (13.9) | 0.0471 |
| **range** | 59-77 | 28-68 | 28-77 |  |
| **No. (%) of females** | 0 (0) | 2 (29) | 2 (18) | 0.0215 |

*Age: Using t-test, two-tailed.*
*Gender: Using Chi-square test*

### Immune cell frequency and functional status in WNV infection

For profiling frequency and functional status of immune cells, we assessed whole blood samples using the SMART Tube system to stabilize cell phenotypes (36). Untreated cryopreserved samples were profiled using CyTOF, employing a 40-marker antibody panel to broadly quantify both immune cell lineages and functional markers (**Table S1**). To reduce experimental variation, all samples were assayed together using a uniform lot of reagents and including a spike-in reference with each sample as described previously (37). Data from the reference cells partitioned from WNV samples was assessed by PCA. No evidence of batch effect from the reference cells was observed across all symptom status or time points of sample collection (**Fig. S1**).

Cell types from whole blood samples were manually annotated based on characteristic lineage markers (**Fig. 1A**) using a defined cell ontology (**Fig. S2B**) (38). Using a whole blood sample allows a detailed examination of 29 cell subsets including polymorphonuclear leukocytes (PMN), the most abundant immune cell type in circulation, and 28 subsets of peripheral blood mononuclear cells (PBMCs). Many infection-related differences in cell frequency were detected in both asymptomatic and symptomatic groups. Analysis of cell frequencies for each individual showed an elevated frequency of PMN in total cells at acute viral infection compared with his/her own convalescent sample at Month 3 time point (**Fig. 1B**; Day 1 vs Month 3 p=0.003 for asymptomatic; p=0.0002 for symptomatic). No significant differences for frequency of PMN were noted between severity groups (**Fig. 1B**). We have previously shown a biphasic role for PMN in murine WNV infection with an initial phase permissive for viral replication and contributing to dissemination and subsequently later in infection controlling viremia (39). Within PBMCs other innate immune cell types also showed significant elevations at the acute time point reflecting expected increases during an acute viral infection (37, 40). Levels of CD14^+^ classical monocytes in PBMCs were elevated at Day 1 for all participants compared to each individual’s Month 3 and Year 1 of convalescence (**Fig. 1B**; acute vs Month 3 p=0.0199 for asymptomatic; p=0.009 for symptomatic). Overall, we detected elevated frequencies > 1.1-fold for 18 of 28 PBMC cell subsets during acute infection compared to convalescent time points, including a marked elevation in the frequency of memory B cells (Bm), which may be expected as antibody responses develop (**Fig. 1B**; Day 1 vs Month 3 p=0.0015 for asymptomatic; p<0.0001 for symptomatic). All subject groups showed elevated levels of activated CD8^+^ T cells (CD8+ T) at Day 1 compared to convalescent timepoints (**Fig. 1B**; Day 1 vs Month 3 p=0.0013 and 0.0005 for asymptomatic; p<0.0001 for symptomatic). A similar pattern was noted in activated CD4^+^T cells (Day 1 vs Month 3 p=0.0013 for asymptomatic; p<0.0001 for symptomatic). These differences are expected in symptomatic participants given the important role for activated CD8^+^ T cells in response to viral infections, related to their recognized functional activities in viral killing and clearance and production of inflammatory mediators. However, our data reveals comparable increased frequencies in asymptomatic donors which might not have been predicted given the absence of clinical symptoms.

**Figure 1.**
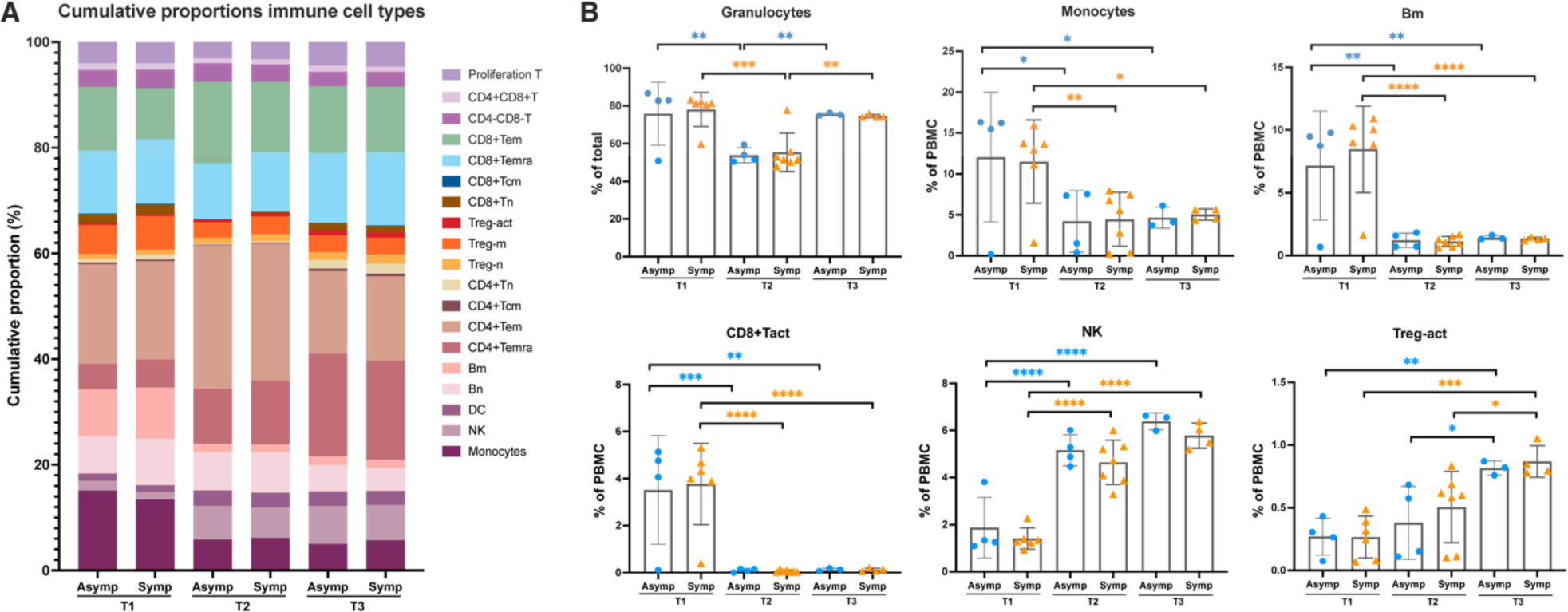
Frequency of immune cell subsets of WNV infected participants. Blood samples were collected into SMART tubes from WNV asymptomatic (n=4) and symptomatic (n=7) participants at acute Day 1 and convalescent time points at Month 3 and Year 1. Whole blood cells were labeled for immune cell markers and profiled by CyTOF in one batch. Frequency of PMN and 28 PBMC subsets were defined by expression of lineage markers (Fig. S2B). (A) Frequency of 28 PBMC subsets shows the variation of cell frequencies in asymptomatic and symptomatic groups at the timepoints indicated for cell types shown (CD4+ and CD8+ subsets for naïve (n), effector memory (em), central memory (CM), activated (act), and effector (Temra). (B) Differences in frequencies of cell subsets between cohorts and time points were assessed by one-way ANOVA with Tukey’s multiple comparisons test (* p<0.05, ** p<0.01, *** p<0.001, and *** p<0.0001). No significant frequency differences were found between asymptomatic and symptomatic groups.

While many cell types were more abundant during acute infection, in contrast, we detected significantly lower frequencies at Day 1 for Natural Killer (NK) cells (**Fig. 1B**; p<0.0001 for asymptomatic; p=0.0002 for symptomatic). We previously showed that a more immature subset of NK cells is highly responsive to WNV with degranulation and cytokine production (41). Significant increases over the time course were also detected for levels of activated Tregs (Treg-act) which have an important role in resolution of inflammation (**Fig. 1B**; Day 1 vs Year 1 p= 0.0028 for asymptomatic; p=0.0002 for symptomatic). These increased levels over time are consistent with previous reports for symptomatic WNV patients (28) and notably were equivalent between asymptomatic and symptomatic participants. Overall, quantifying frequencies of multiple immune lineages showed consistent changes longitudinally in cell type abundance whether or not participants experienced symptomatic illness. This reveals ongoing ‘shadow’ immune responses in asymptomatic individuals and suggests that cell frequency alone is not sufficient to distinguish immune responses leading to severe disease.

In our phenotyping profile, we quantified 20 markers of cellular activation to assess functional changes which may contribute to susceptibility to severe infection. This multiparameter platform allows us to interrogate each marker in 29 cell subsets for a total of 580 marker/cell type combinations. In response to infection with WNV, we detected elevated proinflammatory activity in multiple cell types at the acute time point with levels decreasing at Month 3 and Year 1 (**Fig. 2A, Fig S3A, S3B**). Remarkably, numerous activation markers were altered equivalently across subject severity cohorts with 451/580 combinations being higher at Day 1 compared to Month 3. Highly expressed markers at the acute time point included L-selectin adhesion molecule CD62L, marker of activation of PMN and monocytes and an important factor for lymphocyte trafficking out of the blood (42). Elevated levels of CD41, the integrin glycoprotein IIb/IIIa, were also detected in multiple cell lineages during acute infection, which may reflect interactions with platelets which are readily apparent in the whole blood samples being examined here. This finding is consistent with well-characterized binding of platelets to activated leukocytes (43). In addition, CD41 was expressed on precursor cells which may be more abundant during acute infection (44). The marker CD279 (PD-1) was elevated at the acute compared to Month 3 time point, which may be important for development of CD8^+^ T cell memory responses (45). These elevations in immunomodulatory markers resolved and were not significant between groups at Month 3 or Year 1 (**Fig. S3**).

**Figure 2.**
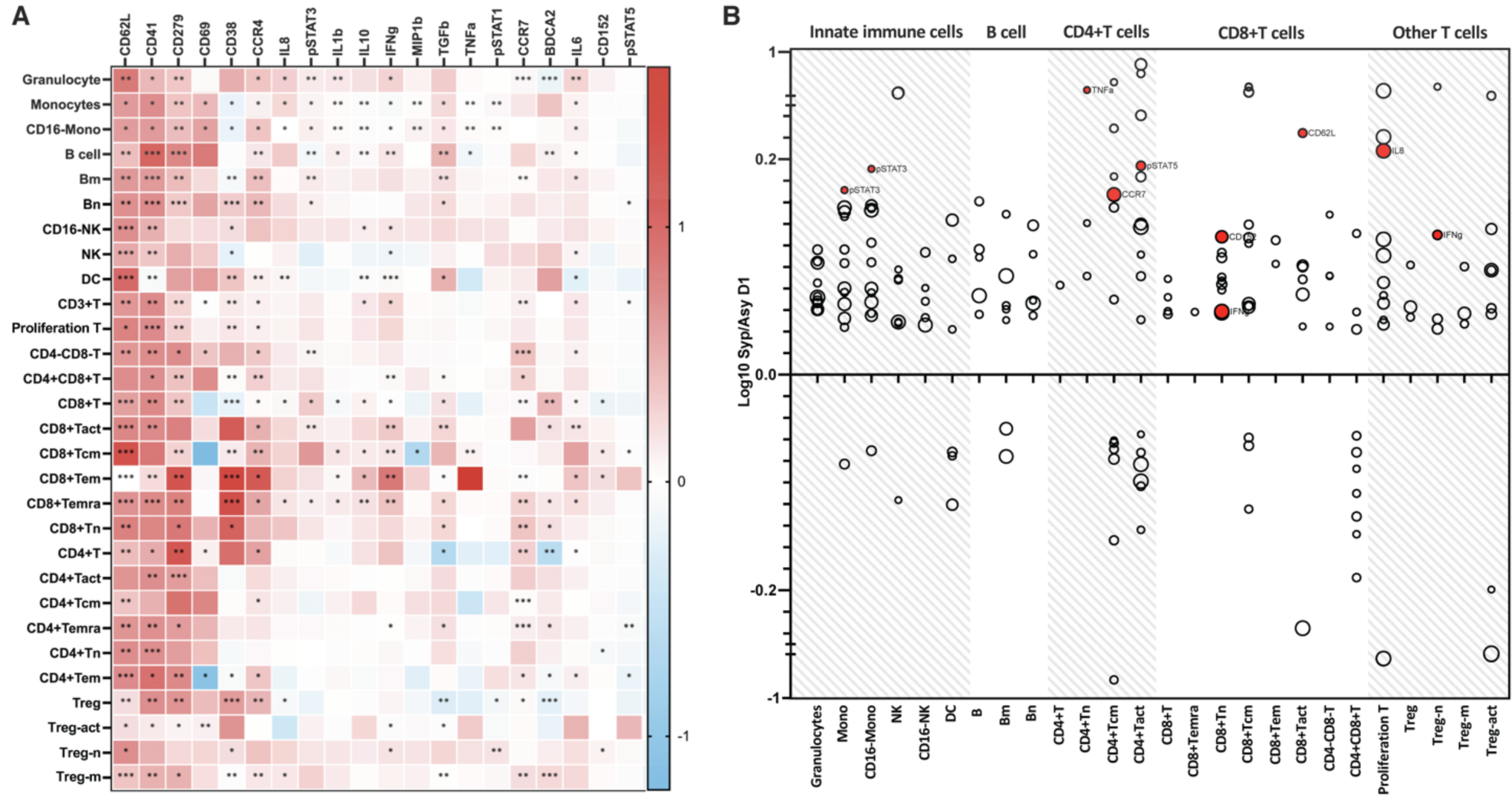
Altered immune functional markers during acute WNV infection. Whole blood cells from WNV patients (n=11) were labeled with metal-conjugated antibodies and analyzed by mass cytometry to quantify levels of 20 functional markers in 28 cell subsets. (A) Change in median channel intensity at acute vs Month 3 convalescent time point (ratio log10 acute/convalescent). Significance by Generalized Linear Mixed Model with * p<0.05, ** p<0.01 and *** p<0.001. (B) Log10 ratio of Symptomatic/Asymptomatic of marker/cell type combinations shown for individual cell subsets with fold change of greater than 10% of baseline where size of circle reflects the average expression level of a marker in a cell type. Statistically significant combinations in permutation analysis are shaded in red, where the observed log ratio of expression in symptomatic relative to asymptomatic occurred less than 5% of the time in the permuted distribution.

To investigate functional differences contributing to the divergence in the clinical course, we identified marker/cell type combinations that were significantly different between symptomatic and asymptomatic participants. Of the 274 cell type/marker combinations above a threshold of 20% of baseline of mean channel intensity, we found 181 cell type/marker combinations expressing intensity differences >10% between severity groups (**Fig. 2B**). At the acute time point in symptomatic participants, we detected significant elevations for combinations of pro-inflammatory functional markers in innate immune cell types, including elevated expression of cytokines (IL-1β, MIP1β, IL-6, IL-8, TNFα) in PMN and in monocytes; phosphorylated signaling intermediates (pSTAT1 and pSTAT3) and the inflammatory marker CD62L, known as L-selectin, in PMN, monocytes, NK and DC (**Fig. 2B**, p=0.038 and 0.024 for pSTAT3 in monocytes and CD16^-^ monocytes). Elevations in functional markers in adaptive cell types in symptomatic participants include TNFα in naïve CD4+T cells (p=0.049), CCR7 in CD4^+^ T cells (p=0.006), pSTAT5 in activated CD4^+^ T cells (p=0.012), and numerous markers in CD8^+^ T cells including CD62L, CD152, and IFNγ and CD69, the C-lectin receptor marker of lymphocyte activation increased in several subsets of T cells (CD4^+^ Tem, CD4^+^Tact cells, and naïve Treg) (**Fig. 2B**, p=0.042/0.034 for IFNγ/CD152 in naïve CD8^+^ T cells; p=0.022 for CD62L in activated CD8^+^ T cells). The elevated expression in symptomatic patients of phospho STAT3 in monocytes and phospho STAT5 in CD4+ T cells identifies key pathways increasing production of inflammatory cytokines such as IL-6. An elevated inflammatory milieu has previously been reported to contribute to permeability of the blood brain barrier (46), thus the skewing towards pro-inflammatory responses in symptomatic participants may contribute to severity of infection. A lower inflammatory profile in asymptomatic participants may be beneficial for antiviral responses. Comparing longitudinally for each individual, no significant differences were detected between asymptomatic and symptomatic participants at convalescent time points suggesting that key differences in clinical trajectory occur during acute infection.

### Transcriptional profiles of immune response to WNV infection

To collect additional in-depth measurements of cellular function during WNV infection and investigate functional differences contributing to the divergence in clinical course, we assessed transcriptional profiles as an additional measure of functional status to track changes in cell responses during infection with WNV. We used PBMCs collected at the same time points from our study participants to identify differential expression patterns longitudinally and relevant to severe infection. PBMCs were assessed using the Seq-Well platform for efficient single-cell RNA-seq (47). PBMC samples provided 5,243 transcripts and 1,776 genes per cell on average for 56,858 cells sequenced and pre-processed. Cell types were identified by unsupervised clustering and differential expression analysis together with 36 canonical lineage marker genes in an iterative manner using the Seurat R package (**Fig. 3A**; **Fig. S4**) (48). The Uniform Manifold Approximation and Projection (UMAP) displays 11 cell types with innate cell lineages (conventional and plasmacytoid dendritic cells (cDCs, pDCs), monocytes, Natural Killer, NK) separated from T cell subsets and B cells (**Fig. 3A**). Notably, UMAP shows separation of the 3 severity groups by time points--acute (T1), Month 3 (T2), and Year 1 (T3)--where samples from the acute time point clustered together most closely and later time points show considerable mixing among severity groups indicating the transcriptional similarity of cells. As expected, the frequencies of cell types defined by gene expression are consistent with but not identical to cell frequencies defined in CyTOF by protein expression markers, showing slight differences between the two platforms. These differences may be due to mRNAs which are not transcribed, or protein markers which may be shed or secreted on activation, or to differences in whole blood vs PBMC samples.

**Figure 3.**
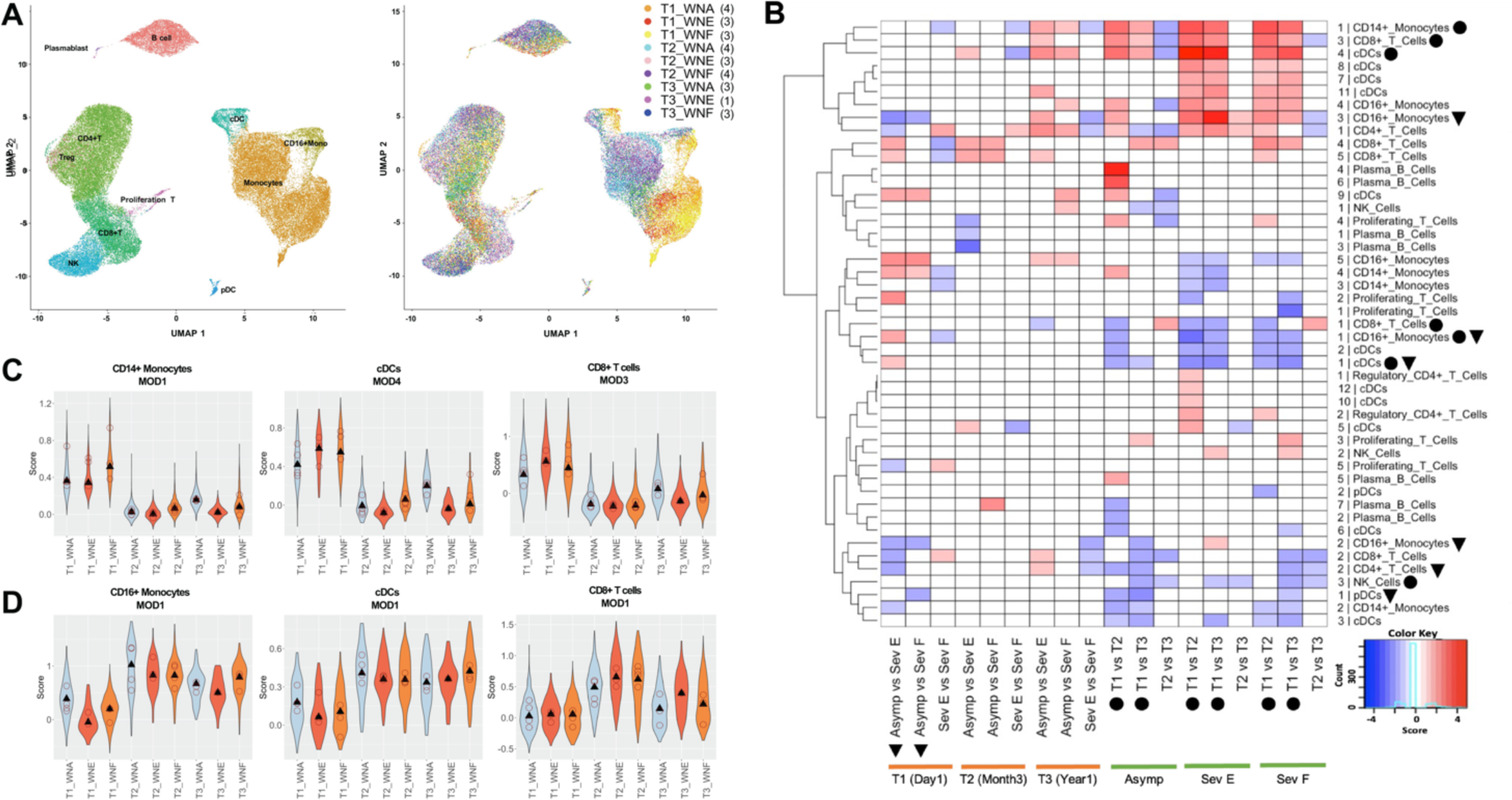
Seq-Well transcriptomic analysis of response to WNV infection. (A) UMAP projection shows distribution of 11 cell types and 9 cell states. WNA = Asymptomatic, WNE = encephalitis, WNF = fever, T1 = Day 1, T2 = Month 3, and T3 = Year 1 with the number of participants in parentheses for each respective condition. (B) Heatmap of signed module scores for 42 differential modules and 18 condition pairs for which there are significant differences with fold change > 1.3 and mean p-value < 0.01. For a given condition pair in each column, positive (red) and negative (blue) reflect a higher module score in the first condition score or second condition, respectively. Condition pairs are marked for significant differences between time points (circle; 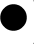) and between severity groups (triangle;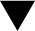). (C) Violin plots of module scores for 3 differential modules of interest elevated at the T1 time point. The subject-level median values in each condition are shown in open circles. (D) Violin plots as in (C) decreased at the T1 time point.

### Gene modules reveal transcriptional responses to WNV infection

Using expression data for the 11 cell types, we used the weighted gene co-expression network analysis (WGCNA)-based module discovery (49) to first identify *significant modules* of co-expressed genes for each of the 11 cell types. We compared all possible condition pairs to identify *differential modules* for the 18 condition pairs. There were 107 significant modules identified from 10 cell types (excluding B cells), from which we identified 47 *differential modules* with fold change > 1.3 and mean p-value < 0.01 in at least one condition pair (**Fig. 3B**; Table 2) with both elevated and reduced gene expression for all severity groups. The numbers of module genes range from 10 in module 2 of pDCs to 161 in module 1 of CD14+ monocytes. We gave particular attention to 12 modules from eight cell types selected by visual assessment of module score distributions showing the most dramatic differences between time points or severity groups as well as the top 10 genes in each module whose expression profiles are most correlated with the module scores (**Table S2**). For temporal signatures, we identified module 1 in CD14+ monocytes, module 1 in CD16^+^ monocytes, modules 1 and 3 in CD8^+^ T cells, modules 1 and 4 in cDCs, and module 3 in NK cells. For signatures reflecting disease severity, we identified modules 1, 2, and 3 in CD16^+^ monocytes, module 2 in CD4^+^ T cells, modules 1 and 4 in cDCs, module 1 in pDCs, and module 6 in plasma B cells. The majority of differential modules revealed signatures related to the time course of infection, with significant changes from the acute time point Day 1 (T1) to the convalescent time point at Month 3 (T2) or Year 1 (T3). For example, the module scores and marker expression were higher in T1 than T2 and T3 in all severity groups for module 1 in CD14^+^ monocytes, for module 4 in cDCs, and for module 3 in CD8^+^ T cells (**Fig. 3C**).

**Table 2.** Top pathways in Seq-Well differential modules. In-depth analysis for 12 selected differential modules identified using Weighted Gene Co-Expression Network Analysis (WGCNA) framework. Top pathways were identified using Enrichr. *Size*, indicates the number of genes in each module.

| Cell Type | Module | Number of Genes per module | Top Pathways in module |
| --- | --- | --- | --- |
| CD14<sup>+</sup> Monocytes | 1 | 162 | HIF-1; hypoxia; glycolysis; IL-2/STAT5 |
| CD16<sup>+</sup> Monocytes | 1 | 16 | Leishmaniasis; Yersinia infection; Apical Junction; mTORC1 |
| CD16<sup>+</sup> Monocytes | 2 | 11 | Amphetamine addiction; IL-17; TNF-α/NF-kB |
| CD16<sup>+</sup> Monocytes | 3 | 16 | Mineral absorption; Iron metabolism; Apoptosis; IL-2/STAT5 |
| CD4<sup>+</sup> T Cells | 2 | 11 | Osteoclast differentiation; TNF- $\alpha$/NF-kB; Neuroinflammation |
| CD8<sup>+</sup> T Cells | 1 | 35 | Bacterial invasion of epithelial cells; Pathogenic Escherichia coli infection; Interferon Gamma Response |
| CD8<sup>+</sup> T Cells | 3 | 20 | NOD-like receptor signaling; Ferroptosis; TGF-β; TNF-α/NF-kB |
| cDCs | 1 | 109 | Hematopoietic cell lineage; Macrophage markers; Complement system of innate immunity |
| cDCs | 4 | 119 | Ferroptosis; HIF-1; NAD metabolism, sirtuins and aging; TNF-α/NF-kB; Inflammatory response |
| NK Cells | 3 | 15 | Neuroinflammation; TNF-α/NF-kB; hypoxia; TGF-β |
| pDCs | 1 | 11 | IL-17; Neuroinflammation; TGF- $\beta$; TNF-α/NF-κB; hypoxia |
| Plasma B Cells | 6 | 32 | N-Glycan biosynthesis; Interferon gamma response; IL-2/STAT5; TNF-α/NF-κB |

Examination of the upregulated genes in these modules revealed many genes relevant for response to acute viral infection and innate immune pathways. In particular, module 1 in CD14^+^ monocytes, which is upregulated in acute infection, contained 162 genes which are enriched in critical pathways to support cell metabolic function and to initiate immune cell responses (hypoxia and glycolysis, HIF-1, and IL-2 signaling). The top differentially expressed genes in CD14^+^ monocyte module 1 include genes involved in cell attachment and integrin signaling reflecting the need for dynamic rearrangements of the actin cytoskeleton for spatial reorganization of receptors and adhesion molecules: *TIMP1*, a metalloproteinase inhibitor which plays a role in integrin signaling and activates cellular signaling cascades via CD63 and ITGB1; *SDC2*, a heparan sulfate proteoglycan which has been implicated in regulation of cytoskeleton organization and integrin signaling; *THBS1*, thrombospondin, an adhesive glycoprotein that mediates cell-to-cell and cell-to-matrix interactions; and *SERPINB2*, a regulator of cell movement (50). Detailed examination of genes highly expressed in CD14^+^ monocytes module 1 at the acute time point includes (a) *CD86*, a marker of activated dendritic cells; (b) cytokines, chemokines and their receptors (*ILIRN, CCL2, IRF1, IL6R, CLEC5A, CCL7, IFITM10, CD9, CSF1, DOCK4, ILIR1*); (c) signaling elements including *MAP3K8, NFkB1*, and *HIPK2* which we have shown is necessary for interferon response in WNV infection by phosphorylating the transcription factor *ELF4* (51); and (d) *DUSP4* and *DUSP5*, dual tyrosine/threonine phosphatases which regulate immune cell activity by inactivating MAP kinases (52). The 162 genes in CD14^+^ monocyte module 1 reflect key responses needed to mount an immune response to viral infection. Notably, the number of monocytes was highly elevated in frequency at the acute time point (**Fig. 1B**) thus further amplifying the net levels of these genes in acute infection.

Module 4 for cDCs, also enriched at the acute time point, includes immune signaling genes related to HIF-1, inflammatory response, and TNFα/NFκβ pathways. Many of the innate immune response genes in this module are shared with CD14^+^ monocytes module 1---36 genes in common--such as *CD86*, SERPIN family genes, *TIMP1, SDC2, JARID2, DUSP4*, and chemokine and TNF pathway genes (**Table S3**). In addition to the responses from innate immune cell types, genes in module 3 from CD8^+^ T cells were highly expressed at the acute time point and enriched in pathways for TNFα/NFκβ as well as NOD signaling pathways. Top genes expressed in this module also include genes involved in transcriptional regulation and signaling, e.g., *JMJD1C* (KDM3C) histone H3K9 demethylase, *PMEPA1* a negative regulator of TGF-β signaling (53), and *LUZP1* (leucine zipper protein 1) an actin-stabilizing protein (54) (**Table S3**). The increased average expression of CD8^+^ T cell module 3 genes correlates with the elevated frequency of CD8+ T cells identified by CyTOF at this time point across severity groups, consistent with a robust anti-viral response mounted by expanded CD8^+^ T cells.

Concomitantly with the elevated modules were gene modules with lower expression at the acute time point, genes which may be downregulated to enhance an effective anti-viral response (**Fig. 3D**). Across severity groups at T1, we detected lower expression of module 1 of CD16^+^ monocytes characterized by mTORC1 pathway; module 1 of cDCs defining pathways of hematopoietic cells and complement; and module 1 of CD8^+^ T cells including genes in the IFNγ response (**Table S3**). Module 1 of CD16^+^ monocytes includes the anti-viral protein *SAMHD1*, actin and actin binding protein *CAP1* and *LCP1* (L-plastin) that stabilizes integrin binding for the immune synapse with cytotoxic T cells (55), and inflammatory mediator *CYBB*, Cytochrome b-245 beta chain), a component of the phagocyte NADPH oxidase complex (**Table S3**). Three of the top 10 genes in cDC module 1 are members of the S100 family and have significant roles in immune signaling and responses. *S100A8* and *S100A9* are the subunits of the calcium- and zinc-binding heterodimer calprotectin, which is a biomarker of inflammation and a potent chemotactic factor (56) (**Table S3**). S100A4 is a calcium binding hypoxia-inducible gene (**Table S3**). Reduced expression of these genes reflects the regulated program of cellular responses to viral infection and for these genes was not found to be different between asymptomatic and symptomatic cases.

### Gene modules distinguish WNV severity groups

In addition to longitudinal changes in gene expression, our analysis detected modules which distinguish severity groups. The module scores were lower in asymptomatic compared to symptomatic groups for modules 2 and 3 in non-classical (CD16^+^CD14^-^/dim) monocytes, module 2 in CD4^+^ T cells, and module 1 in pDCs. Asymptomatic participants had higher expression of module 1 in CD16^+^ monocytes and module 1 for cDCs (**Fig. 3B**). These gene changes identified in asymptomatic participants may be critical elements of immune responses relevant to reduced susceptibility to severe disease. Notably 4 of the 6 modules which distinguish severity groups were expressed at lower levels in asymptomatic donors suggesting either control of exuberant response to the virus may be a key part of asymptomatic infection or higher inflammatory responses occurred with symptomatic infection. Of the six modules showing the highest significant differences between severity groups, five modules are related to responses of innate cell types. In particular, CD16^+^ monocytes, considered anti-inflammatory in some settings (57), are the most highly represented (3 of 6 modules). Of particular interest is higher expression of genes at the acute timepoint by asymptomatic donors in CD16^+^ monocytes module 1 (also shown to be lower at acute timepoint in all groups) which includes anti-viral protein *SAMHD1*, actin and actin binding protein *LCP1* and *CAP1* as detailed above (**Table S3**). Module 1 in cDCs is higher in asymptomatic donors, including *CD62L* (SELL) previously noted to be elevated in CyTOF assays (**Fig. 2A**); 5 genes in the S100 immune signaling family; and genes predominantly included in the type I IFN response such as *OAS3*. The IFN response is a very important element of anti-viral activity and higher expression of these genes in asymptomatic participants suggests the early expression of these pathways may control infection effectively.

Further, asymptomatic donors at the acute time point show reduced expression of genes in modules 2 and 3 of CD16^+^ monocytes compared to the severe patients with encephalitis. These modules include immune signaling intermediates *FOS, FOSB,* and *JUN*, in the TNFα signaling pathway, and genes related to apoptosis. Another top gene expressed by CD16^+^ monocytes in module 3 is anti-inflammatory *ANAX1* (annexin A1) (58). Asymptomatic donors also show reduced expression of inflammatory genes in module 2 by CD4^+^ T cells which may contribute to less severe responses. Top genes in CD4^+^ T cells module 2 include *CITED2*, a transactivator which negatively regulates HIF-1, and transcription factors such as *RXRα, NFκB, STAT2* (59) as well as other transcriptional regulators including *JUN, JUNB, NUNB, FOS, FOSB, SOCS3*. The collective differences between asymptomatic and symptomatic responses revealed here through the transcriptional analysis reflect reduced expression of pro-inflammatory and apoptotic signaling genes in asymptomatic donors. The asymptomatic response is also characterized by increased expression of genes involved in cytokine responses IFNγ and TNFα/NFκΒ, IL-2/STAT5 signaling, and an increased expression of IFN anti-viral genes. These features of the asymptomatic response to acute infection would be expected to increase viral clearance and overall lower the degree of inflammation in asymptomatic donors.

### Ligand-target interactions in transcriptional responses

Having identified those differential modules of co-regulated genes associated with stage of infection or with disease severity, we employed a ligand-target inference tool, *NicheNet* (60), to infer potentially functional ligand-target pairs for the 12 modules of interest and to discover critical genes which may mediate communication between immune cells in response to WNV (**Table 3**). The 12 modules show dramatic changes over time or with disease severity and we wanted to know which ligands and pathways could be driving those changes and targeting genes in those modules. With data at a population level, we identified a particularly prominent role for IL-1β from CD14^+^ monocytes as the top ligand mediating interactions between multiple cells including an autocrine interaction from module 1 targeting CCL7 in CD14^+^ monocytes; and paracrine interactions from CD14^+^ monocytes to target IL1R1 in module 4 of cDCs as well as CD83 in module 2 of CD16^+^ monocytes. A critical role for IL-1β has been noted previously in distinguishing severity of WNV infection (26) and in control of WNV in the brain (27). This analysis also identified TNF expressed by CD16^+^ monocytes as the top ligand for autocrine signaling to target CSF1R and AHR in modules 1 and 3, respectively. The top ligand produced by CD8^+^ T cells, INFγ, was found to multiply target JUNB in both module 2 of CD4+ T cells and in module 1 of pDCs as well as IRF8 in module 6 of plasma B cells, all of which are associated with disease severity.

**Table 3.**
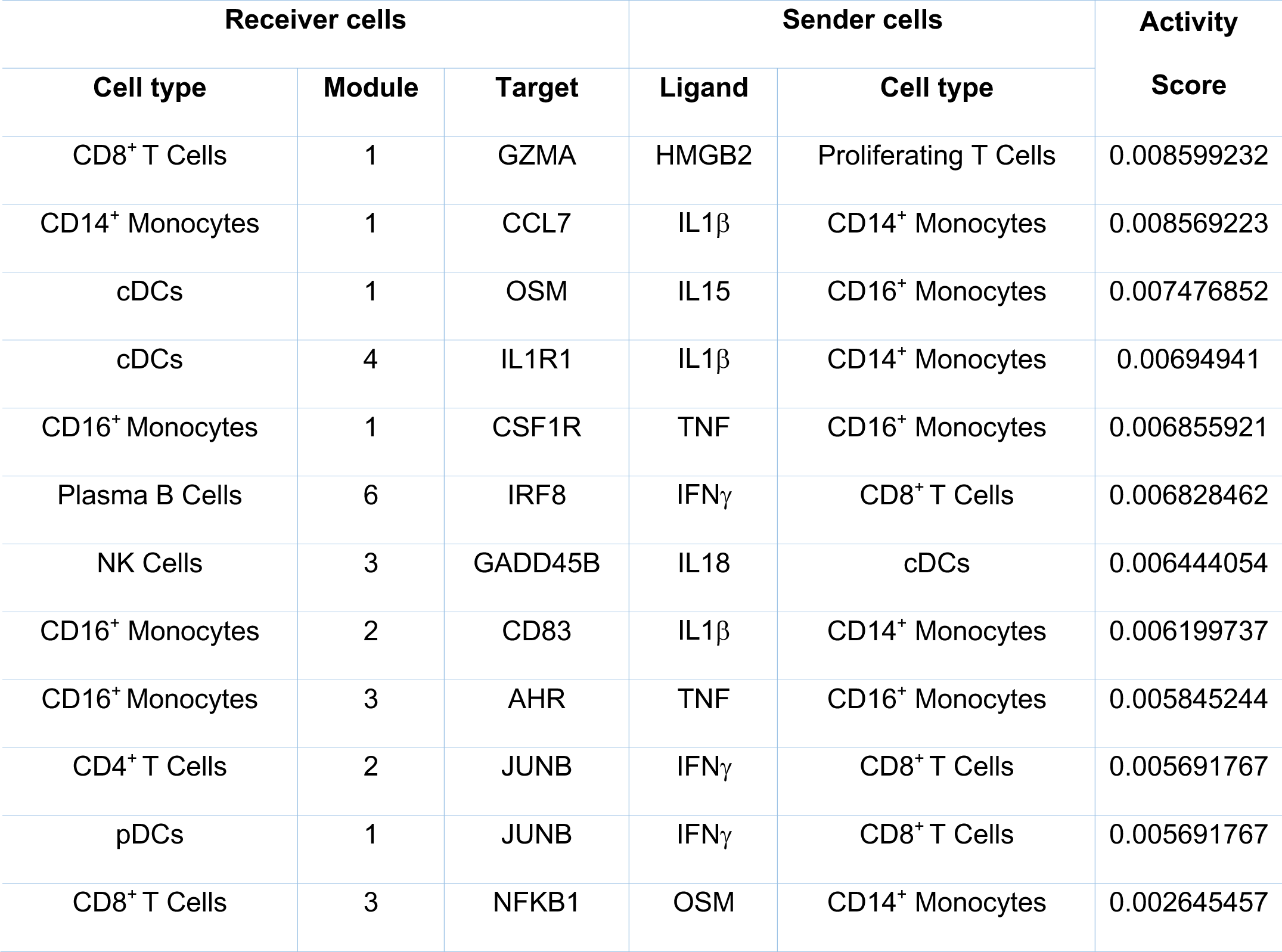
Top ligand-target pairs in cell-cell communication. The top ligand-target pairs were inferred by NicheNet analysis with the highest activity score for the 12 differential modules in Table 2. Each top ligand shows the highest expression in the ‘sender’ cell type shown.

Ligand-target interactions were also inferred in chains or inter-cellular cascades, i,e., NFκβ1 in module 3 of CD8^+^ T cells, which is enriched in NFκβ signaling as mentioned above, is a target of OSM (oncostatin M), the top ligand of CD14^+^ monocytes. OSM is an interleukin-6 family cytokine (61, 62) and module 1 of cDCs serves as a target of IL-15 from CD16^+^ monocytes. OSM was reported to be elevated during acute human anaphylaxis together with S100A8/A9 (63), two of the top three genes in module 1 of cDCs. Thus, we identified potentially important interacting factors that are highly relevant to anti-viral responses.

### Circulating soluble factors show proteins and metabolites affected by infection with WNV

To investigate the systemic milieu of the cellular immune responses and to determine which circulating factors are relevant to severity of WNV infection, we quantified proteins and metabolites in the serum of our subject cohort. We employed the MStern blotting-based proteomics platform for non-depleting profiling of the serum proteome with all high-abundance proteins remaining in the sample (64–68) We examined longitudinally collected samples to quantify differentially abundant proteins from acute infection through convalescence and to distinguish the severity groups of WNV patients (**Table S4**). When comparing participants across time points, we identified 30 significant proteins, grouped into specific clusters which showed distinct patterns of enrichment over the time course and severity of WNV infection (**Fig. 4**. p< 0.05; one-way ANOVA). In particular, we identified several proteins associated with acute phase response and inflammatory processes including the complement and coagulation pathways (69).

**Figure 4.**
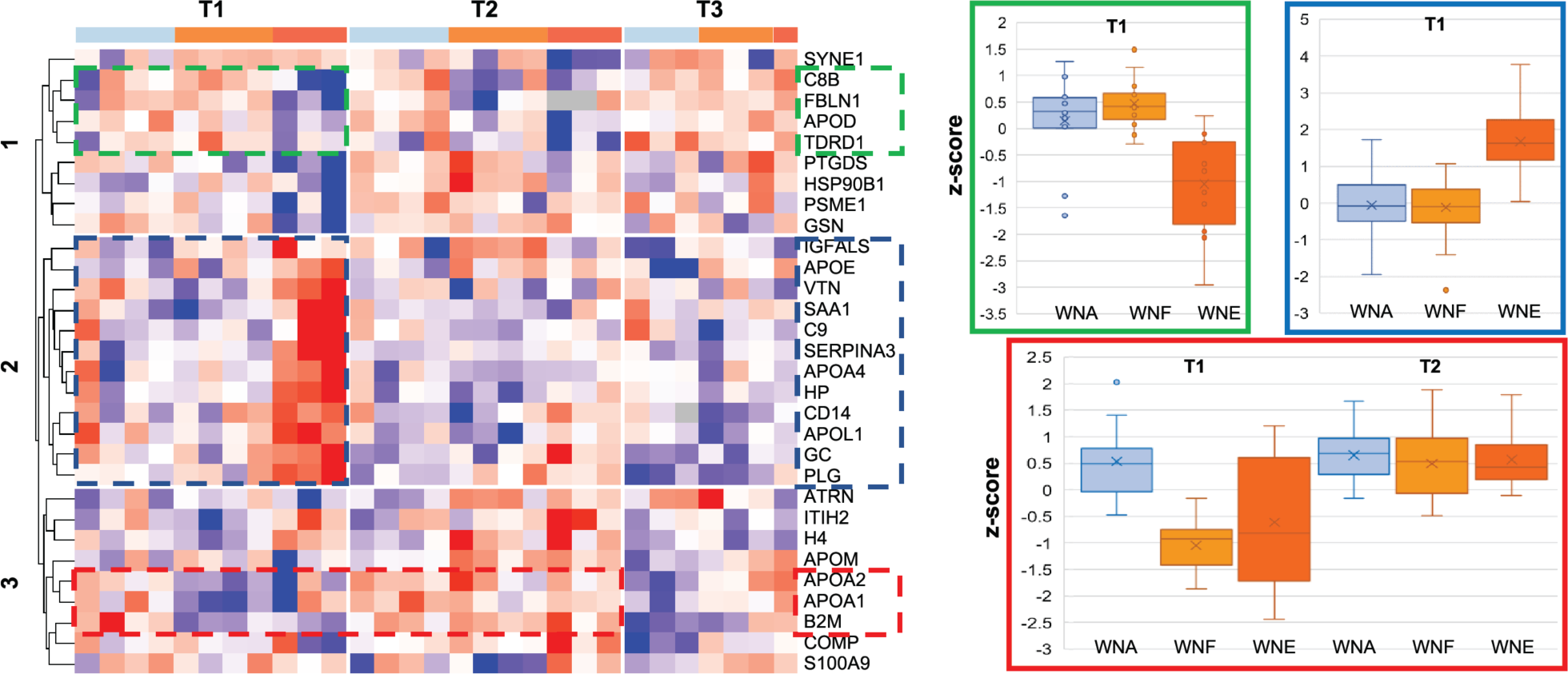
Plasma proteomics investigation in WNV infection. Serum proteins were assessed using MStern blotting followed by a highly multiplexed targeted MRM LC-MS experiment. A) Hierarchical clustering-based heat map showing the significant proteins (ANOVA; p<0.05) based on the disease severity across the different time points. Protein groups of interest are indicated by green, blue and red boxes. Box/whisker plots of the z-scores across all proteins and participants indicated by the green (B), blue (C) and red (D) boxes.

Three groups of proteins featured most prominently. Symptomatic WNF and WNE patients at the acute time point showed lower abundance of 3 proteins (B2M, APOA1 and APOA2; red box) relative to the levels observed in the asymptomatic participants which all equalize at the 3 month time point. This includes beta 2-microglobulin (B2M), a component of class I major histocompatibility complexes which is known to be elevated in viral infections (70). This pattern suggests that higher abundance of these proteins may be beneficial or predictive of an improved clinical trajectory. A group of four proteins (C8A, FBLN1, APOD, and TDRD1; green box) demonstrated elevated levels in the serum of both acutely infected asymptomatic and WNF patients but were significantly lower in WNE samples (although some heterogeneity is noted in Asymp 1 at the acute time point).

The highest number of significantly different serum proteins was a group of 12 proteins which were elevated specifically in WNE symptomatic patients at the acute time point (IGFALS, APOE, VTN, SAA1, C9, SERPINA3, APOA4, HP, CD14, APOL1, GC, and PLG; blue box). These markers include GC (vitamin D-binding protein); plasminogen (PLG), an acute phase reactant; and apolipoprotein A1 (APOL1) which responds to inflammatory cytokines and is involved in autophagy (71), and CD14, the soluble form of the monocyte surface marker which is an indicator of monocyte activation consistent with CyTOF profiles noted above in symptomatic participants. Elevated sCD14 is associated with inflammation and viral infections (72). Expression levels of these proteins decreased at Month 3 and further declined at the Year 1 time points. Our study also identified elevated levels in symptomatic participants of the serum protein IGFALS, insulin-like growth factor binding protein acid-labile subunit, which has recently been shown to facilitate ubiquitination of signaling mediators IRAK1 and TRAF6 to inhibit influenza viral replication and in addition is a sensitive biomarker of COVID-19 infection (73, 74). This proteomic profile highlights important elements in the host response to acute WNV infection and identifies differences between the severity groups, specifically reflecting higher systemic inflammation in symptomatic participants. Understanding the milieu of immune cells enriches our understanding of the response to infection and provides additional evidence of inflammatory differences distinguishing severity groups.

Interactions between metabolic and immune pathways are incompletely understood but are a critical element of host response. Circulating metabolites are a link that both reflects systemic conditions as well as modulating cell states. To identify metabolic pathways that change in response to infection with WNV, we conducted untargeted LC-MS based metabolomics on serum samples from our cohort. From a small sample set (n=11 participants, n = 22 samples), we conducted an exploratory evaluation and identified 641 metabolites across all participants. When we compared acute and Month 3 timepoints for all participants together, we detected differentially abundant metabolites with 42 increased and 35 decreased (**Fig. 5A**). In this longitudinal comparison, the biggest differences were detected in 5 glycerophopholipids of which 4 were more abundant at acute infection. These metabolites include lysophosphatidylethanolamine (LPE), lysophosphatidylcholine (LPC), and phosphatidylethanolamine (PE), which are amphipathic lipids in lipid bilayers that play a role in immune activation (**Fig. 5A**, C22:6 LPE, C22:5 LPC, C22:6 LPC. C38:6 PE). Elevations in LPE have previously been correlated with higher disease activity in rheumatoid arthritis suggesting they may be important for immune activity (75). In addition, 3 members of the alpha-linolenic acid and linoleic acid metabolism pathway were higher in acute infection and metabolites in the bile acid biosynthesis pathway showed an increased abundance at Month 3 (**Fig. 5A**). Bile acids have a regulatory role in inhibiting inflammasome activation (76), thus elevations at Month 3 may indicate a trajectory towards resolution of inflammation.

**Figure 5.**
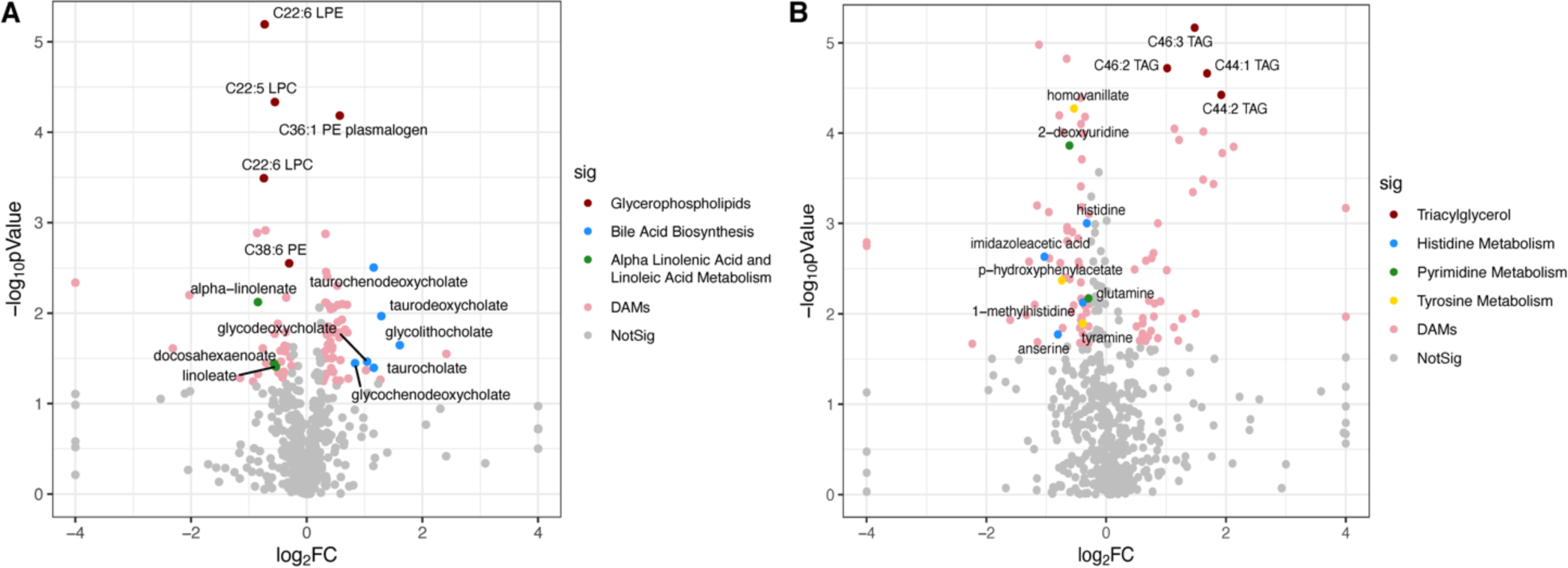
Differentially abundant metabolites in WNV infection. Serum metabolites were assessed using LC-MS, median normalized and log transformed. Differentially abundant metabolites (DAMs) are shown in volcano plots comparing (A) time point (Month 3 vs. Day 1) for all subjects at and (B) symptom severity (symptomatic n = 7 vs. asymptomatic n = 4). Colored points represent DAMs with absolute value of fold change (FC) ≥1.2, p value < 0.05 (one-way ANOVA), and FDR (Storey’s q-value) < 0.2 (A) or < 0.05 (B).

We also compared metabolic profiles between samples from asymptomatic participants (n = 8) and symptomatic patients (n =14) to highlight any changes associated with clinical severity. When we compared subject groups, we detected distinct patterns of differentially abundant metabolites with 35 increased and 70 decreased metabolites in symptomatic compared to asymptomatic participants. WNV symptomatic participants exhibited the highest increase in triacylglycerols (C46:3 TAG, C46;2 TAG, C44:1 TAG, C44:2 TAG), while asymptomatic participants showed an increase abundance of histidine, pyrimidine, and tyrosine metabolites (**Fig. 5B**). A higher abundance of histidine in asymptomatic participants is consistent with previous reports of an anti-inflammatory effect (75). These differences suggest increased regulation of metabolic pathways in asymptomatic participants during acute WNV infection which may contribute to the improved clinical trajectory. This is consistent with cell functional markers and transcriptional targets, showing elevated levels of inflammatory pathways in each omics platform in symptomatic participants is likely relevant for disease severity.

### Integrative analysis by Multi-Omics Factor Analysis (MOFA)

Each of the multi-dimensional profiling platforms (cellular, transcriptional, proteomic, and metabolomic profiling) identified distinct factors that exhibit significant associations with the time points and disease groups and which may contribute to efficacy of responses and severity of WNV infection. A key question is whether individual elements, which alone may provide limited specificity or biological insight, may be highly correlated across platforms and interact together to drive the immune response. Thus, considering features derived from the combined systems analysis may identify signatures critical to clinical outcome. For more insight into immune components, we integrated the data from all the assay platforms using a joint dimension reduction with Multi-Omics Factor Analysis (MOFA) based on features from our four different assays (CyTOF, Seq-Well, proteomics and metabolomics) (77, 78). MOFA identified 7 multi-omic factors (maximum number of factors suggested by MOFA) which explain 46.4% of the variance averaged over assays. For each factor, we performed two-way ANOVA analysis and assessed whether infection time point and disease severity explain additional variation in the MOFA factor scores across participants (**Table S5**). We focus on MOFA factor 1 which demonstrated the most significant difference across time points (p value 7.90E-07, adjusted p value 5.50E-06) and MOFA Factor 5 (p value 2.20E-02, adjusted p value 1.60E-01) with the strongest difference distinguishing between the multiple severity groups.

We found MOFA factor 1 was significantly associated with the time points. MOFA factor 1 was highest at the acute time point and decreased over time in the convalescent samples (**Fig. 6A**, p-value of 7.9E-07, adjusted p-value of 5.5E-5). The top features of MOFA factor 1 are from CyTOF and Seq-Well and include 17 cell type frequencies and 109 cell type functional marker combinations from CyTOF, 10 proteins, 15 metabolites, and 1,464 genes (**Table S6**). MOFA factor 1 expression level represents a state with elevated frequencies of activated CD8^+^ and CD4^+^ T cells as well as lower levels of NK cells, Tregs, and DCs. Increased frequency of CD8^+^ T cells and decreased frequency of NK cells at the acute time point was also identified when analyzing the cell frequencies directly (**Fig. 1**). Functional markers identified in MOFA factor 1 and elevated at the acute time point include the leukocyte adhesion marker CD62L (L-selectin) on monocytes, DCs, NKs, CD4^+^ and CD8^+^ T cells; elevated expression of IFNγ in DCs, B cells, and CD8^+^ T cells; elevated immune regulator CD279 on granulocytes and T and B cells; elevated chemokine receptors CCR4 on monocytes and CD8^+^ T cells and CCR7 on CD4^+^ and CD8^+^ T cells. MOFA factor 1 also included lower expression of checkpoint inhibitor CD152 on CD4^+^ T cells and NK cells. The most significant protein signature in MOFA factor 1 was complement and coagulation cascades (WikiPathway 3.66e-04). MOFA factor 1 includes serum proteins PROS1, involved in coagulation and clearance of apoptotic cells as a ligand for receptor tyrosine kinases TAM receptors (Tyro3, Axl, and Mer) (79) and APOA4, APOL1, which have primary roles in lipid transfer as well as immune-responsive role in in autophagy, and SERPING1 which acts to inhibits the complement cascade. Among the metabolomic profiles, MOFA factor 1 identified decreased abundance at acute infection of enriched metabolites in the steroid biosynthesis pathway and spermidine pathway, ribothymidine and methylthioadenosine, reported to have anti-apoptotic effects (80), and C34:2 PC plasmalogen, C36:1 PE plasmalogen, membrane lipids associated with immune signaling and protective against reactive oxygen species (81).

**Figure 6.**
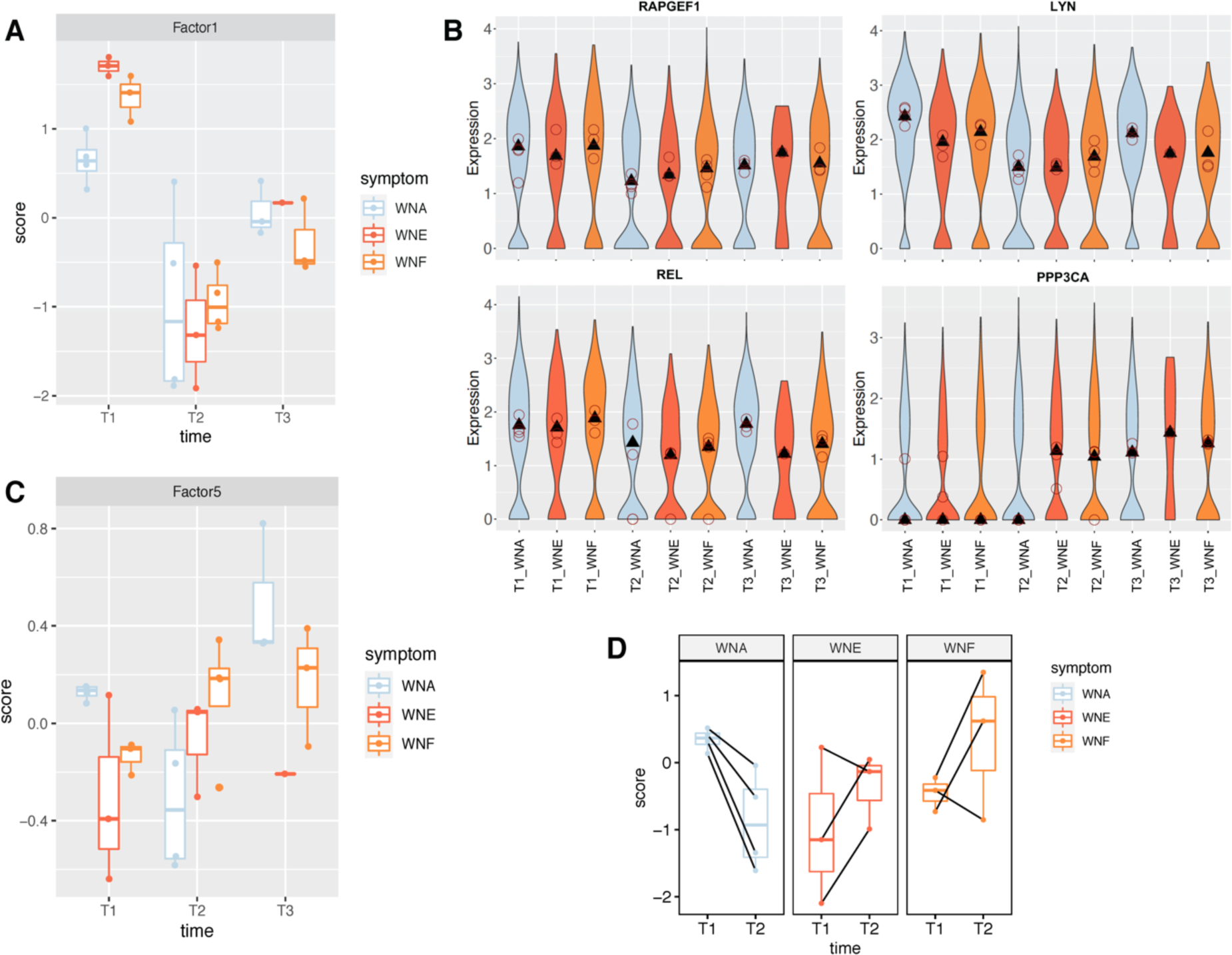
Multi-Omics Factor Analysis (MOFA). Data from all the assay platforms were integrated using a joint dimension reduction with Multi-Omics Factor Analysis (MOFA). (A.) Plots for MOFA factor 1 across time and severity groups. A two-way ANOVA test suggests a strong difference in Factor 1 (p = 8E-07) after accounting for across disease group variability. (B.) Expression of BCR signaling genes in B cell subsets across time points and severity groups. (C.) Plot across time and severity groups for MOFA factor 5. A two-way ANOVA test suggests mild differences in Factor 5 (p = 4E-3) across times and across disease groups (p = 2E-2). A Wilcoxon-rank sum test suggests that both asymptomatic and symptomatic groups show differences at the acute time point (p=8E-3). (D.) Paired-plot on changes between T1 and T2 for different participants. A Wilcoxon-rank sum test suggests that asymptomatic and symptomatic groups show differences regarding the trajectory from T1 to T2 (p = 1E-3).

In particular, B cell receptor (BCR) signaling was found to be the top enriched pathway among the 1,464 genes for MOFA factor 1 (WikiPathway, 2.01e-05), although our module-based Seq-Well analysis did not identify a B cell module. Notably, in CyTOF, the frequency of memory B cells was significantly elevated at the acute time point for all participants (**Fig. 1B**, p<0.01). These results highlight importance of B cells in early responses to infection with WNV. Since the MOFA analysis is based on global average gene expression across all cells regardless of cell types, we sought to determine whether the B cell signaling identified in MOFA factor 1 was due to changes in cell frequency or reflected activation within the B cell population. Therefore, we re-examined genes in the Seq-Well data related to BCR signaling (134 BCR signaling genes identified by two databases (WikiPathway (n=97) and KEGG (n=81) including 44 shared genes) to investigate whether there was differential gene expression within the B cell populations. We examined expression for the 22 genes in the BCR signaling pathway included in MOFA factor 1, comparing B cells in the 18 pairs of severity states and timepoints. Strikingly, we found differential expression for 21 of 22 genes in at least one condition (p-value < 0.01, Wilcoxon rank sum test) and highly significant differential expression for 16 of 22 (p-value < 0.001). Particularly interesting are 3 genes elevated in the acute samples that are significant across time and severity groups (**Fig. 6B**): *RAPGEF1*, a guanine nucleotide exchange factor involved in signal transmission and lineage specific differentiation (82); *LYN* and *REL*, activating src family tyrosine kinases. In contrast, *PPP3CA*, a calcineurin protein which activates T cells, showed higher expression during convalescence particularly in the symptomatic cases reflecting their higher inflammatory status. For a subset of samples with sufficient cell numbers to support a separate analysis, we identified significant differences within plasma B cells specifically and similarly found lower expression at the acute time point for expression of *HCLS1* and *PPP3CA (*p-value < 0.01) consistent with expected kinetics of plasma cells in early response and the differential levels for B cell subsets. Thus, our MOFA factor 1 highlights the importance of BCR signaling at the acute response and, when combined with elevated cell frequency, identifies B cell signaling as a key element in response to WNV infection. This finding, while consistent with our transcriptomic data, was not evident from the Seq-Well module analysis alone.

We found a significant severity-related effect in MOFA factor 5 (**Fig. 6C**, factor 5, ANOVA p-value of 2.3E-2, adjusted p value 0.16) with asymptomatic participants having higher expression of MOFA factor 5. In addition, MOFA factor 5 also showed a difference across time points (3.90E-3, adjusted p value 2.30E-02). Further, the trajectory of paired MOFA factor 5 scores for each individual, calculated as the difference between levels at acute and 3 month time points, were significantly different between severity groups, with the symptomatic group increasing compared to asymptomatic groups (**Fig. 6D**. p-value of 1.7E-3, adjusted p-value of 1.2E-2; Wilcoxon rank-sum test). The top features of MOFA factor 5 are balanced between CyTOF, Seq-well, and proteomics and include 2 cell type frequencies and 4 functional marker combinations from CyTOF, 14 proteins, 14 metabolites, and 744 genes (**Table S7**). Asymptomatic participants had higher expression of MOFA factor 5 at the acute time point which represents an immune state of lower frequency of total B cells, and lower pro-inflammatory signaling pathways including reduced TGFβ in CD8^+^ T cells, pSTAT1 in mDCs, and pSTAT5 in CD4^+^ T cells. Serum proteins with elevated levels in MOFA factor 5 include components of the complement and coagulation cascades, e.g., complement protein C1S, Alpha2-HS glycoprotein (AHSG), involved in endocytosis and previously associated with severity in a murine model of WNV (83). MOFA factor 5 includes reduced levels of complement protein C1QB and APOC3 which has a primary role in triglyceride transport but may also be relevant in inflammation as a competitor of anti-inflammatory apoE (84, 85). Metabolites identified in MOFA factor 5 were propanoate metabolism which is important in mitochondrial energy metabolism (86), and alpha-hydroxybutyrate, an alternate energy source and regulator of cellular signaling (87). MOFA factor 5 transcriptional features were highly enriched in mRNA processing (WikiPathway 1.8e-05). Taken together, the key cellular immune responses and metabolic pathways of MOFA factor 5 identify a constellation of critical immune elements which contribute to successful responses. Moreover, the significance of opposite trajectories of MOFA factor 5 between severity groups from acute to Month 3 time points suggests that the different immune profiles evident between clinical groups during acute infection may drive different trajectories for clinical outcome.

### Discussion

WNV infection can be severe or fatal for only a minority of individuals. In this study we for the first time employed a longitudinal Multi-Omic approach beginning with acute infection to define early cellular and molecular signatures corresponding to severe WNV infection. Insights we gained from defining effective immune responses may inform future investigation of specific cellular mechanisms and novel preventative and therapeutic approaches to mitigate severe WNV in susceptible people. We took advantage of an unusual population of acutely infected but asymptomatic individuals and interrogated immune responses to identify individual variations that may contribute to infection severity. We applied powerful single cell approaches to interrogate immune responses at acute infection and through convalescence. Following multiple measures over time from the same study participants reduces variation in our assays and supports development of the trajectory of infection response. Through an integrated analysis comparing multiple immune components in asymptomatic and symptomatic cases, we defined elements characterizing different clinical outcomes.

A striking finding from our study is the similarity of responses in the study participants. Across severity groups we detected significant alterations in cell frequencies and functions, transcriptional, proteomic, and metabolomic pathways, reflecting that WNV is a potent virus which evokes strong immune responses. We have previously quantified antibody responses to WNV proteins (envelope and non-structural proteins 1 and 5) and assessed humoral response to WNV on a single-cell and repertoire level and shown no significant differences between groups whether they experienced asymptomatic or severe infection (31, 88). Changes in cell frequencies — such as elevated innate immune PMN and monocytes at acute infection, and the expansion of effector CD4^+^ (Temra) cells at Month 3 and Year 1 — were similar across severity groups. We showed differences, however, in cellular functional responses with relatively elevated levels of inflammatory cytokines (TNF, IL-8, IFNγ) and signaling mediators (phospho STAT3, phospho STAT5) in the group with severe WNV infection. In transcriptional responses, similarly, all participants demonstrated elevated transcription of cDC, CD8^+^ T cell, and CD14^+^ monocyte gene signatures at acute infection which declined by Month 3 and Year 1. We also identified particularly notable differences in gene modules in non-classical CD16^+^ monocytes of asymptomatic participants, a subset specialized for wound healing and anti-viral responses (89). Through a ligand-target analysis of cell-type and state based on average signals of individuals, we identified markers of interest enriched during infection and in different disease trajectories to ground generated gene-sets in known cellular interactions.

We broadened our examination of immunity by profiling serum proteins and metabolites, identifying significant differences longitudinally in all study participants as well as specific components that distinguish severity groups. In particular, these analyses while limited by sample size highlighted elevated serum levels in more severely ill WNV participants of acute phase reactants and inflammatory processes including the complement and coagulation pathways (69), and that serum levels of some proteins remained elevated over 3 months, consistent with other clinical observations (90). Notably, insulin like growth factor binding protein acid-labile subunit (IGFALS) was also higher in more severely ill WNV study participants which may reflect its role inhibiting viral infection (73, 74). Asymptomatic participants had elevated levels of complement and coagulation, and elevated levels of the metabolic pathways enhancing energy production. These elevated proteins are consistent with acute serum proteome responses in Lyme disease, another arthropod-borne infection, although microbiologically distinct and transmitted by a different vector, suggesting similar immune activation (68). With respect to serum metabolites, all groups at acute infection showed elevated levels of glycerophospholipid, which is associated with immune activation and has recently been associated with inflammatory responses to influenza vaccination Further, we detected an increased abundance of anti-inflammatory histidine metabolites in asymptomatic individuals.

With this data resource as input, we applied Multi-Omics Factor Analysis (MOFA) to generate salient hybrid features derived from the combined systems analyses. With the integrated data from all the assay platforms, several distinct findings emerged. As noted in each individual assay platform, there were many elements of response to WNV infection which were shared by all severity groups. Despite engagement of these responses, individuals show highly divergent clinical outcomes.

The coordinated elements identified via integrated MOFA multi-omics modeling provide mechanistic insight and may identify signatures critical to clinical outcome. Specifically, MOFA factor 1 identified significant elements associated with the time course trajectory of infection with a focus on B cells that correlates with cell frequency data and is consistent with higher gene expression levels at the acute time point. This response may be critical, with early activity of B cells having an important protective role in infections in both asymptomatic and symptomatic subjects. Studies in the mouse model showed that B cells were essential for resistance to WNV infection We have previously shown that antibody levels examined in convalescence were not significantly different between asymptomatic and severely ill participants and did not predict clinical severity (31, 88) and we similarly found strong anti-WNV E responses by immunoblot from all participants in the current study. Thus the importance of the B cell transcriptional signatures as identified in the MOFA analysis may be related to other B cell roles and interactions besides secretion of antibody.

Through our analysis we determined that effective immune responses in asymptomatic donors include an activated immune status involving cell types and pathways that initially controlled pro-inflammatory innate immune cell activation and elevated numbers of Tregs noted in MOFA factor 1. This observation is consistent with previous measures of Tregs in infection with WNV in studies restricted to symptomatic participants (28). Our MOFA analysis identifies an immune state combining multiple immune components and suggests that less severe WNV infection is associated with a combined immune state of reduced proinflammatory activity and elevated levels of B2m, ApoA1, Apo A2 proteins. While very few individual features independently can distinguish disease severity groups, especially in a small cohort, with the combined activity of many potentially small contributions of several immune components, MOFA identifies an immune state whose joint effect is more significant for separating the disease cohorts.

This observation could not be addressed in previous studies examining only severely infected patients or from profiles of convalescent cohorts enrolled after recovery from infection (28, 29, 93). Although stratified by disease severity, they were evaluated after recovery from acute illness when differences in initiation of B cell responses were not evident (31, 88). The mechanisms identified here in profiles of asymptomatic study participants may be useful for understanding mechanisms of pathogenesis for other viral infections as well, particularly for related flaviviruses such as Zika and dengue viruses (37, 94, 95).

Our study cohort is unique but limited by enrollment size which is dependent on variations in infection frequency at our recruitment site (Houston, TX) and was conducted during disruptive seasons which included Hurricane Harvey as well as the COVID-19 pandemic. Focusing only on WNV-exposed individuals allows us to assess the determinants of different responses to the virus. Healthy uninfected individuals, while easier to recruit, are not appropriate controls for this study, as their anti-WNV efficiency is undefined prior to infection. Although WNV infections are typically asymptomatic (31), it is unknown how individual uninfected controls would respond to infection, making such a group incompatible with our goal of defining molecular charactertistics that differentiate asymptomatic and symptomatic infection responses. Our inclusion of a convalescent timepoint at Year 1 served as a proxy to healthy conditions. Beyond the distinct immune elements identified here, additional factors and mechanistic conclusions may be revealed in a study using a larger cohort and where participant sex could be considered, mild fever and severe neurologic subjects could be distinguished, and detailed medication history could be collected from asymptomatic participants. In addition, WNV is a vector mediated infection, and our study does not specifically examine the impact of mosquito salivary components. The host immune response to the vector has recently been highlighted as contributing to resistance in some tick-borne infections (96) and shows promising protection with anti-mosquito vaccine or interventions directed at viral proteins such as non-structural (NS)-1 (97).

Our approach, including a longitudinal stratified profiling plan and MOFA analysis, offers a data resource of acute human viral infection and a model for multifactorial immune inquiry. Through this in-depth profile, we uncovered distinctions between patients who have very dramatic differences in clinical outcome. Medicine is developing a rich understanding of cellular immune pathways and how they may be leveraged for clinical advantage. With an additional understanding of the proteomic and metabolomic datasets, the profiles provide potentially actionable biomarkers for vulnerable individuals. Beyond prognosis, defining measurable differences in immune response between disease trajectories can inform targeted approaches to prevent and/or treat WNV infection, including selective immunomodulation.

## METHODS

### Enrollment of Human Study Participants and Ethics Statement

This study was conducted in accordance with guidelines approved by the Human Investigations Committees of Baylor College of Medicine and Yale School of Medicine. Participants were untreated and enrolled and samples collected with written informed consent approved annually by each institution. All eligible patients infected with WNV during the enrollment period were recruited at Baylor within seven days of symptomatic onset of mild disease (fever only, WNF) or severe disease (invasive neurological illness such as encephalitis or meningitis, WNE) as defined by Center for Disease Control (CDC)-defined clinical criteria of neurologic involvement with infection confirmed by RT-PCR (98). All eligible viremic asymptomatic donors during the enrollment period were identified at Gulf Coast Regional Blood Center donation sites (Houston, TX) through routine screening of blood by rapid nucleic acid test. As the threshold of WNV viremia detectable by the screening test indicates viral exposure occurred within 7 days (99) and asymptomatic participants (WNA) were enrolled 2-4 days following screening, duration of infection was comparable in both groups of participants. All participants regardless of symptoms showed antibody recognition of WNV envelope protein by immunoblot (31) to support comparison of immune responses directly. Asymptomatic donors had no acute illness and took no antibiotics or nonsteroidal anti-inflammatory drugs at the time of sampling. All enrolled participants were Caucasian. Medical history was obtained at the time of enrollment. As this study specifically focuses on molecular characterization of symptomatic and asymptomatic infection responses, we did not include healthy uninfected individuals as their anti-WNV immune efficiency is undefined prior to infection.

### Sample collection and cell isolation

Blood samples were collected from WNV infected participants over a total period of 19 months taking advatgae of stable characteristics in individuals (32–34) with collection from each individual at enrollment (T1) and at 3-month (T2) and 1-year (T3) convalescent time points when participants had resolved their illnesses. Blood draws at Baylor were collected in BD Vacutainer tubes, sodium heparin tubes, or BD Vacutainer CPT Mononuclear Cell Preparation Tubes with Sodium Citrate as an anti-coagulant (BD Biosciences, #362761). Whole blood without anti-coagulant was collected for preparation of serum that was stored at -80°C until assay. Whole blood samples from sodium heparin tubes were processed in SMART tubes containing Proteomic Stabilizer PROT1 fixation buffer (SMART Tube, Inc., Las Vegas, NV), frozen on site according to the manufacturer’s instructions, and shipped frozen. Blood samples in CPT tubes were processed within 2 hours of collection according to the manufacturer’s instructions and shipped overnight for processing the next day (26). Fresh peripheral blood mononuclear cells (PBMCs) were collected from CPT tubes, washed twice with 90% RPMI + 10% FBS, and processed fresh for transcriptomics or cryopreserved in 10% DMSO, 90% FBS in liquid nitrogen at a concentration of 5 x 10^6^cells/mL.

### Cell labeling and mass cytometry acquisition

Cryopreserved whole blood samples were harvested from SMART tubes according to the manufacturer’s instructions for mass cytometry (CyTOF). Samples were labeled with a uniform antibody cocktail as described previously (37, 100). Metal-conjugated antibodies (**Table S1**) for labeling cells were purchased from Fluidigm (Fluidigm Inc, South San Francisco, CA), or carrier-free antibodies were conjugated in house using MaxPar X8 labeling kits according to manufacturer’s instructions (Fluidigm). Briefly, cells from SMART tubes were thawed in batches and washed with Maxpar® Cell Staining Buffer (Fluidigm), incubated with surface labeling antibodies for 30 min on ice, permeabilized (BD FACS Perm II) for labeling with intracellular antibodies for 45 min on ice. Cells were suspended overnight in iridium interchelator (125 nM; Fluidigm) in 2% paraformaldehyde in PBS and washed 1X in PBS and 2X in H_2_O immediately before acquisition. A spike-in reference sample of prelabeled Ta181-PBMC (Biolegend, Cat#427201, San Diego, CA) was added to each subject sample as a control to minimize the effects of batch variation (101). Samples were run on Helios 2 CyTOF instrument at a flow rate of 30 µL/min and a minimum of 10,000 events was collected in the presence of EQ Calibration beads (Fluidigm) for normalization. An average of 17,870 ± 8,013 cells (mean ± s.d.) from each sample were acquired and analyzed by CyTOF.

### CyTOF data processing and analysis

CyTOF FCS files were bead normalized using CyTOF® Software v7.0 (102). Manual gating was performed on the Cytobank platform by exclusion of debris (Iridium^low^, DNA^low^), multi-cell events (Iridium^hi^, DNA^hi^), and doublets using the default data transformation of a hyperbolic arcsine (37, 100). Differences in frequencies of cell subsets between cohorts and time points were assessed by one-way ANOVA for multiple comparisons (Prism; GraphPad Software, San Diego, CA). Mean expression level of markers from spike-in reference cell data was assessed by PCA analysis and no significant difference was found between time points and severity groups thus no batch corrections were applied (**Fig. S1**; two-way ANOVA (Tukey’s multiple comparisons test). For the functional status reflected by cell lineage-functional marker combinations, given the limited sample size, an exploratory statistical analysis was conducted using a permutation sampling approach. Mean channel intensity (arcsine) of functional markers was compared for the 580 cell type/marker combinations resulting in 274 combinations above a threshold of 20% expression for that marker. The fold change of mean responses for each cell type and marker was computed for symptomatic patients and asymptomatic cases. A distribution of permuted fold changes was determined for 1000 iterations of group assignment without replacement. Associated p-values were computed from the rank of the observed fold change in the empirically constructed distribution using MatLab 2021a.

### Single cell transcriptional assays

PBMCs were washed twice in RPMI/10% FBS and residual red blood cells were lysed with 1x RBC Lysis Buffer (eBioscience, Invitrogen, #00433357) according to the manufacturer’s instructions. Cells were processed for Seq-Well S^3 single-cell RNA-seq with second strand synthesis (47, 103). Cells were loaded on Seq-Well arrays at a density of 15,000 cells. Paired-end sequencing libraries were prepared from amplified cDNA with the Nextera XT DNA sample Prep Kit (Illumina, #FC-131) according to the manufacturer’s instructions. Indexed cDNA libraries were pooled at equimolar ratios and sequenced on an Illumina NovaSeq 6000 system. Final libraries were sequenced with a 21-50-50 read structure. Libraries prepared from fresh or cryopreserved cell libraries were consistent and data from both methods are presented.

### scRNA-seq raw data processing

BCL files were demultiplexed by nine base-pair assay barcodes (one 3’ and one 5’ barcode per assay) and converted to FASTQs using bcl2fastq. Paired end reads were aligned to the hg19 genome using STAR 2.7 alignment software (104) via the Broad Institute dropseq_workflow pipeline (https://github.com/broadinstitute/Drop-seq). To generate count matrices, reads were sorted on the cell level (where a 12bp barcode corresponds to an individual cell/mRNA-capture bead) and then sorted on the transcript level (where an 8bp barcode corresponds to a unique transcript), yielding gene-by-cell count matrices for each set of paired FASTQs. Low quality cells were filtered out if they had fewer than 500 total transcripts, fewer than 300 unique mapped genes, or more than 25% mitochondrial reads. The count matrices were log-normalized (with a scale-factor of 10,000) before further data-cleaning and downstream analysis.

### scRNA-seq cell type identification

Cell types were identified by unsupervised clustering and differential expression analysis together with canonical lineage markers in an iterative manner using the Seurat R package (48). First, 3,000 highly variable genes were selected from the merged dataset as detailed previously (105). The distribution of expression for each gene was then centered around zero, using the Seurat package ScaleData function. These standardized expression values served as inputs for principal component analysis (PCA). The first n principal components (PCs) are selected to capture 75% of the variance and a k parameter for clustering was chosen as well (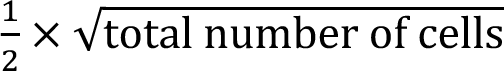). Using the top n=13 PCs and the chosen k parameter (k = 130), a k-nearest neighbors’ algorithm was applied using the Seurat package. A Shared Nearest Neighbor (SNN) graph was constructed, and clusters were established by the Seurat function, FindClusters (resolution parameter = 0.5). Clusters were visualized with the dimension reduction technique, Uniform Manifold Approximation and Projection (UMAP) (106), with a Wilcoxon rank-sum test to determine positive gene markers for each cluster compared to all other clusters (log-fold change > 0.25 and an adjusted p-value < 0.05). Cell type assignments for each cluster were assigned based on differentially expressed genes and the relative expression of 36 canonical lineage markers (**Fig. S4**). Every cell type was re-clustered individually using the same method. At this re-clustered level, doublets (determined by co-expression of lineage markers) could be identified and filtered out and more refined cell types could be defined (e.g., identifying regulatory T cells within CD4+ T cells). We first identified a total of 31 cell types (**Fig. S4**) and then focused on 11 main PBMC cell types which largely correspond to our CyTOF data (**Fig. 3A**): B cells, CD14^+^ monocytes, CD16^+^ monocytes, CD4^+^ T cells, CD8+ T cells, cDCs, pDCs, NK cells, plasma B cells, proliferating T cells, and T-regs.

### scRNA-seq differential gene module discovery using WGCNA

We used the Weighted Gene Co-expression Network Analysis (WGCNA) framework (49, 107) to discover *significant gene modules* for each cell type as detailed previously (108). Gene co-expression was considered across all cells at all timepoints for each cell type and also within each time point for each cell type. As scRNA-seq data are typically very noisy with large numbers of dropout events, we did not assume that gene co-expression networks should follow a scale-free topology to determine the soft-power threshold. We instead used a fixed soft-power threshold of 5, which gives more consistency in terms of mean connectivity across different networks from our data. *Significant modules* were identified by comparing the observed module gene dissimilarity (from the topological overlap matrix) to 10,000 randomly generated gene modules using the Wilcoxon rank sum test with FDR < 0.05. Overall module expression was assessed by scores calculated by *AddModuleScore* in Seurat. We identified *differential modules*, among those *significant modules*, for 18 pairs of different disease severity states and time points (i.e., all 3 severity pairs x 3 timepoints and all 3 timepoint pairs x 3 severity states) by performing the Wilcoxon rank sum test and the Kolmogorov-Smirnov (K-S) test for differential distributions of module scores in each condition pair in question (restricted to conditions with at least 10 cells). In addition, we examined distributions of fold changes between all condition pairs for each module. P-values from the Wilcoxon and K-S tests were systematically varied from 0.001 to 0.1 (from 0.001 to 0.009 by 0.001 and from 0.01 to 0.1 by 0.01) without adjustment for multiple testing at this stage. This lenient filtering method favors identification of potentially relevant modules for subsequent analysis and biological interpretations. In this work, we deem *differential modules* to be those with the mean p-value of the two tests < 0.01 and the fold change of median module scores > 1.3 in any of the 18 condition pairs (**Table S2, S3**). Genes in each differential module were ranked by Pearson correlation coefficients between gene expression values and module scores. For functional gene set enrichment analysis of modules, we used *Enrichr* (109).

### scRNA-seq intercellular communication predictions with NicheNet

To identify upstream ligands that affect expression patterns of the WGCNA-based differential gene modules above, we employed the NicheNet method that integrates prior knowledge of ligand-receptor interactions, signaling networks, and gene regulatory networks compiled from public data sources and validated using more than 100 microarray studies (60). We considered the 11 cell types for both receiver and sender cells. Each differential gene module was considered as a gene group of interest in the receiver cells, i.e., a group of target genes for ligands in the sender cells for each cell type. NicheNet identifies “active” ligands that significantly changes expression of module genes by calculating ligand activity scores for ligand-target pairs. We note that active ligands are considered to be ligand genes that are expressed in >10% of the sender cells in question. We analyzed the top 20 active ligands for each differential gene module for each cell type.

### Proteomics

Samples were processed according to the MStern blotting protocol developed in the Steen lab (65, 67). In brief, 1 µL of serum (∼50 µg of proteins) was mixed in 100 µL of urea buffer (8M urea in 50 mM ammonium bicarbonate) without depleting any high abundance proteins.

Following reduction (50 mM DTT) and alkylation (250 mM iodoacetamide) of the cysteine residues, 10-15 µg of proteins were loaded onto a 96-well plate with an activated and primed hydrophobic polyvinylidene fluoride (PVDF) membrane at the bottom (Multiscreen HTS 0.45 μm Hydrophobic High Protein Binding Membrane 96-well Filtration plate (Product No. MSIPS4510; Millipore-Sigma) (65, 67). Trypsinization of the proteins adsorbed to the membrane was achieved by incubation with the trypsin for 2h at 37°C. Resulting tryptic peptides were eluted off the membrane with 40% acetonitrile (ACN)/0.1% formic acid (FA). The resulting peptide mixtures were subsequently desalted using a 96-well MACROSPIN C18 plate (TARGA, The NestGroup Inc.). The equivalent of 1 μg of peptide was analyzed using a microLC system (Nexera Mikros, Shimadzu, Columbia, MD) equipped with SHIM-Pack MC-C18 column (5 cm length, 300 μm internal diameter, 1.9 μm particle size, Shimadzu) coupled online to an LCMS 8060 triple quadrupole mass spectrometer (Shimadzu) operated in targeted ‘multi reaction monitoring’ (MRM) mode using a 10.3 min gradient (5 to 40% buffer B: acetonitrile with 0.1% formic acid).

The final scheduled MRM method monitored 1,887 transition pairs associated with 629 unique tryptic peptides and 306 proteins (i.e., 1-3 peptides per protein; 3 transition pairs per peptide). A detailed list of transitions is given in **Table S4**. Using Skyline software (v20.2.1), the average peptide intensities were calculated for each protein, and the protein abundance in all samples were exported into a .csv format file for further analysis in PRISM/GraphPad and/or R.

### Metabolomics

Serum metabolomic profiles of participant samples were measured using a combination of four LC–MS methods that measure complementary metabolite classes, as previously described (91, 110): two methods that measure polar metabolites, a method that measures metabolites of intermediate polarity (e.g., fatty acids and bile acids), and a lipid profiling method. For the analysis queue in each method, participants were randomized and longitudinal samples from each participant were randomized and analyzed as a group. A pooled QC sample for the LC-MS methods was created by combining aliquots from each sample. Additionally, pairs of pooled reference samples were inserted into the queue at intervals of approximately 20 samples for QC and data standardization. Samples were prepared for each method using extraction procedures that are matched for use with the chromatography conditions. Data was acquired using LC-MS systems consisting of Nexera X2 U-HPLC systems (Shimadzu Scientific Instruments; Marlborough, MA) coupled to Q Exactive/Exactive Plus orbitrap mass spectrometers (Thermo Fisher Scientific; Waltham, MA).

HILIC-pos (positive ion mode MS analyses of polar metabolites), HILIC-neg (negative ion mode MS analysis of polar metabolites), C18-neg (negative ion mode analysis of metabolites of intermediate polarity (e.g. bile acids and free fatty acids), and C8-pos (positive ion mode analysis of polar and nonpolar lipids) LC-MS methods were carried out as previously described (91).

#### Metabolomics Data processing

Raw data were processed using TraceFinder 3.3 (Thermo Fisher Scientific; Waltham, MA) and Progenesis QI (Nonlinear Dynamics; Newcastle upon Tyne, UK). Peak identification was conducted by matching measured retention times (RT) and mass to charge ratios (m/z) to mixtures of reference metabolites analyzed in each batch. We matched unknown features to an internal database of >2600 compounds that have been characterized using methods as previously described (91, 110). This library contains compounds that have been confirmed by matching their RT, m/z and MS/MS fragmentation patterns in multiple human biofluids using authentic reference standards. To annotate unknowns based on this library, we used in-house alignment scripts to adjust the RT and m/z and match study unknowns to the compound library. No MS/MS was generated for this study. Temporal drift was monitored and normalized with the intensities of features measured in one of the doubly injected QC pooled reference samples using a nearest neighbor approach, where sample intensities in each pool are used to scale their closest samples. To determine the analytical precision for each measured metabolite, we computed coefficients of variation (CV) for annotated and unknown features using the remaining QC pools not used for scaling temporal drifts. Average CV values per method for annotated compounds was 7-11%, which is within the historical analytical. Finally, principal component analyses were generated and scores plots used to identify potential outlying samples.

For computational and statistical analysis of metabolomics data, relative metabolite intensities were pre-processed, normalized, and log-transformed. Undetected metabolites were imputed with half of the minimum value of that metabolite. Metabolites were median normalized within samples, and intensities were scaled by multiplying by 10^6^ and log transformed to stabilize variance. We used the R package *lme4* to fit a linear mixed-effects model with age, gender, symptoms, and time as fixed effects and participants as random effects (111). One-way ANOVA was used to test full vs. reduced model based on F-test to evaluate p-values and corrected using multiple comparisons (112) to calculate false discovery rate (FDR). Full model: ∼ Time + Symptoms + Age + Gender + l(1|Subject). Reduced model in the longitudinal group was ∼ Symptoms + Age + Gender + (1|Subject) and in the symptoms group was ∼ Time + Age + Gender + (1|Subject). Differentially abundant metabolites (DAM) were defined by thresholds of *p* value < 0.05 and |FC| ≥ 1.2 and FDR) estimation was performed from a collection of *p*-values. We used the human metabolome database (HMDB) (113) to classify metabolites into subclass or pathway.

### Latent Factor Models for multi-omics data-integration

Data from all the assay platforms were integrated using a joint dimension reduction with multi-omics factor analysis (MOFA2) to construct multi-omics modules (77, 78). MOFA factors were derived to explain variabilities across samples for CyTOF (34 cell type proportions), per-cell type functional marker expression levels (580 combinations), proteomics (265 proteins), metabolomics (578 metabolites), Seq-Well cell-type-specific average gene expression levels (7610) and average cell-type-specific module expression levels from Seq-Well (47). The omics data were aligned for the 28 available samples. Each factor was linked to features from multiple omics, which allowed us to interpret their biological functions using public resources for different omics: Enrichr (109) for transcriptomics and proteomics (using two databases of KEGG_2021_Human and WikiPathway_2021_Human) and MetaboAnalyst (114) for metabolomics (using Over Representation Analysis by the hypergeometric test with default parameters). We assessed 7 MOFA factors and tested whether their factor scores were associated with different time points or disease states across samples. We performed an Analysis of Variance (ANOVA) test, and compared variance explained for each factor using time points, disease groups, or both. We also considered a Wilcoxon rank-sum test comparing the paired changes from acute phase to month 3 for different disease groups. These changes are calculated using the paired observation for each participant, i.e., the change Δ_*i*_ for participant *i* is calculated as Δ_*i*_ = *X*_*i*2_ − *X*_*i*1_, where *X*_*i*1_ is the factor score at the acute phase and *X*_*i*2_ is the factor score at month 3 for this participant. We then tested whether the distribution of Δ_*i*_ is the same for the asymptomatic group and the symptomatic group. Investigation of Δ_*i*_ can reveal dynamics in immune profiles during early infections. In addition, when interpreting MOFA features, we focused on a set of top features that meet either of the following two criteria: (1) factor-feature Pearson correlation p-value < 0.01 and (2) MOFA absolute weights > Q3 + 1.5*IQR, where Q3 is the third quartile and IQR is the interquartile range. Adjusted p-values remain significant after multiple testing correction.

### Data deposition

The data supporting this study is available at ImmPort (immport.org) under study accession SDY 1396 and SDY 1397.

### Code availability

The custom code used for Seq-Well differential modules and MOFA analyses is in the Supplemental Software available using GNU General Public License version 3.

## Supporting information

Supplemental Software

## Acknowledgements

This work was supported in part by awards from the NIH/NIAID Human Immunology Project Consortium (HIPC) grant (AI089992 to RRM, DAH, SHK, JCL, AKS; AI118608 to BF, KKS, HS, OL), the Koch Institute Support (core) NIH Grant P30-CA14051 from the National Cancer Institute, and the Claude D. Pepper Older Americans Independence Center from the NIH/NIA (P30AG021342) to BvW. The *Precision Vaccines Program* is supported in part via the Department of Pediatrics at Boston Children’s Hospital. The authors gratefully acknowledge the valuable contributions of the HIPC group and Yale School of Medicine colleagues and core facilities (Yale CyTOF core and Yale Center for Genome Analysis), Ms. Jeanne Ruff and Ms. Allison Lino for assistance with the Houston West Nile Cohort, Dr. Susan N. Rossman and members of the Gulf Coast Regional Blood Bank for identifying asymptomatic participants, and all the study participants. The funders had no role in study design, data collection and analysis, decision to publish, or preparation of the manuscript.

## Author Contributions

Conceived and planned the study DAH, SHK, RRM,KOM. . Performed experiments YZ, IF, XW, HK, CHC, BF, KKS. Analyzed and visualized data HJL, YZ, IF, SM, XW, BVW, CBC, HK, CHC, BF, KKS, OL, HS, AKS, LG, SHK, RRM. All authors contributed to writing and reviewing the manuscript. RRM coordinated the contribution of all authors.

## Supplemental Figures

**Figure S1.**
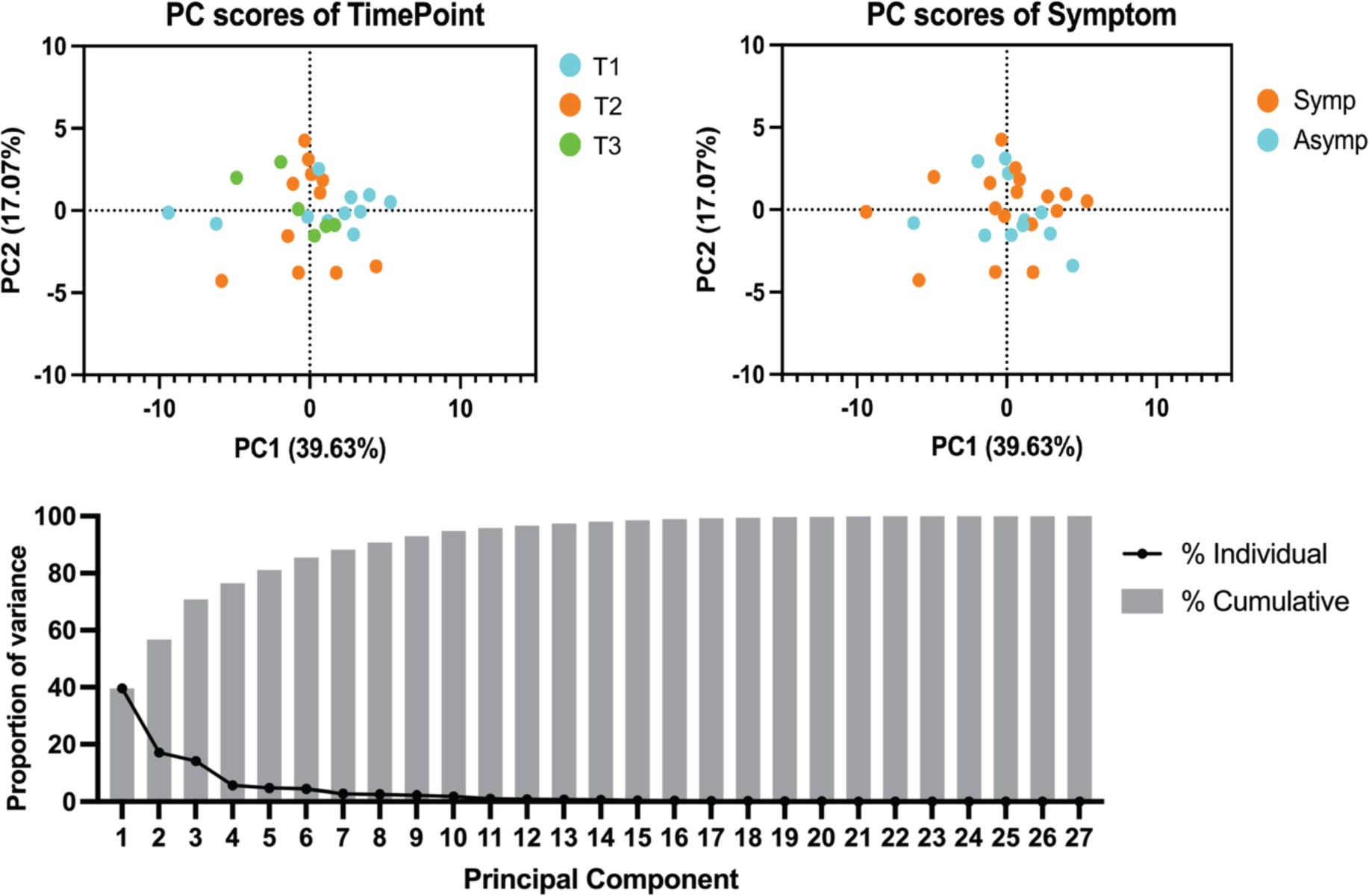
Principal Component Analysis (PCA) of experimental variant effects between individual samples in CyTOF analysis. No significant difference was found in a reference sample included between time points and clinical groups analyzed by two-way ANOVA (Tukey’s multiple comparisons test).

**Figure S2.**
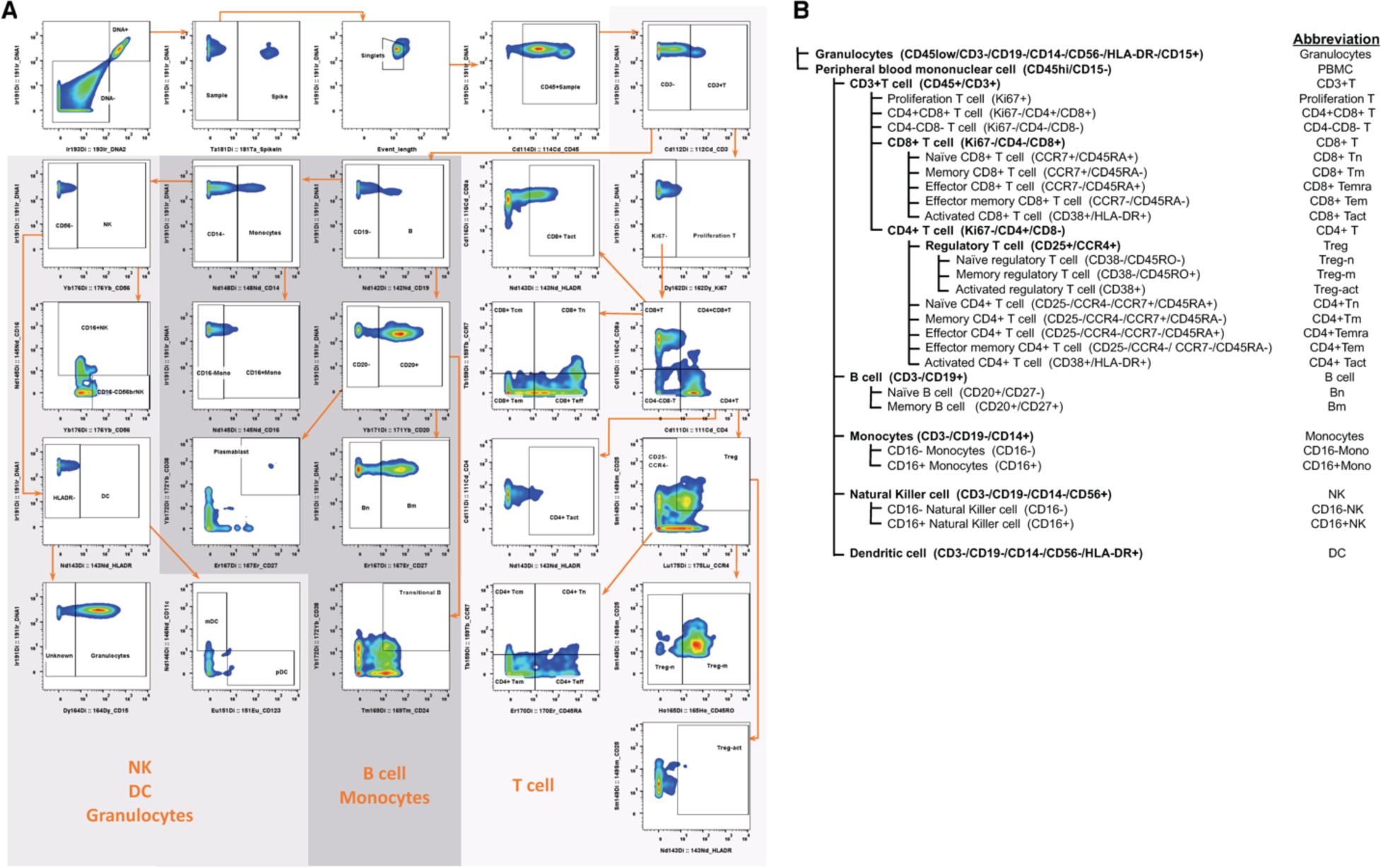
CyTOF Gating strategy. Representative gating strategy for CyTOF (A) and subset population definitions defined by hierarchical gating (B).

**Figure S3.**
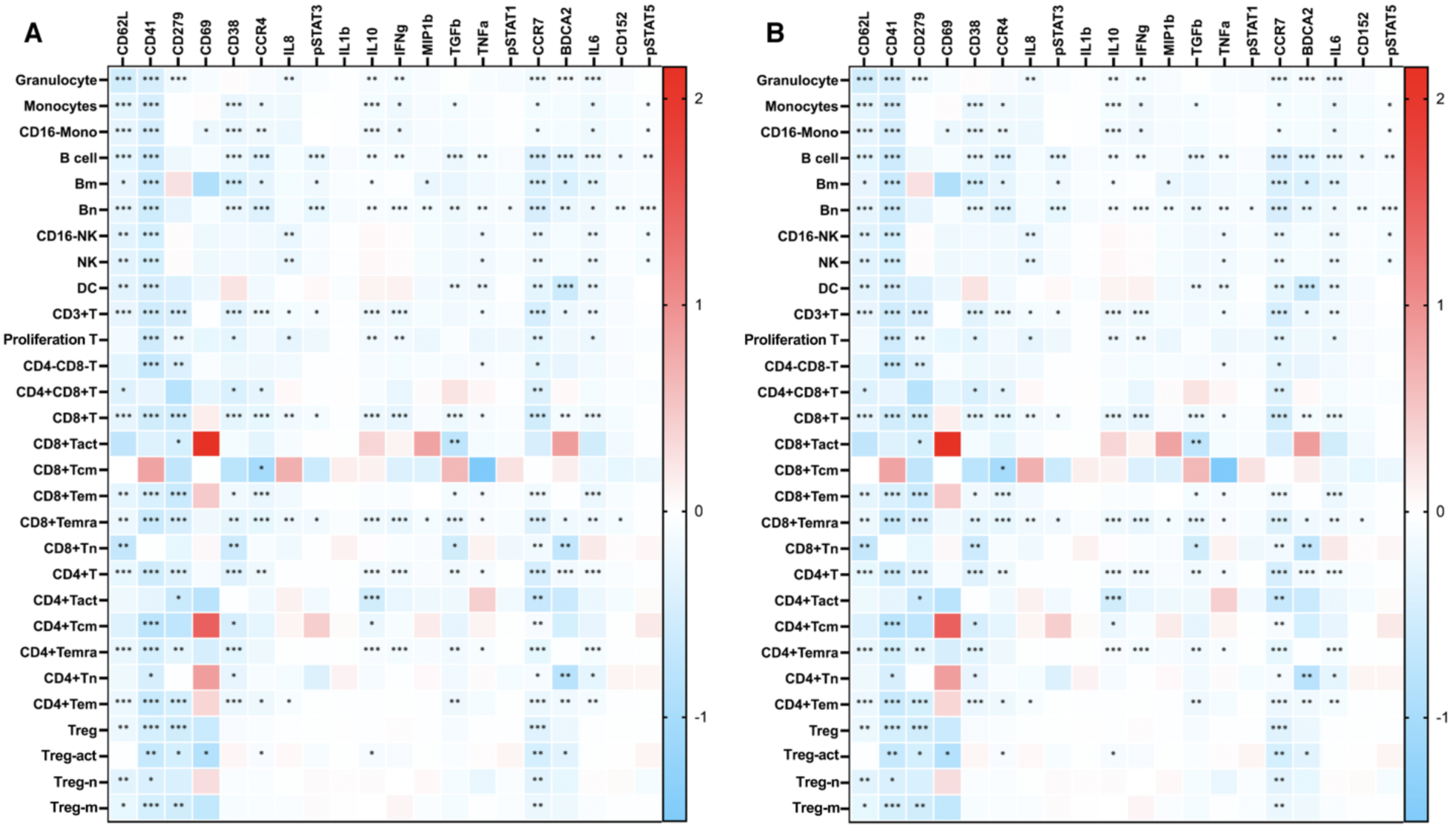
Altered immune functional markers during acute WNV infection. Whole blood cells from WNV patients (n=11) were labeled with metal-conjugated antibodies and analyzed by mass cytometry to quantify levels of 20 functional markers in 28 cell subsets. (A) Change in median channel intensity (A) Day 1 vs Year 1; (B) Month 3 vs Year 1 (ratio log10 acute/convalescent). Significance by Generalized Linear Mixed Model with * p<0.05, ** p<0.01 and *** p<0.001.

**Figure S4.**
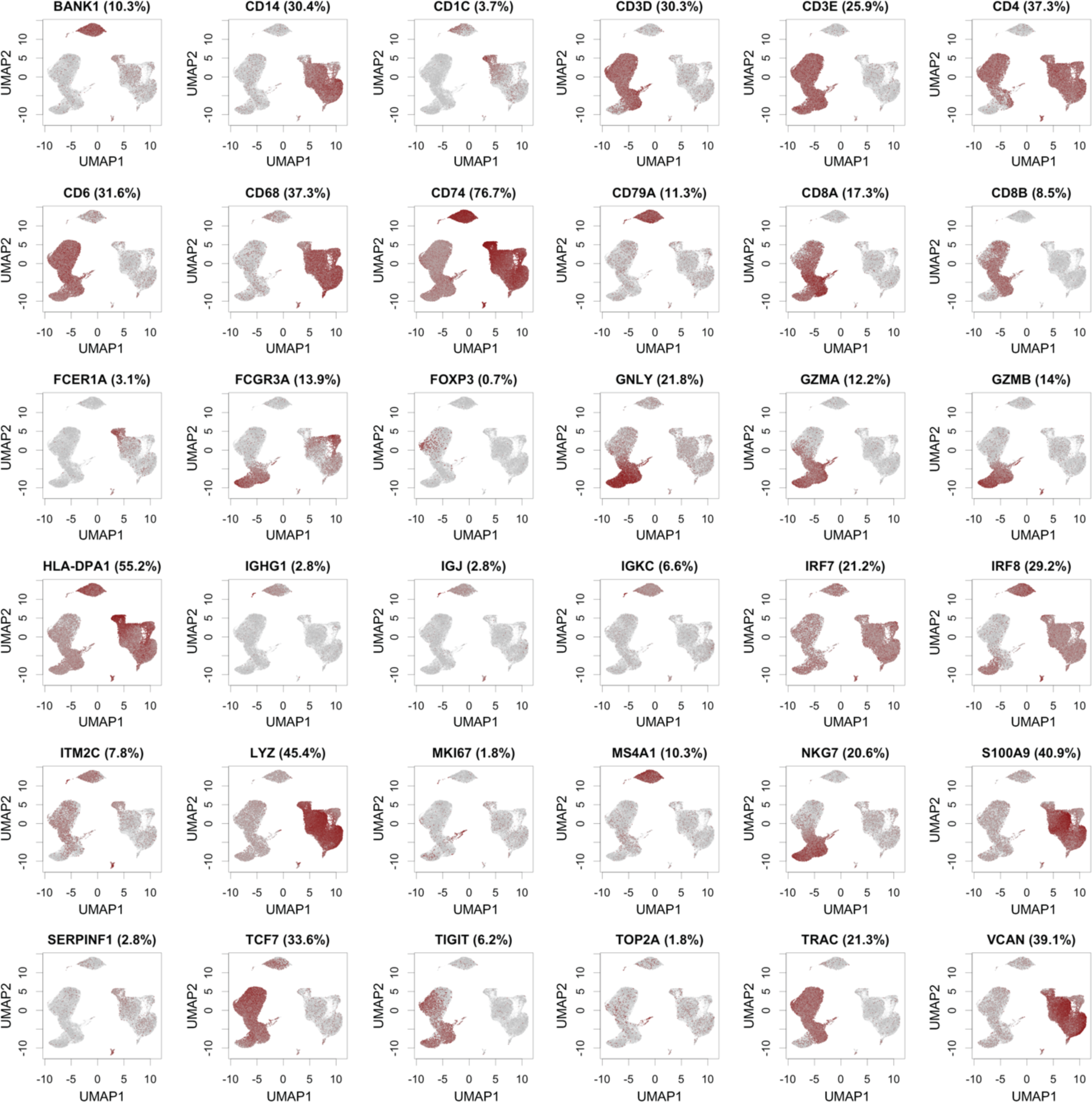
UMAP plots for 40 cell type marker genes. These 40 marker genes were used to define the 11 cell types in the Seq-Well data. Cells with non-zero expression are shown in red and their percentage is shown in the parentheses in the plot title.

**Figure S5. Gene expression profiles of the top 10 genes in each module for each cell type.** See Supplemental Data.

## Supplemental Tables

**Table S1.**
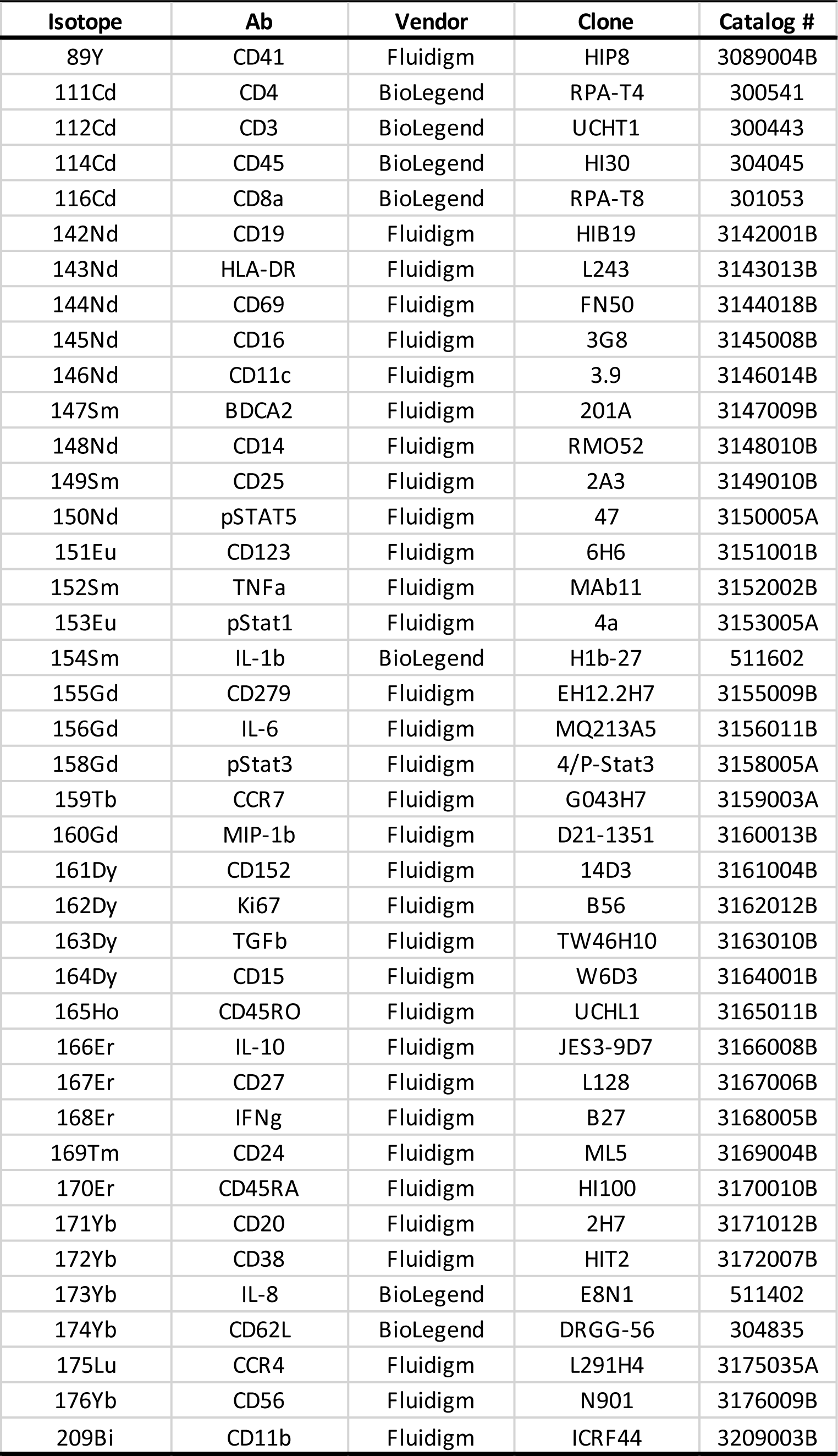
Antibodies used for mass cytometry (CyTOF)

**Table S2.** Lists of the top 10 genes for each differential SeqWell module

| Cell Type | Module | Top 10 Genes |  |  |  |  |  |  |  |  |  |
| --- | --- | --- | --- | --- | --- | --- | --- | --- | --- | --- | --- |
| CD14+_Monocytes | 1 | TIMP1 | JARID2 | SDC2 | THBS1 | SEMA6B | CTSL | PLIN2 | DHRS3 | SLA | SERPINB2 |
| CD16+_Monocytes | 1 | LCP1 | ACTB | FYB | CYBB | FGL2 | CAP1 | SAMHD1 | FAM65B | CSF1R | CPPED1 |
| CD16+_Monocytes | 2 | FOSB | FOS | ZFP36 | JUN | DUSP1 | NR4A2 | CD83 | SGK1 | RGS1 | TIPARP |
| CD16+_Monocytes | 3 | ANXA1 | LMNA | VIM | PTGES | ATP2B1 | GABARAPL1 | EMP1 | FTH1 | TREM1 | MYADM |
| CD4+_T_Cells | 2 | FOS | JUNB | PPP1R15A | FOSB | JUN | IER2 | CITED2 | SOCS3 | HEXIM1 | IRS2 |
| CD8+_T_Cells | 1 | GZMH | GZMA | CD52 | B2M | AHNAK | ANXA6 | FGFBP2 | S100A4 | GNLY | IL32 |
| CD8+_T_Cells | 3 | SLC7A5 | FTH1 | GABARAPL1 | SKIL | FAM102A | SKI | PDE4B | PPP1R16B | METRNL | GGA2 |
| cDCs | 1 | S100A9 | FCN1 | S100A8 | VCAN | NCF2 | CTSS | CYBB | S100A4 | AHNAK | CSF1R |
| cDCs | 4 | CXCL16 | FTH1 | JARID2 | NR4A3 | SRGN | DUSP4 | INSIG1 | GABARAPL1 | MALT1 | SAT1 |
| NK_Cells | 3 | FOS | JUN | DUSP1 | IER2 | PPP1R15A | NFKBIA | JUNB | ZFP36 | IER5 | GADD45B |
| pDCs | 1 | FOS | FOSB | KLF4 | JUN | ZFP36 | RGS1 | DUSP1 | JUNB | RGS2 | PPP1R15A |
| Plasma_B_Cells | 6 | RPN2 | CD38 | TPST2 | C19orf10 | CPEB4 | RRBP1 | SEC24A | PRDX4 | LAMC1 | ITM2B |

**Table S3.**
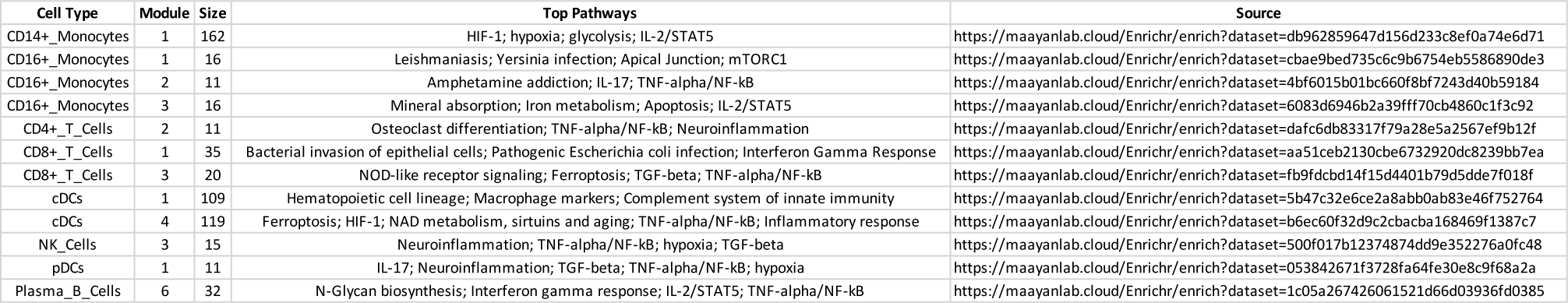
Top enriched pathways by Enrichr for the 12 modules of interest

**Table S4** Protein Matrix: Transitions, peptides and proteins from the MStern proteomic analysis See Supplemental Data.

**Table S5.**
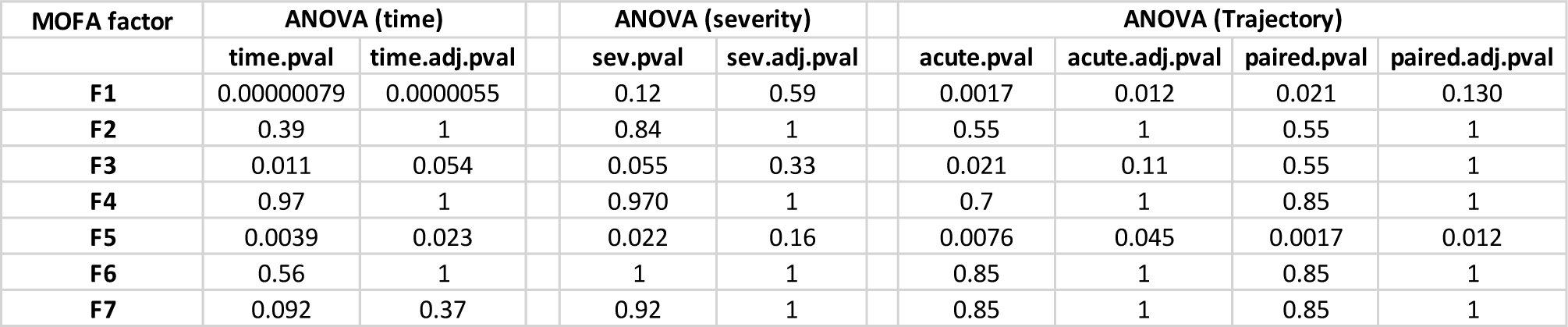
Analysis of Variance for MOFA factors

**Table S6** Signature features of MOFA factor 1 See Supplemental Data.

**Table S7** Signature features of MOFA factor 5 See Supplemental Data.

