## Supplemental Software for "Early cellular and molecular signatures correlate with severity of West Nile Virus infection": demo.m

```
% permutation tests 


%load 
p29= readtable('wnv_datamatrix_29.xlsx','sheet','pop29');


%%
time_subset = 1 ;
iterations = 1000 ;

t  = permutation_wnv_ind(p29, time_subset, upper_thresh, lower_thresh, iterations)
writetable(t,'PermutationByIndividualCombos.xlsx');
```
