## Supplemental Software for "Early cellular and molecular signatures correlate with severity of West Nile Virus infection": permutation_wnv_ind.m

```
function [t]  = permutation_wnv_ind(data, time_subset, iterations)
t = [];
data_width = width(data) ;

% extract out time subset
subdata = data(data.time == time_subset,:) ;

%extract labels


% compute mean per measure for a and s
am = mean(subdata{ismember(subdata.SympStat, 'Asymptomatic'),4:data_width},'omitnan') ;
sm = mean(subdata{ismember(subdata.SympStat,'Symptomatic'),4:data_width},'omitnan') ;


% compute fold
fold = sm./am ;

iterated_fold = zeros(iterations,numel(fold)) ;


for i = 1 : iterations
    %extract data
    temp_table = subdata ;
    
    %randomize the orders
    %  scramble labels
    group_list = temp_table.SympStat ;
    group_list_rand = group_list(randperm(length(group_list))) ;
    temp_table.group_rand = group_list_rand ;
    
    amr = mean(temp_table{ismember(temp_table.group_rand, 'Asymptomatic'),4:data_width},'omitnan') ;
    smr= mean(temp_table{ismember(temp_table.group_rand,'Symptomatic'),4:data_width},'omitnan') ;

    % compute fold
    iterated_fold(i,:) =smr./amr ;

    % count number > 1.1 and < .9

end 

cdist = zeros(1,numel(fold)) ;
for j = 1 : numel(fold)
    cdist(j) =sum(iterated_fold(:,j) > fold(1,j)) ;
    
end 
cdist = cdist / iterations ;


t = table([subdata.Properties.VariableNames(4:data_width)]', cdist');


end
```
