## Supplemental Software for "Early cellular and molecular signatures correlate with severity of West Nile Virus infection": readme.txt

1. System requirements
This software was designed and tested on Matlab 2021a running on Windows 10 PC.  

2. Installation guide
The matlab function permutation_wnv_ind.m. performs the permutation test.  The function needs to be callable within the Matlab environment.  This can be done by navigating to the directory where the function is located or it can be added to the Matlab path (see https://www.mathworks.com/help/matlab/ref/addpath.html) .  No further installation is necessary

3. Demo
A demo script (demo.m) and the associated data set (wnv_datamatrix_29.xlsx) are provided.  The demo.m file demonstrates setting parameters and calling the function.  These commands can be run as a batch within a script (as per the demo.m) or individually from the command line.

4. Instructions for use
permutation_wnv_ind requires three arguments in order:
A)	A data set formatted as a Matlab table structure.  This minimally needs to include columns:
	a.	SympStat – the symptomatic status of the individual
	b.	ID – patient ID information
	c.	Time – indexed observation 
	d.	One or more CyTOF activation values corresponding to combinations of cells and markers
B)	Time_subset – this indicates which of the observations to analyze.  The data includes 3 observations, but the permutation was only performed for the first. 
C)	Iterations- this is a integer indicating the number of permutation iterations to run.  

The demo.m file demonstrates loading a table from an excel spreadsheet, setting the parameters, and calling the function. 
The function computes the ratio of CyTOF activation between the Symptomatic and Asymptomatic individuals a each cell and marker combination.  It then permutes the group labels n=iterations times.  
The function generates an output that details, for each cell and marker, the proportion of iterations for which the ratio (given the permuted labels) was greater than the actual ratio.  

MIT License

Copyright (c) 2022 Brent Vander Wyk

Permission is hereby granted, free of charge, to any person obtaining a copy
of this software and associated documentation files (the "Software"), to deal
in the Software without restriction, including without limitation the rights
to use, copy, modify, merge, publish, distribute, sublicense, and/or sell
copies of the Software, and to permit persons to whom the Software is
furnished to do so, subject to the following conditions:

The above copyright notice and this permission notice shall be included in all
copies or substantial portions of the Software.

THE SOFTWARE IS PROVIDED "AS IS", WITHOUT WARRANTY OF ANY KIND, EXPRESS OR
IMPLIED, INCLUDING BUT NOT LIMITED TO THE WARRANTIES OF MERCHANTABILITY,
FITNESS FOR A PARTICULAR PURPOSE AND NONINFRINGEMENT. IN NO EVENT SHALL THE
AUTHORS OR COPYRIGHT HOLDERS BE LIABLE FOR ANY CLAIM, DAMAGES OR OTHER
LIABILITY, WHETHER IN AN ACTION OF CONTRACT, TORT OR OTHERWISE, ARISING FROM,
OUT OF OR IN CONNECTION WITH THE SOFTWARE OR THE USE OR OTHER DEALINGS IN THE
SOFTWARE.
