## Supplemental Software for "Early cellular and molecular signatures correlate with severity of West Nile Virus infection": helpers.R

```
sourcedata <- function(path = ''){
  
  load(paste0(path,"seqWell_summary.RData"))
  data6 = mean_gene
  data6 = t(data6)
  
  data7 = read.csv(paste0(path,"metabolomics.merged_0615.csv"))
  data7_info = read.csv(paste0(path,"metabolomics_sample_info.csv"))
  data7_info = data7_info[1,!is.na(data7_info[1,])]
  data7 = cbind(data7[,c(5,6)],data7[,colnames(data7_info)])
  colnames(data7)[3:ncol(data7)] =sapply(data7_info, as.character)
  data7 = data7[,-2]
  data7[,1] = as.character(data7[,1])
  data7= data7[data7[,1]!="",]
  rownames(data7) = data7[,1]
  data7 = data7[,-1]
  data7 = t(data7)
  
  tmp = strsplit(patient_state_timepoint0,'_')
  symptom0  = sapply(tmp, function(z) z[2])
  patient0  = sapply(tmp, function(z) z[3])
  time0  = sapply(tmp, function(z) z[1])
  
  sample_id6 = paste0(patient0,'_',time0)
  
  data1 = read.csv(paste0(path,"CyTOF_Frequency_total0505.csv"), header = T)
  data2 = read.csv(paste0(path,"CyTOF_Frequency_PBMC0505.csv"), header = T)
  data3 = read.csv(paste0(path,"CyTOF_MeanFunctionalMarkers1209.csv"), header = T)
  colnames1 =  colnames(data1)[-1]
  rownames(data1) = data1[,1]
  data1 = data1[,-1]
  colnames2 =  colnames(data2)[-1]
  rownames(data2) = data2[,1]
  data2 = data2[,-1]
  colnames3 =  colnames(data3)[-1]
  rownames(data3) = data3[,1]
  data3 = data3[,-1]
  data4 = readRDS(paste0(path, 'seqwell_lexi.RDS'))
  
  dat4.patient = data4$SubjectID
  dat4.symptom = data4$Sevs
  dat4.times = data4$Times
  
  data5 = read.csv(paste0(path,"WNVsamples_Proteomics_SteenLab.csv"), header = T)
  dat5.symptom =  as.vector(colnames(data5)[-1])
  dat5.times= data5[1,-1]
  dat5.times = sapply(dat5.times, function(z) as.character(z))
  dat5.patient = data5[3,-1]
  data5 = data5[-c(1:3),]
  rownames(data5) = data5[,1]
  data5 = data5[,-1]
  data5 = apply(data5, c(1,2), as.numeric)
  fillin = apply(data5 ,1,min, na.rm = T)/2
  for(j in 1:nrow(data5)){
    ll = which(is.na(data5[j,]))
    data5[j,ll] = fillin[j]/2
  }
  data5 = log(data5+1)
  data5 = t(data5)
  
  tmp = strsplit(rownames(data1),"_")
  dat1.patient = sapply(tmp,function(z) z[2])
  dat1.times = sapply(tmp,function(z) z[3])

  
  dat1.times[dat1.times=='Day1'] = 'T1'
  dat1.times[dat1.times=='Month3'] = 'T2'
  dat1.times[dat1.times=='Year1'] = 'T3'
  
  tmp = strsplit(rownames(data2),"_")
  dat2.patient = sapply(tmp,function(z) z[2])
  dat2.times = sapply(tmp,function(z) z[3])
  
  dat2.times[dat2.times=='Day1'] = 'T1'
  dat2.times[dat2.times=='Month3'] = 'T2'
  dat2.times[dat2.times=='Year1'] = 'T3'
  
  tmp = strsplit(rownames(data3),"_")
  dat3.patient = sapply(tmp,function(z) z[2])
  dat3.times = sapply(tmp,function(z) z[3])
  dat3.times[dat3.times=='Day1'] = 'T1'
  dat3.times[dat3.times=='Month3'] = 'T2'
  dat3.times[dat3.times=='Year1'] = 'T3'

  
  dat5.symptom = sapply(strsplit(dat5.symptom,'[.]'), function(z) z[1])
  dat5.patient = sapply(strsplit(sapply(dat5.patient[1,], function(z)as.character(z)),'[.]'), function(z) z[1])
  
  tmp =strsplit(rownames(data7),'_')
  dat7.patient =sapply(tmp, function(z) z[[1]])
  dat7.times =sapply(tmp, function(z) z[[2]])
  dat7.times = ifelse(dat7.times=="T1", "T2", "T1")
  sample_id1 = paste0(dat1.patient,'_',dat1.times)
  sample_id2 = paste0(dat2.patient,'_',dat2.times)
  sample_id3 = paste0(dat3.patient,'_',dat3.times)
  sample_id4 = paste0(dat4.patient,'_',dat4.times)
  sample_id5 = paste0(dat5.patient,'_',dat5.times)
  sample_id7 = paste0(dat7.patient,'_',dat7.times)
  ll1 = c()
  ll2 = c()
  ll3 = c()
  ll5 = c()
  ll6 = c()
  ll7 = c()
  for(i in 1:length(sample_id4)){
    ll1 = c(ll1, which(sample_id1==sample_id4[i]))
    ll2 = c(ll2, which(sample_id2==sample_id4[i]))
    ll3 = c(ll3, which(sample_id3==sample_id4[i]))
    ll5 = c(ll5, which(sample_id5==sample_id4[i]))
    ll6 = c(ll6, which(sample_id6==sample_id4[i]))
    if(sample_id4[i] %in%sample_id7){
        ll7 =c(ll7, which(sample_id7==sample_id4[i]))
    }else{
        ll7 = c(ll7, NA)
    }

  }
  data1 = data1[ll1,]
  data2 = data2[ll2,]
  data3 = data3[ll3,]
  data5 = data5[ll5,]
  data6 = data6[ll6,]
  data7 = data7[ll7,]
  metas = data4[,c(2:4)]
  data4 = data4[,-(1:4)]
  rownames(data7) =sample_id4
  data7 = apply(data7, c(1,2), as.numeric)
  
  
  data = list(cytof_total = data1, cytof_pbmc = data2,
              cytof_markers = data3, seqwell = data4[,grepl( 'freq',colnames(data4))],
              seqwell_mean = data4[,grepl( 'mean',colnames(data4))],
              seqwell_median = data4[,grepl( 'median',colnames(data4))],
              proteomics = data5, seqwell_global_mean = data6,
              metabolomics = data7)
  for(l in 1:length(data)){
    rownames(data[[l]]) = paste(metas[,1], metas[,3], sep='-')
  }
  return(list(data = data, metas = metas))
}


data_filtering <- function(data, missing_percent = 0.25,
                           RNA_quantile = 0.8){
  for(i in 1:length(data)){
    tmp = apply(is.na(data[[i]]),2,sum)
    data[[i]] = data[[i]][,tmp<=(nrow(data[[i]]) * missing_percent)]
  }
  tmp = data[["seqwell_global_mean"]]
  tmp1 = apply(tmp,1,var)
  thr = quantile(tmp1,  RNA_quantile )
  data[["seqwell_global_mean"]] =data[["seqwell_global_mean"]][,tmp1>=thr]
  for(l in 1:length(data)){
    data[[l]] = apply(data[[l]],c(1,2), as.numeric)
    v1 = apply(data[[l]],2,sd, na.rm =T)
    print(sum(v1==0))
    data[[l]] = data[[l]][,!is.na(v1)]
    data[[l]] = data[[l]][,v1[!is.na(v1)]!=0]
    v1 = apply(data[[l]],2,sd, na.rm =T)
    print(sum(v1==0))
    colnames(data[[l]]) = paste0(as.character(l),":", colnames(data[[l]]))
  }
  return(data)
}


data_imputation <- function(data, outfile = "mofa_imputation.hf5", rerun = FALSE){
  ###mofa: data imputation
  mofa_data = list()
  mofa_data[[1]] = t(data$cytof_total)
  mofa_data[[2]] = t(data$cytof_markers)
  mofa_data[[3]] = t(data$seqwell)
  mofa_data[[4]] = t(data$seqwell_mean)
  mofa_data[[5]] = t(data$proteomics)
  mofa_data[[6]] = t(data$seqwell_global_mean)
  mofa_data[[7]] = t(data$metabolomics)
  mofa_data[[8]] = t(data$cytof_pbmc)
  
  names(mofa_data) = c("cytof_total",  "cytof_markers_mean",
    "seqwell_total" ,"seqwell_module_mean",
    "proteomics", "seqwell_global_mean", "metabolomics","cytof_pbmc")
  
  
  MOFAobject <- create_mofa(mofa_data)
  data_opts <- get_default_data_options(MOFAobject)
  model_opts <- get_default_model_options(MOFAobject)
  model_opts$num_factors= nrow(data$cytof_total)-5
  train_opts <- get_default_training_options(MOFAobject)
  train_opts$drop_factor_threshold <- 0.01
  train_opts$maxiter = 500
  train_opts$convergence_mode="slow"
  train_opts$startELBO = 50
  
  train_opts$seed <- 2021
  
  MOFAobject <-  prepare_mofa(
    object = MOFAobject,
    data_options = data_opts,
    model_options = model_opts,
    training_options = train_opts
  )
  
  mofa_file = paste0("/gpfs/ysm/project/guan_leying/lg689/WNV/",  outfile)
  if((!file.exists(mofa_file)) | (rerun)){
    MOFAobject <- run_mofa(MOFAobject, outfile = mofa_file)
  }else{
    MOFAobject <- MOFA2::load_model(file =  mofa_file, remove_inactive_factors = T)
  }
  
  MOFAobject = impute(MOFAobject)
  return(MOFAobject)
}

normalization <- function(X, method = "quantile", stabilized = 0.1){
  if(method == "quantile"){
    p = ncol(X)
    n = nrow(X)
    a0 = qnorm((c(1:n)-0.5)/(n))
    for(j in 1:p){
      X[,j] = a0[rank(X[,j])]
    }
  }
  return(X)
}

mofa_run <- function(mofa_data, outfile = "mofa_imputation.hf5", 
                     rerun = FALSE, maxiter = 1000, drop_factor_threshold = 0.01,
                     num_factors = 15){
  MOFAobject <- create_mofa(mofa_data)
  data_opts <- get_default_data_options(MOFAobject)
  model_opts <- get_default_model_options(MOFAobject)
  model_opts$num_factors= num_factors
  train_opts <- get_default_training_options(MOFAobject)
  train_opts$drop_factor_threshold <- drop_factor_threshold
  train_opts$maxiter = maxiter
  train_opts$convergence_mode="slow"
  train_opts$startELBO = 50
  
  train_opts$seed <- 2021
  
  MOFAobject <-  prepare_mofa(
    object = MOFAobject,
    data_options = data_opts,
    model_options = model_opts,
    training_options = train_opts
  )
  
  mofa_file = paste0("/gpfs/ysm/project/guan_leying/lg689/WNV/",  outfile)
  if((!file.exists(mofa_file)) | (rerun)){
    MOFAobject <- run_mofa(MOFAobject, outfile = mofa_file)
  }else{
    MOFAobject <- MOFA2::load_model(file =  mofa_file, remove_inactive_factors = T)
  }
  
  MOFAobject = impute(MOFAobject)
  return(MOFAobject)
}


residualize<- function(imputed_moda_data, metas){
  moda_data2 = list()
  for(i in 1:length(imputed_moda_data)){
    tmp = imputed_moda_data[[i]]$group1
    ll = (metas$Times =='T1')
    tmp[,ll] = tmp[,ll]-apply(tmp[,ll],1,mean)
    ll = (metas$Times =='T2')
    tmp[,ll] = tmp[,ll]-apply(tmp[,ll],1,mean)
    ll = (metas$Times =='T3')
    tmp[,ll] = tmp[,ll]-apply(tmp[,ll],1,mean)
    moda_data2[[i]] = tmp
  }
  moda_data2
}

effect_tests <- function(metas_use, allfactors,type = 'acute', acute_test = 'spearman'){
  pvals_all = rep(NA, ncol(allfactors))
  pvals_times = rep(NA, ncol(allfactors))
  pvals_symptom = rep(NA, ncol(allfactors))
  for(k in 1:ncol(allfactors)){
    lf = allfactors[,k]
    #all effects
    y = lf
    if(type == 'acute'){
      x = ifelse(metas_use$Sevs =='WNA', 0, 1)
      tmp = cor.test( x, y, method = acute_test)
      pvals_symptom[k] = tmp$p.value
    }else{
      x = matrix(0, nrow = length(y), ncol = 5)
      x[metas_use$Sevs != 'WNA',1] = 1
      x[metas_use$Times=='T2' & metas_use$Sevs == 'WNA',2] = 1
      x[metas_use$Times=='T3' & metas_use$Sevs == 'WNA',3] = 1
      x[metas_use$Times=='T2' & metas_use$Sevs != 'WNA',4] = 1
      x[metas_use$Times=='T3' & metas_use$Sevs != 'WNA',5] = 1
      tmp = summary(lm(y~x))
      pvals_all[k] = pf(tmp$fstatistic[1], df1 = tmp$fstatistic[2], df2 = tmp$fstatistic[3], lower.tail = F)
      # time effects 
      tmp = lm(y~x)
      # if all time is the same
      tmp1 = lm(y~x[,1])
      a = (sum(tmp1$residuals^2) - sum(tmp$residuals^2))/(6-2)
      b =  sum(tmp$residuals^2)/(length(y)-6)
      pvals_times[k] = pf(a/b, df1 = 6-2, df2 =(length(y)-6) , lower.tail = F)
      # symptom effects
      y = lf
      x = matrix(0, nrow = length(y), ncol = 5)
      x[metas_use$Times == 'T2',1] = 1
      x[metas_use$Times == 'T3',2] = 1
      #diease * T1, disease * T2, disease * T3
      x[metas_use$Times == 'T1' &  metas_use$Sevs !='WNA',3] = 1
      x[metas_use$Times == 'T2' &  metas_use$Sevs !='WNA',4] = 1
      x[metas_use$Times == 'T3' &  metas_use$Sevs !='WNA',5] = 1
      tmp = lm(y~x)
      tmp1 = lm(y~x[,1:2])
      a = (sum(tmp1$residuals^2) - sum(tmp$residuals^2))/(6-3)
      b = sum(tmp$residuals^2)/(length(y)-6)
      pvals_symptom[k] = pf(a/b, df1 = 6-3, df2 = (length(y)-6), lower.tail = F)
    }
  }
  pvals = cbind(cbind(pvals_symptom, pvals_times),pvals_all)
  colnames(pvals) = c('symptom', 'times', 'all')
  return(pvals )
}
```
