## Supplemental Software for "Early cellular and molecular signatures correlate with severity of West Nile Virus infection": joint_analysis_farnam_share.nb.html

R Notebook


Code 

- Show All Code
- Hide All Code
- Download Rmd

### R Notebook

### Loading Data and packages


```
#path = '/Users/lg689/Dropbox/COVID19/WNV/data/code'
#setwd(path)
library(readr)
library(reticulate)
reticulate::use_virtualenv("~/project/python3.7venv", required = TRUE)
require(MOFA2)
library(ggplot2)
library(tidyr)
source("helpers.R")
predict = stats::predict
if(file.exists("processed_filtered_data.RData")){
  load("processed_filtered_data.RData")
}else{
  
  if(file.exists("processed_data.RDS")){
    processed_info = readRDS(file = "processed_data.RDS")
  }else{
    processed_info <- sourcedata(path ="")
    saveRDS(processed_info, file = 'processed_data.RDS')
  }
  data = processed_info$data
  metas = processed_info$metas
  
  data["seqwell_median"] = NULL
  
  data_processed = data_filtering(data = data, missing_percent = 0.3,
                                  RNA_quantile = 0.8)
  
  sapply(data_processed , dim)
  tmp = sapply(data_processed, dim)
  rownames(tmp) = c("nsample", "nfeature")
  
  save(data_processed, metas, file = "processed_filtered_data.RData")
}

###mofa: data imputation
analysis_date = "_1209"
MOFAobject = data_imputation(data= data_processed, outfile = paste0("mofa_imputation",analysis_date,".hf5"), rerun = F)

calculate_variance_explained(MOFAobject)

factors_joint = get_factors(MOFAobject)


data_imputed = get_imputed_data(MOFAobject)
data_mofa = get_data(MOFAobject)

for(i in 1:length(data_imputed)){
  data_imputed[[i]] = t(data_imputed[[i]][[1]])
  data_mofa[[i]] = t(data_mofa[[i]][[1]])
}
sapply(data_mofa,dim)
sapply(data_imputed ,dim)


for(i in 1:length(data_imputed)){
tmp = normalization(data_imputed[[i]], method = "quantile")
tmp[is.na(as.matrix(data_mofa[[i]]))] = NA
data_mofa[[i]] = tmp
}
```

### Module identification: meta-joint and joint-estimation

In this section, we derive two sets of factors: Derive \(K\) factors with MOFA pooling all data together.

#### Joint omics analysis


```
time_used = c('T1', 'T2','T3')

ll_use =  which(metas$Times %in% time_used)

mofa_data = list()
for(k in 1:length(data_mofa)){
  mofa_data[[k]] =  t(data_mofa[[k]][ll_use,])
}
names(mofa_data) = names(data_mofa)
  

MOFAobject_joint_all = mofa_run(mofa_data, outfile = paste0("mofa_model_",type,analysis_date, ".hf5"), rerun = F,
maxiter = 5000, drop_factor_threshold = 1e-3,num_factors = 25)

library(RColorBrewer)

sample_metadata <- data.frame(
  sample = samples_names(MOFAobject_joint_all)[[1]],
  Sev =ifelse(metas$Sev[ll_use] == "WNA","0","1"),
  Time = metas$Time[ll_use]
)

samples_metadata(MOFAobject_joint_all) <- sample_metadata

#plot_factor_cor(MOFAobject_joint_all)
factors_joint = get_factors(MOFAobject_joint_all)
calculate_variance_explained(MOFAobject_joint_all)
tmp = calculate_variance_explained(MOFAobject_joint_all)
tmp$r2_per_factor$group1
apply(tmp$r2_per_factor$group1,1,mean)

#  Factor1  Factor2  Factor3  Factor4  Factor5  Factor6  Factor7  Factor8 
#     14.1     12.7      5.9      4.7      4.0      2.6      2.4      2.0 
#  Factor9 Factor10 Factor11 Factor12 Factor13 Factor14 Factor15 Factor16 
#      2.0      1.8      1.6      1.3      1.3      1.2      1.2      1.1 
# Factor17 Factor18 Factor19 Factor20 Factor21 Factor22 Factor23 
#      1.1      0.9      0.6      0.6      0.6      0.5      0.5 
     
write.csv(round(tmp$r2_per_factor$group1,2), file = "variance_explained01026.csv")
```

### Testing for associations between MOFA factors and responses

#### Two-way anova


```
factor_rank = 7  #maximum number of factors suggested by MOFA
metas_use = metas[ll_use,]
allfactors = factors_joint[[1]][,c(1:factor_rank)]
allfactors=normalization(allfactors, method = "quantile") #quantile normailziation to remove outliers

#two-way anova tests
pvals_pearson = effect_tests(metas_use=metas_use, allfactors=allfactors,type = 'all', acute_test = 'pearson')
pvals_pearson
apply(pvals_pearson,2,function(z) qvalue(as.vector(z), fdr.level = 0.05, pi0 = 1)$qvalue)
qvalue::qvalue(as.vector(pvals_pearson[,1]), fdr.level = 0.05, pi0 = 1)
pvals_pearson1 = matrix(0, ncol = nrow(pvals_pearson), nrow = 4)
pvals_pearson1[1,] =   pvals_pearson[,2]
pvals_pearson1[3,] =   pvals_pearson[,1]
pvals_pearson1[2,]=p.adjust( pvals_pearson1[1,])
pvals_pearson1[4,]=p.adjust( pvals_pearson1[3,])
pvals_pearson1=formatC(pvals_pearson1, format = "e", digits = 1)

#      [,1]      [,2]      [,3]      [,4]      [,5]      [,6]      [,7]     
# [1,] "7.9e-07" "3.9e-01" "1.1e-02" "9.7e-01" "3.9e-03" "5.6e-01" "9.2e-02"
# [2,] "5.5e-06" "1.0e+00" "5.4e-02" "1.0e+00" "2.3e-02" "1.0e+00" "3.7e-01"
# [3,] "1.2e-01" "8.4e-01" "5.5e-02" "9.7e-01" "2.2e-02" "1.0e+00" "9.2e-01"
# [4,] "5.9e-01" "1.0e+00" "3.3e-01" "1.0e+00" "1.6e-01" "1.0e+00" "1.0e+00"

colnames(pvals_pearson1) = paste0("F",1:factor_rank)
rownames(pvals_pearson1) = c("time.pval", "time.adj.pval", "sev.pval", "sev.adj.pval")
print(pvals_pearson)
rownames(pvals_pearson) = paste0("F",1:nrow(pvals_pearson))
pvals_pearson1 = t(pvals_pearson1)
write.csv(pvals_pearson1, file = paste0("supplement_pvalues_totaleffects",analysis_date,".csv"))
```

#### Paired difference


```
allfactors = factors_joint[[1]][,c(1:factor_rank)]
allfactors=normalization(allfactors, method = "quantile")
aa = intersect(metas$SubjectID[metas$Times == "T1"],
               metas$SubjectID[metas$Times == "T2"])
ll1 = c()
ll2 = c()
for(i in 1:length(aa)){
  ll1 = c(ll1, which(metas$Times == "T1" & (metas$SubjectID==aa[i])))
  ll2 = c(ll2, which(metas$Times == "T2" & (metas$SubjectID==aa[i])))
}
x1 = allfactors[ll1,]
x2 = allfactors[ll2,]
yy = metas$Sev[ll1]
yy = ifelse(yy=="WNA",0,1)
Deltas = array(0, dim = dim(allfactors))
Deltas = x2-x1
pvals_paired = matrix(0, ncol = 2, nrow =ncol(Deltas))
pvals_acute = matrix(0, ncol = 2, nrow =ncol(Deltas))
for(k in 1:ncol(x1)){
  z = Deltas[,k]
  tmp = cor.test(z,yy, method = "pearson")
  pvals_paired[k,1] =  tmp$p.value
  tmp = cor.test(z,yy, method = "spearman")
  pvals_paired[k,2] =  tmp$p.value
  tmp = cor.test(x1[,k],yy, method = "spearman")
  pvals_acute[k,1] = tmp$p.value
}
pvals_acute[,2] =p.adjust(pvals_acute[,1]) 
# 0.021, factor1, 0.13
# 0.001713, factor 5, 0.012
pvals_paired1 = matrix(0, ncol = 4, nrow =ncol(Deltas))
pvals_paired1[,1] = pvals_paired[,1]
pvals_paired1[,2] = p.adjust(pvals_paired1[,1])
pvals_paired1[,3] = pvals_paired[,2]
pvals_paired1[,4] = p.adjust(pvals_paired1[,3])

pvals_all = data.frame(pvals_pearson1)
pvals_all$acute.pval = formatC(pvals_acute[,1], format = "e", digits = 1)
pvals_all$acute.adj.pval = formatC(pvals_acute[,2] , format = "e", digits = 1)
pvals_all$paired.pval = formatC(pvals_paired1[,3], format = "e", digits = 1)
pvals_all$paired.adj.pval = formatC(pvals_paired1[,4], format = "e", digits = 1)

write.csv(pvals_all, file = paste0("supplement_pvalues_all",analysis_date,".csv"), quote = F, row.names = T)
```

#### Plots on factors of interests


```
library(tidyr)
allfactors = factors_joint[[1]][,1:factor_rank]
allfactors=normalization(allfactors, method = "quantile")
colnames(allfactors)=paste0('Factor', c(1:ncol(allfactors)))
allfactors= data.frame(allfactors)
allfactors =allfactors[ll_use,]
allfactors$time = metas$Times[ll_use]
allfactors$symptom = factor(metas$Sevs[ll_use], level = c("WNA", "WNE", "WNF"))
allfactors = gather(allfactors, key = "factor", value = "score", -time, -symptom)
allfactors =  allfactors[ allfactors$factor%in%c("Factor1"),]
# WNA, WNE, WNF
pdf(paste0("boxplotF1_",analysis_date,".pdf"), height = 4, width = 4)
my.color = c(brewer.pal(5, 'Blues')[2], brewer.pal(5, 'Reds')[3], brewer.pal(5, 'Oranges')[3])

ggplot(allfactors, aes(x=time, y = score,colour=symptom)) + geom_boxplot()+scale_color_manual(values=my.color)+geom_point(size=1.2, position=position_jitterdodge(dodge.width = 0.75,jitter.width=0.05, seed=1))+facet_wrap(~factor)
dev.off()
```


```
library(tidyr)
allfactors = factors_joint[[1]][,1:factor_rank]
allfactors=normalization(allfactors, method = "quantile")
colnames(allfactors)=paste0('Factor', c(1:ncol(allfactors)))
allfactors= data.frame(allfactors)
allfactors =allfactors[ll_use,]
allfactors$time = metas$Times[ll_use]
allfactors$symptom = factor(metas$Sevs[ll_use], level = c("WNA", "WNE", "WNF"))
allfactors = gather(allfactors, key = "factor", value = "score", -time, -symptom)
allfactors =  allfactors[ allfactors$factor%in%c("Factor5"),]
# WNA, WNE, WNF

my.color = c(brewer.pal(5, 'Blues')[2], brewer.pal(5, 'Reds')[3], brewer.pal(5, 'Oranges')[3])

gplot1 = ggplot(allfactors, aes(x=time, y = score,colour=symptom)) + geom_boxplot()+scale_color_manual(values=my.color)+geom_point(size=1.2, position=position_jitterdodge(dodge.width = 0.75,jitter.width=0.05, seed=1))+facet_wrap(~factor)


allfactors = factors_joint[[1]][,c(1:factor_rank)]
allfactors=normalization(allfactors, method = "quantile")
aa = intersect(metas$SubjectID[metas$Times == "T1"],
               metas$SubjectID[metas$Times == "T2"])
ll1 = c()
ll2 = c()
for(i in 1:length(aa)){
  ll1 = c(ll1, which(metas$Times == "T1" & (metas$SubjectID==aa[i])))
  ll2 = c(ll2, which(metas$Times == "T2" & (metas$SubjectID==aa[i])))
}
x1 = allfactors[ll1,]
x2 = allfactors[ll2,]
yy = metas$Sev[ll1]

paired_data = data.frame(T1=x1[,5], T2 = x2[,5], symptom = yy)
gplot2 = ggpubr::ggpaired(data=paired_data, cond1 = "T1", cond2 = "T2", facet.by = "symptom", color ="symptom")+scale_color_manual(values=my.color)
pdf(paste0("boxplotF5_",analysis_date,".pdf"), height = 4, width = 8)
print(ggpubr::ggarrange(gplot1, gplot2,  ncol = 2))
dev.off()
```

### Feature weights


```
view_names = names(data_mofa)
pdf(paste0("joint_correlation_factor",1,analysis_date,".pdf"), height = 12, width = 10)
for(k in 1:length(view_names)){
  print(k)
  print(plot_data_scatter(MOFAobject_joint_all, 
  view = view_names[k], 
  factor = 1, 
  features = min(ncol(data_processed[[k]]),20),
  add_lm = T,
  dot_size = 1,
  color_by = "Sev",
  shape_by = "Time"))
}
dev.off()

pdf(paste0("joint_correlation_factor",5,analysis_date,".pdf"), height = 12, width = 10)

for(k in 1:length(view_names)){
  print(k)
  print(plot_data_scatter(MOFAobject_joint_all, 
  view = view_names[k], 
  factor = 5, 
  features = min(ncol(data_processed[[k]]),20),
  add_lm = T,
  dot_size = 1,
  color_by = "Sev",
  shape_by = "Time"))
}
dev.off()


pdf(paste0("umap_joint",analysis_date,".pdf"))
mofa_umap <- run_umap(MOFAobject_joint_all, n_neighbors = 7, min_dist = 0.01)

plot_dimred(mofa_umap,
  method = "UMAP",  # method can be either "TSNE" or "UMAP"
  color_by = "Sev",
  shape_by = "Time",
  dot_size = 5
)
dev.off()
```


```
weights <- get_weights(MOFAobject_joint_all, views = "all", factors = "all")
factor_feature_correlations = list()
for(k in 1:ncol(allfactors)){
  print(k)
  factor_feature_correlations[[k]] = list()
  for(d in 1:length(data_mofa)){
    X =data_mofa[[d]]
    factor_feature_correlations[[k]][[d]] = data.frame(matrix(0, ncol = 2, nrow = ncol(X)))
    rownames(factor_feature_correlations[[k]][[d]]) = colnames(X)
    for(j in 1:ncol(X)){
      tmp = cor.test(X[,j], factors_joint[[1]][,k],use = "pairwise.complete")
      factor_feature_correlations[[k]][[d]][j,1]=tmp$estimate
      factor_feature_correlations[[k]][[d]][j,2]=tmp$p.value
    }
  }
}

for(k in c(1,5)){
  all_features = list()
  for(d in 1:length(factor_feature_correlations[[k]])){
    all_features[[d]] = factor_feature_correlations[[k]][[d]]
    all_features[[d]]$mofa_weigths = weights[[d]][,k]
    colnames(all_features[[d]]) = c("correlation", "correlation.pval", "mofa.weights")
    all_features[[d]] = all_features[[d]][,c("mofa.weights", "correlation", "correlation.pval")]
    tmp = rownames(all_features[[d]])
    tmp =  sapply(strsplit(tmp,":"), function(z) z[[2]])
    rownames(all_features[[d]]) = tmp
  }
  names(all_features) = names(data_mofa)
  saveRDS(all_features, file = paste0("feature_importance_factor",k,analysis_date,".RDS"))
}
```
