## Supplemental Software for "Early cellular and molecular signatures correlate with severity of West Nile Virus infection": README.txt

platform: x86_64-pc-linux-gnu
R version: R version 3.6.1 (2019-07-05)

Required R packages:
readr (1.3.1)
reticulate (1.22)
MOFA2 (1.1.7)
ggplot2 (3.3.3)
tidyr (1.1.3)

Install instruction for MOFA2: https://biofam.github.io/MOFA2/installation.html

This scripts use existing packages to perform multi-omics analysis. There is no demo for using these packages, but we have one version of data after basic preprocessing and filtering (processed_filtered_data.RData), and you can use it as a start and run the  markdown file chunk by chunk to reproduce results in the paper (if r packages are correctly installed, ~1hr).
