## Supplemental Software for "Early cellular and molecular signatures correlate with severity of West Nile Virus infection": code.wgcna.groups.demo.R

```
##### WGCNA-based differential module analysis - Demo #####

# Adapted from Kazer et al., Nature Medicine 2020,
# "Integrated single-cell analysis of multicellular immune dynamics during hyperacute HIV-1 infection"
# https://pubmed.ncbi.nlm.nih.gov/32251406/

INPUT.DATA.DEMO = 'cellgroups.bycelltype.seuratobj.d211015.sub1.rdata'

{
  library(Seurat)
  library(WGCNA)
  library(flashClust)
  library(Hmisc)
  library(dplyr)
  library(openxlsx)
  library(ggplot2)
  library(gplots)
  library(RColorBrewer)
  library(cowplot)
  library(viridis)
  library(preprocessCore)
  library(reshape2)
  library(data.table)
  library(stringr)
}


##########################################################################################

##### Functions for WGCNA-based module analysis #####

##########################################################################################

# Choosing the appropriate soft-threshold power for generating the adjacency matrix.
FindPower = function(datExpr) {
  powers = c(c(1:10), seq(from = 12, to = 20, by = 2))
  sft = pickSoftThreshold(datExpr, dataIsExpr = T, powerVector = powers, corFnc = cor,
                          corOptions = list(use = 'p', method = 'pearson'), networkType = 'signed')
  
  # Plot the results
  par(mfrow = c(1, 2))
  cex1 = 0.9
  
  # Scale-free topology fit index as a function of the soft-thresholding power
  plot(sft$fitIndices[, 1], -sign(sft$fitIndices[, 3]) * sft$fitIndices[, 2], 
       xlab = 'Soft Threshold (power)', 
       ylab = 'Scale Free Topology Model Fit, signed R^2', 
       type = 'n', main = paste('Scale independence'))
  text(sft$fitIndices[, 1], -sign(sft$fitIndices[, 3]) * sft$fitIndices[, 2], labels = powers, cex = cex1, col = 'red')
  abline(h = 0.80, col = 'red') # Red line corresponds to using an R^2 cut-off
  
  # Mean connectivity as a function of the soft-thresholding power
  plot(sft$fitIndices[, 1], sft$fitIndices[, 5], 
       xlab = 'Soft Threshold (power)', ylab = 'Mean Connectivity', 
       type = 'n', main = paste('Mean connectivity'))
  text(sft$fitIndices[, 1], sft$fitIndices[, 5], labels = powers, cex = cex1, col = 'red')
}


# Generating the adjacency matrix and performing clustering
ClusterTOM = function(datExpr, softPower) {
  adj = adjacency(datExpr, type = 'signed', power = softPower)
  
  # Turn adjacency matrix into a topological overlap matrix (TOM) to minimize the effects of noise and spurious associations.
  TOM = TOMsimilarityFromExpr(datExpr, networkType = 'signed', TOMType = 'signed', power = softPower, corType = 'bicor');
  
  colnames(TOM) = rownames(TOM) = colnames(datExpr)
  dissTOM = 1 - TOM
  
  # Hierarchical clustering of the genes based on the TOM dissimilarity measure
  dist.dissTOM = as.dist(dissTOM)
  dist.dissTOM[is.na(dist.dissTOM)] = 0
  dist.dissTOM[is.nan(dist.dissTOM)] = 0
  geneTree = flashClust(dist.dissTOM, method = 'complete')
  
  # Plot the resulting clustering tree (dendrogram)
  plot(geneTree, xlab = '', sub = '', cex = 0.3)
  
  # returns a list with dissimilarity TOM, and the clustered gene tree.
  return(list(dissTOM = dissTOM, geneTree = geneTree))
}


# Cut clustering dendrogram using 'tree' method (keep minimum module size at 10 genes)
# Cut the resulting clustering dendrogram using the 'tree' method for cutreeDynamic. Minimum module size can be specified.
CutTOMTree = function(datExpr, dissTOM, geneTree, minModuleSize = 10) {
  # Module identification using dynamic tree cut, you can also choose the hybrid method
  dynamicMods = cutreeDynamic(dendro = geneTree,  method = 'tree', minClusterSize = minModuleSize)
  
  # Get the module labels and the size of each module. Lable 0 is reserved for unassigned genes
  print(table(dynamicMods))
  
  # Plot the module assignment under the dendrogram; note: The grey color is reserved for unassigned genes
  dynamicColors = labels2colors(dynamicMods)
  table(dynamicColors)
  plotDendroAndColors(geneTree, dynamicColors, 'Dynamic Tree Cut', 
                      hang = 0.03, guideHang = 0.05, 
                      dendroLabels = F, addGuide = T, 
                      main = 'Gene dendrogram and module colors')
  
  # Set the diagonal of the dissimilarity to NA 
  diag(dissTOM) = NA
  
  #extract modules
  module_colors= setdiff(unique(dynamicColors), 'grey')
  modules = lapply(module_colors, function(x) {colnames(datExpr)[which(dynamicColors==x)]})
  names(modules) = module_colors
  
  #returns list with module colors, and the modules themselves
  return(list(dyanmicColors = dynamicColors, modules = modules))
}

# Merge similar modules
# Merge modules with low dissimilarity. Cutoff for dissimilarity merge can be specified
MergeSimilarModules = function(datExpr, dynamicColors, geneTree, MEDissThres = 0.5) {
  # Cacluate eigengenes
  MEList = moduleEigengenes(datExpr, colors=dynamicColors)
  MEs = MEList$eigengenes
  
  # Calculate dissimilarity of module eigengenes
  MEDiss = 1 - cor(MEs);
  
  # Cluster module eigengenes
  METree = hclust(as.dist(MEDiss), method = 'average');
  
  # Plot the result
  options(repr.plot.width = 12, repr.plot.height = 6)
  plot(METree, main = 'Clustering of module eigengenes', xlab = '', sub = '')
  abline(h = MEDissThres, lwd = 2, col = 'red')
  
  # Call an automatic merging function
  merge = mergeCloseModules(datExpr, dynamicColors, cutHeight = MEDissThres, verbose = 3)
  
  # Plot showing how merged modules exist
  options(repr.plot.width = 12, repr.plot.height = 6)
  plotDendroAndColors(geneTree, cbind(dynamicColors, merge$colors),
                      c('Dynamic Tree Cut', 'Merged dynamic'),
                      hang = 0.03, guideHang = 0.05,
                      dendroLabels = F, addGuide = T)
  
  #extract merged modules
  merged_module_colors= setdiff(unique(merge$colors), 'grey')
  merged_modules = lapply(merged_module_colors, function(x){colnames(datExpr)[which(merge$colors == x)]})
  names(merged_modules) = merged_module_colors
  
  return(list(mergedResults = merge, 
              mergedColors = merge$colors, 
              merged_modules = merged_modules))
}


## Compare modules against randomly generated background modules
## Functions for assaying module significance against background and temporal variability in module score ##

## Test to determine if the genes within the module are truly the least dissimilar compared to randomly generated modules of the same size.
TestModuleSignificance = function(mod, dissTOM, expr.data, n_perm = 10000, pval = 0.05, n.bin = 10) {
  #vectorize the actual distribution of (dis)similarities, and remove zeros!
  true.diss = as.vector(dissTOM[mod, mod])
  true.diss = true.diss[true.diss != 0]
  
  # Bin all genes by expression
  expr.avg = sort(rowMeans(expr.data))
  expr.avg.cut = as.numeric(cut2(x = expr.avg, m = round(length(expr.avg) / n.bin)))
  names(expr.avg.cut) = names(expr.avg)
  
  # Create a table of binnings of all genes and of our module genes
  all.bin.table = table(expr.avg.cut)
  mod.bin.table = table(expr.avg.cut[mod])
  
  # Randomly generate module with same expression binning structure and run t.test and record results
  test.results = data.frame(statistic = rep(NA, n_perm), pval = rep(NA, n_perm)) #container for results
  
  for (i in 1:n_perm){ #by permutation
    random.mod = list() #create an empty list we will fill with gene names (each element will be gene names by bin)
    
    # For each bin of the mod bin table, randomly select that number of genes from the full set with that expression
    for (j in names(mod.bin.table)){ 
      bin.genes = sample(names(expr.avg.cut)[which(expr.avg.cut == as.numeric(j))], mod.bin.table[j], replace = F)
      random.mod[[as.numeric(j)]] = bin.genes #stick those genes into the random.mod list
    }
    
    # Unlist and vectorize the distribution of dissimilarities (remember to remove zeros)
    random.mod = unlist(random.mod)
    random.diss = as.vector(dissTOM[random.mod, random.mod])
    random.diss = random.diss[random.diss != 0]
    
    # Perform a one-sided wilcoxon test and record the statistic and p-value.
    # Note, IMPORTANT: here we perform the test asking if the true diss is LESS THAN the random diss, 
    # as we are trying to minimize dissimilarity
    test = wilcox.test(x = true.diss, y = random.diss, alternative = 'less')
    test.results[i,] = c(test$statistic, test$p.value)
  }
  
  # Correct for multiple hypothesis testing, and then report the proportion of bad tests
  test.results$FDR = p.adjust(test.results$pval, method = 'fdr')
  num.failed.tests = sum(test.results$FDR > pval)
  print(paste(paste0(num.failed.tests, '/', n_perm), ' tests with FDR > 0.05 by the Wilcoxon rank sum test.', sep = ''))
  
  # Is the percentage of failed tests less than or equal to the p-val specified?
  return(num.failed.tests / n_perm <= pval) #returns a vector of booleans indicating if each module was significant based on the specific pval
}


## Test time-point and disease-state variation
## Test for differences in module scores for each group pair by MW or KS
TestModuleStateVariation = function(meta.data) {
    test.pairs = rbind(c('T1_WNA', 'T1_WNE'), c('T1_WNA', 'T1_WNF'), c('T1_WNE', 'T1_WNF'), 
                       c('T2_WNA', 'T2_WNE'), c('T2_WNA', 'T2_WNF'), c('T2_WNE', 'T2_WNF'), 
                       c('T3_WNA', 'T3_WNE'), c('T3_WNA', 'T3_WNF'), c('T3_WNE', 'T3_WNF'), 
                       c('T1_WNA', 'T2_WNA'), c('T1_WNA', 'T3_WNA'), c('T2_WNA', 'T3_WNA'), 
                       c('T1_WNE', 'T2_WNE'), c('T1_WNE', 'T3_WNE'), c('T2_WNE', 'T3_WNE'), 
                       c('T1_WNF', 'T2_WNF'), c('T1_WNF', 'T3_WNF'), c('T2_WNF', 'T3_WNF'))

    mod.pvals = apply(data.frame(meta.data[, grep('^MOD', colnames(meta.data), value = T)]), 2, function(mod) {
        # Get the scores for this module separated by group
        scores.by.grp = split(mod - min(mod), meta.data$state_timepoint)
        scores.by.grp = scores.by.grp[sapply(scores.by.grp, length) >= 10]
        
        # Group size (#cells)
        grsz = t(apply(test.pairs, 1, function(x) 
                        if ( all(x %in% names(scores.by.grp)) ) {
                            c(length(scores.by.grp[[x[1]]]), length(scores.by.grp[[x[2]]]))
                        } else c(NA, NA)
                      ))
        
        # Wilcoxon and KS tests for the predefined group pairs
        wc.p = apply(test.pairs, 1, function(x) 
                        if ( all(x %in% names(scores.by.grp)) ) {
                            wilcox.test(scores.by.grp[[x[1]]], scores.by.grp[[x[2]]])$p.value
                        } else NA
                     )

        ks.p = apply(test.pairs, 1, function(x) 
                if ( all(x %in% names(scores.by.grp)) ) {
                    ks.test(scores.by.grp[[x[1]]], scores.by.grp[[x[2]]])$p.value
                } else NA
             )

        # Absolute fold change
        fcab = t(apply(test.pairs, 1, function(x) 
                        if ( all(x %in% names(scores.by.grp)) ) {
                            tmp.med1 = median(scores.by.grp[[x[1]]])
                            tmp.med2 = median(scores.by.grp[[x[2]]])
                            tmp.fcab = 2 ^ abs(log2(tmp.med2 / tmp.med1))
                            c(tmp.med1, tmp.med2, tmp.fcab)
                        } else rep(NA, 3)
                      ))
        
        test.out = cbind(test.pairs, grsz, wc.p, ks.p, fcab)
        colnames(test.out) = c('Group1', 'Group2', 'Size1', 'Size2', 
                               'PvalWC', 'PvalKS', 'Med1', 'Med2', 'AbsFC')
        data.frame(test.out)
    })
    
    if ( length(mod.pvals) == 1 ) names(mod.pvals) = 'MOD1'
    
    return(mod.pvals)
}


# Function mainly to calculate PCs, but also gives some other outputs
prep.for.cluster = function(ob) {
    ob = FindVariableFeatures(ob, selection.method = 'vst', nfeatures = 3000)
    
    all.genes = rownames(ob) 
    ob = ScaleData(ob, features = all.genes) 
    
    ob = RunPCA(ob, features = VariableFeatures(object = ob))
    
    print(DimPlot(ob, reduction = 'pca') + NoLegend())
    print(ElbowPlot(ob, ndims = 50))
    print((((ob@reductions$pca@stdev)^2 / sum((ob@reductions$pca@stdev)^2)) * 100))
    
    return(ob)
}


# Function completely re-written to work with seurat v3 that returns top genes based on PC loadings
## Function completely re-written to work with seurat v3 that returns top genes based on PC loadings (positive and negative)
## Inputs:
# object = cell type object
# npcs = number of PCs to use
# num.genes = number of top genes to return

PCTop = function(object, npcs, num.genes) {
    num.genes = num.genes/2 #we only want have the num.genes for positive and negative loadings
    pcs = object@reductions$pca@feature.loadings #get matrix of gene loadings for PCs
    
    gene_vector = c()
    for (i in 1:npcs) { #iterate through desired PCs
        pc_header = colnames(pcs)[i] #get genes for one PC
        new_PC1 = pcs[order(pcs[,pc_header]), ] #order those genes by their loadings for the desired PC
        top = rownames(new_PC1)[1:num.genes] #get the highest loading genes
        bottom = rownames(new_PC1)[(3000-(num.genes-1)):3000] #get highest negative loading genes
        gene_vector = c(gene_vector, top, bottom) #combine those genes with the set I already have
    }
    
    return(gene_vector)
}


###################################################################################################

##### Case study of plasma B cells: WGCNA-based differential modules #####

###################################################################################################

(group.list = get(load(INPUT.DATA.DEMO)))

(grp = names(group.list))
# [1] "Plasma_B_Cells"
(S.cell_object = group.list[[grp]])
# An object of class Seurat 
# 37919 features across 118 samples within 1 assay 
# Active assay: RNA (37919 features, 3000 variable features)
# 2 dimensional reductions calculated: pca, umap

table(S.cell_object@meta.data$state_timepoint)

#choose the number of PCs we want (I choose npc by a 75% variance explained threshold)
pca_explained = ((S.cell_object@reductions$pca@stdev)^2 / sum((S.cell_object@reductions$pca@stdev)^2))
pc_use = 1
while(sum(pca_explained[1:pc_use]) < .75) {
  pc_use = pc_use + 1
}

message('#PCs to use: ', pc_use)
#PCs to use: 34

# Get the expression matrix for the top genes contributing to significant structure in PC-space
S.cell_object = ProjectDim(S.cell_object)

cell_object.top.genes = unique(PCTop(S.cell_object, npcs = pc_use, num.genes = 50))
length(cell_object.top.genes)
# [1] 1188

cell_object.mat.top.genes = as.matrix(S.cell_object@assays$RNA@data[cell_object.top.genes, ])
dim(cell_object.mat.top.genes)
# [1] 1188  118

# Look at the structure of the Scale Free Topology and Mean Connectivity as a function of Soft Threshold (Power)
options(repr.plot.width = 12, repr.plot.height = 6)
FindPower(datExpr = t(cell_object.mat.top.genes))

# Calculate the TOM, cluster the genes, and cut the tree based on desired module parameters
cell_object.ClusterTOM = ClusterTOM(datExpr = t(cell_object.mat.top.genes), softPower = 5)
dim(cell_object.ClusterTOM$dissTOM)
# [1] 1188 1188

cell_object.ColorsModules = CutTOMTree(datExpr = t(cell_object.mat.top.genes), 
                                       dissTOM = cell_object.ClusterTOM$dissTOM,
                                       geneTree = cell_object.ClusterTOM$geneTree, 
                                       minModuleSize = 10)

# Merge modules with disimilarity < 0.5
if ( length(cell_object.ColorsModules$modules) > 1 ) {
  cell_object.MergedModules = MergeSimilarModules(datExpr = t(cell_object.mat.top.genes), 
                                                  dynamicColors = cell_object.ColorsModules$dyanmicColors,
                                                  geneTree = cell_object.ClusterTOM$geneTree, MEDissThres = 0.5)        
} else {
  cell_object.MergedModules = list(mergedColors = cell_object.ColorsModules$dyanmicColors, 
                                   merged_modules = cell_object.ColorsModules$modules)
  
  plotDendroAndColors(cell_object.ClusterTOM$geneTree, 
                      cbind(cell_object.ColorsModules$dyanmicColors, cell_object.ColorsModules$dyanmicColors),
                      c('Dynamic Tree Cut', 'Merged dynamic'),
                      dendroLabels = F, hang = 0.03,
                      addGuide = T, guideHang = 0.05)
}

# Test for module significance against randomly generated modules
system.time({
  cell_object.Modules.isSig = sapply(cell_object.MergedModules$merged_modules, function(module) {
    TestModuleSignificance(mod = module, 
                           dissTOM = cell_object.ClusterTOM$dissTOM, 
                           expr.data = cell_object.mat.top.genes,
                           n_perm = 1000, pval = 0.05, n.bin = 10) #nperm = 10000 for full analysis
  })
})
#   user  system elapsed 
# 95.345   5.743 101.876

cell_object.Modules.isSig
sum(cell_object.Modules.isSig)

### Significant modules ###
cell_object.Sig.Modules = cell_object.MergedModules$merged_modules[cell_object.Modules.isSig]
names(cell_object.Sig.Modules) = paste0('MOD', 1:length(cell_object.Sig.Modules))
length(cell_object.Sig.Modules)
#[1] 32

# Score each cell for each module using Seurat's built-in AddModuleScore
(S.cell_object = AddModuleScore(S.cell_object, features = cell_object.Sig.Modules, ctrl.size = 5, name = 'MOD'))

dim(S.cell_object@meta.data)
head(S.cell_object@meta.data)

tmp.metadt = S.cell_object@meta.data[, c('state_timepoint', names(cell_object.Sig.Modules))]

wgcna.signi.mods = list(M = cell_object.Sig.Modules, S = tmp.metadt)


# Visualize scores by box & whisker
cell_object.gene.clusters = data.frame(S.cell_object@meta.data)[grep('MOD', colnames(S.cell_object@meta.data))]

par(omi = c(0, 0, 0, 0), mfrow = c(1, 1), mar = c(5, 4, 3, 1), cex = 1.3)
boxplot(cell_object.gene.clusters, las = 2, ylab = 'Module score')

# Look at how many cells are present at each time point to choose the appropriate sample size and time points 
table(S.cell_object@meta.data$state_timepoint)
# T1_WNA T1_WNE T1_WNF T2_WNA T2_WNE T2_WNF T3_WNA T3_WNE T3_WNF 
#     71      1      5     11     10     10      9      0      1

## Test for differential module scores for each group pair using MW and KS
cell_object.clust.tests = TestModuleStateVariation(meta.data = S.cell_object@meta.data)
length(cell_object.clust.tests)
sapply(cell_object.clust.tests, dim)

cell_object.clust.tests.table = NULL
for ( i in 1:length(cell_object.clust.tests) ) {
  cell_object.clust.tests.table = rbind(cell_object.clust.tests.table, 
                                        cbind(Module = names(cell_object.clust.tests)[i], 
                                              cell_object.clust.tests[[i]]))
}

cell_object.clust.tests.table = cell_object.clust.tests.table[apply(cell_object.clust.tests.table, 1, 
                                                                    function(x) all(!is.na(x))), ]

## Conditions for differential modules
cond.diffmod = sapply(cell_object.clust.tests, function(x) 
  sum(apply(x, 1, function(y) mean(as.numeric(y[c('PvalWC', 'PvalKS')])) < 0.05 & y['AbsFC'] > 1.3), na.rm = T))
sum(cond.diffmod > 0)
# [1] 11

## Keep differential modules with the minimum pval < 0.05 and absFC > 1.3
(cell_object.Sig.Var.Modules = cell_object.Sig.Modules[cond.diffmod > 0])
length(cell_object.Sig.Var.Modules)
# [1] 11

# remove the original module scores, and add scores for only those modules that were variant in time
S.cell_object@meta.data = S.cell_object@meta.data[, -grep('MOD', colnames(S.cell_object@meta.data))]
S.cell_object = AddModuleScore(S.cell_object, features = cell_object.Sig.Var.Modules, ctrl.size = 5, name = 'MOD')
(cell_object.gene.clusters.names.final = grep('MOD', colnames(S.cell_object@meta.data), value = T))
dim(S.cell_object@meta.data)
head(S.cell_object@meta.data)

# rename the modules with the appropriate naming scheme (e.g. MOD1)
names(cell_object.Sig.Var.Modules) = cell_object.gene.clusters.names.final
if ( length(cell_object.Sig.Var.Modules) > 1 ) {
  easy_mod = t(plyr::ldply(cell_object.Sig.Var.Modules, rbind))[-1, ]
} else {
  easy_mod = data.matrix(cell_object.Sig.Var.Modules[[1]])
}
colnames(easy_mod) = cell_object.gene.clusters.names.final
dim(easy_mod)
head(easy_mod)

# Plot box & whiskers of final modules
tmp1 = table(S.cell_object@meta.data$state_timepoint)
tmp2 = names(which(tmp1 >= 10))
tmp.data = S.cell_object@meta.data[S.cell_object@meta.data$state_timepoint %in% tmp2, ]

cell_object.geneclusts.time.final = lapply(cell_object.gene.clusters.names.final, function(c_name) {
  p = ggplot(tmp.data, aes_string(x = 'state_timepoint', y = c_name, fill = 'state_timepoint')) +
    geom_boxplot() + scale_fill_manual(values = rainbow(nlevels(tmp.data$state_timepoint))) + xlab('') + 
    theme(legend.position = 'none', axis.text = element_text(size = 16), axis.text.x = element_text(angle = 90), 
          axis.title = element_text(size = 16))
  return(p)
})

plot_grid(plotlist = cell_object.geneclusts.time.final, ncol = 4)


### Number of significant modules with different p-value thresholds and the maximum FC > 1.3 ###
{
  thres.range = c(seq(0.001, 0.009, 0.001), seq(0.01, 0.1, 0.01))
  sigmods.thres = list()
  for ( THRES in thres.range ) {
    tmp.sigmods = lapply(cell_object.clust.tests, function(x) {
      z = t(apply(x, 1, function(y) 
        c(State = paste0(y['Group1'], '|', y['Group2']), 
          Cond = (mean(as.numeric(y[c('PvalWC', 'PvalKS')])) < THRES & 
                    y['AbsFC'] > 1.3))))
      z = z[apply(z, 1, function(j) j[2] == 'TRUE'), 'State']
      z[!is.na(z)]
    })
    
    tmp.sigmods = tmp.sigmods[sapply(tmp.sigmods, length) > 0]
    sigmods.thres[[as.character(THRES)]] = tmp.sigmods
  }
  
  sapply(sigmods.thres, length)
}

### Differential modules by definition with p = 0.01
diffmods = sigmods.thres$`0.01`
length(diffmods)
#[1] 7


#########################################################################################

##### Correlation between module scores and gene expression #####

#########################################################################################

tmp.diffmods = wgcna.signi.mods$M[names(diffmods)]
tmp.diffmods = tmp.diffmods[!sapply(tmp.diffmods, is.null)]

tmp.metadt = wgcna.signi.mods$S[, c('state_timepoint', names(tmp.diffmods))]
tmp.metadt = cbind(patient = as.character(group.list[[grp]]@meta.data[, 'U19.ID']), tmp.metadt)

names(tmp.diffmods) = paste0('MOD', 1:length(tmp.diffmods))
colnames(tmp.metadt)[grep('MOD', colnames(tmp.metadt))] = names(tmp.diffmods)

tmp.expr = t(data.frame(group.list[[grp]]@assays$RNA@data)[unlist(tmp.diffmods), rownames(tmp.metadt)])
tmp.metadt = cbind(tmp.metadt, tmp.expr)

tmp1 = table(tmp.metadt$state_timepoint)
tmp2 = names(which(tmp1 >= 10))

expr.mtx = tmp.metadt[tmp.metadt$state_timepoint %in% tmp2, ]

mods.genes.corr = lapply(tmp.diffmods, as.null)
for ( i in 1:length(tmp.diffmods) ) {
  tmp.data = expr.mtx[, c(names(tmp.diffmods)[i], tmp.diffmods[[i]])]
  mods.genes.corr[[i]] = sort(apply(tmp.data[, -1], 2, function(x) cor(tmp.data[, 1], x)), decreasing = T)
}

sort(sapply(mods.genes.corr, mean))
#     MOD1      MOD4      MOD6      MOD2      MOD5      MOD7      MOD3 
#0.4311331 0.4469708 0.4618680 0.5559188 0.6590177 0.7178579 0.7246333


##### END OF DEMO #####
```
