## Supplemental Software for "Early cellular and molecular signatures correlate with severity of West Nile Virus infection": code.wgcna.groups_copy.R

```
system('R --version', intern=T)
#[1] "R version 4.0.5 (2021-03-31) -- \"Shake and Throw\""

##### WGCNA-based differential module analysis #####
# Adapted from the supplementary code of Kazer et al.,
# "An Integrated Single-Cell Analysis of Multicellular Immune Dynamics during Hyper-Acute HIV-1 #Infection."

{
  library(WGCNA)
  library(flashClust)
  library(Hmisc)
  library(dplyr)
  library(openxlsx)
  library(ggplot2)
  library(gplots)
  library(cowplot)
  library(viridis)
  library(impute)
  library(preprocessCore)
  library(Seurat)
  library(reshape2)
  library(data.table)
  library(stringr)
}


##########################################################################################

##### Functions for WGCNA-based module analysis #####

##########################################################################################

# Choosing the appropriate soft-threshold power for generating the adjacency matrix.
FindPower = function(datExpr) {
  powers = c(c(1:10), seq(from = 12, to = 20, by = 2))
  sft = pickSoftThreshold(datExpr, dataIsExpr = T, powerVector = powers, corFnc = cor,
                          corOptions = list(use = 'p', method = 'pearson'), networkType = 'signed')
  
  # Plot the results
  par(mfrow = c(1, 2))
  cex1 = 0.9
  
  # Scale-free topology fit index as a function of the soft-thresholding power
  plot(sft$fitIndices[, 1], -sign(sft$fitIndices[, 3]) * sft$fitIndices[, 2], 
       xlab = 'Soft Threshold (power)', 
       ylab = 'Scale Free Topology Model Fit, signed R^2', 
       type = 'n', main = paste('Scale independence'))
  text(sft$fitIndices[, 1], -sign(sft$fitIndices[, 3]) * sft$fitIndices[, 2], labels = powers, cex = cex1, col = 'red')
  abline(h = 0.80, col = 'red')
  
  # Mean connectivity as a function of the soft-thresholding power
  plot(sft$fitIndices[, 1], sft$fitIndices[, 5], 
       xlab = 'Soft Threshold (power)', ylab = 'Mean Connectivity', 
       type = 'n', main = paste('Mean connectivity'))
  text(sft$fitIndices[, 1], sft$fitIndices[, 5], labels = powers, cex = cex1, col = 'red')
}


# Generating the adjacency matrix and performing clustering
ClusterTOM = function(datExpr, softPower) {
  adj= adjacency(datExpr,type = 'signed', power = softPower)
  
  # Turn adjacency matrix into a topological overlap matrix (TOM) to minimize the effects of noise and spurious associations.
  TOM = TOMsimilarityFromExpr(datExpr, networkType = 'signed', TOMType = 'signed', power = softPower, corType = 'bicor');
  
  colnames(TOM) = rownames(TOM) = colnames(datExpr)
  dissTOM = 1 - TOM
  
  # Hierarchical clustering of the genes based on the TOM dissimilarity measure
  dist.dissTOM = as.dist(dissTOM)
  dist.dissTOM[is.na(dist.dissTOM)] = 0
  dist.dissTOM[is.nan(dist.dissTOM)] = 0
  geneTree = flashClust(dist.dissTOM, method = 'complete')
  
  # Plot the resulting clustering tree (dendrogram)
  plot(geneTree, xlab = '', sub = '', cex = 0.3)
  
  # returns a list with dissimilarity TOM, and the clustered gene tree.
  return(list(dissTOM = dissTOM, geneTree = geneTree))
}


# Cut clustering dendrogram using 'tree' method (keep minimum module size at 10 genes)
# Cut the resulting clustering dendrogram using the 'tree' method for cutreeDynamic. Minimum module size can be specified.
CutTOMTree = function(datExpr, dissTOM, geneTree, minModuleSize = 10) {
  # Module identification using dynamic tree cut (can also choose the hybrid method)
  dynamicMods = cutreeDynamic(dendro = geneTree,  method = 'tree', minClusterSize = minModuleSize);

  # Get the module labels and the size of each module. Lable 0 is reserved for unassigned genes
  print(table(dynamicMods))
  
  # Plot the module assignment under the dendrogram; note: The grey color is reserved for unassigned genes
  dynamicColors = labels2colors(dynamicMods)
  table(dynamicColors)
  plotDendroAndColors(geneTree, dynamicColors, 'Dynamic Tree Cut', 
                      hang = 0.03, guideHang = 0.05, 
                      dendroLabels = F, addGuide = T, 
                      main = 'Gene dendrogram and module colors')
  
  # Set the diagonal of the dissimilarity to NA 
  diag(dissTOM) = NA
  
  #extract modules
  module_colors= setdiff(unique(dynamicColors), 'grey')
  modules = lapply(module_colors, function(x) {colnames(datExpr)[which(dynamicColors==x)]})
  names(modules) = module_colors

  #returns list with module colors, and the modules themselves
  return(list(dyanmicColors = dynamicColors, modules = modules))
}

# Merge similar modules
# Merge modules with low dissimilarity. Cutoff for dissimilarity merge can be specified
MergeSimilarModules = function(datExpr, dynamicColors, geneTree, MEDissThres = 0.5) {
  # Calculate eigengenes
  MEList = moduleEigengenes(datExpr, colors=dynamicColors)
  MEs = MEList$eigengenes
  
  # Calculate dissimilarity of module eigengenes
  MEDiss = 1 - cor(MEs);
  
  # Cluster module eigengenes
  METree = hclust(as.dist(MEDiss), method = 'average');
  
  # Plot the result
  options(repr.plot.width = 12, repr.plot.height = 6)
  plot(METree, main = 'Clustering of module eigengenes', xlab = '', sub = '')
  abline(h = MEDissThres, lwd = 2, col = 'red')
  
  # Call an automatic merging function
  merge = mergeCloseModules(datExpr, dynamicColors, cutHeight = MEDissThres, verbose = 3)
  
  # Plot showing how merged modules exist
  options(repr.plot.width = 12, repr.plot.height = 6)
  plotDendroAndColors(geneTree, cbind(dynamicColors, merge$colors),
                      c('Dynamic Tree Cut', 'Merged dynamic'),
                      hang = 0.03, guideHang = 0.05,
                      dendroLabels = F, addGuide = T)
  
  #extract merged modules
  merged_module_colors= setdiff(unique(merge$colors), 'grey')
  merged_modules = lapply(merged_module_colors, function(x){colnames(datExpr)[which(merge$colors == x)]})
  names(merged_modules) = merged_module_colors
  
  return(list(mergedResults = merge, 
              mergedColors = merge$colors, 
              merged_modules = merged_modules))
}


## Compare modules against randomly generated background modules
## Functions for assaying module significance against background and temporal variability in module score ##

## Test to determine if the genes within the module are truly the least dissimilar compared to randomly generated modules of the same size.
TestModuleSignificance = function(mod, dissTOM, expr.data, n_perm = 10000, pval = 0.05, n.bin = 10) {
  #vectorize the actual distribution of (dis)similarities, and remove zeros
  true.diss = as.vector(dissTOM[mod, mod])
  true.diss = true.diss[true.diss != 0]
  
  # Bin all genes by expression
  expr.avg = sort(rowMeans(expr.data))
  expr.avg.cut = as.numeric(cut2(x = expr.avg, m = round(length(expr.avg) / n.bin)))
  names(expr.avg.cut) = names(expr.avg)
  
  # Create a table of binnings of all genes and of our module genes
  all.bin.table = table(expr.avg.cut)
  mod.bin.table = table(expr.avg.cut[mod])
  
  # Randomly generate module with same expression binning structure and run t.test and record results
  test.results = data.frame(statistic = rep(NA, n_perm), pval = rep(NA, n_perm)) #container for results
  
  for (i in 1:n_perm){ #by permutation
    random.mod = list() #create an empty list we will fill with gene names (each element will be gene names by bin)
    
    # For each bin of the mod bin table, randomly select that number of genes from the full set with that expression
    for (j in names(mod.bin.table)){ 
      bin.genes = sample(names(expr.avg.cut)[which(expr.avg.cut == as.numeric(j))], mod.bin.table[j], replace = F)
      random.mod[[as.numeric(j)]] = bin.genes #stick those genes into the random.mod list
    }
    
    # Unlist and vectorize the distribution of dissimilarities (remember to remove zeros)
    random.mod = unlist(random.mod)
    random.diss = as.vector(dissTOM[random.mod, random.mod])
    random.diss = random.diss[random.diss != 0]
    
    # Perform a one-sided wilcoxon test and record the statistic and p-value.
    # Note, IMPORTANT: here we perform the test asking if the true diss is LESS THAN the random diss, 
    # as we are trying to minimize dissimilarity
    test = wilcox.test(x = true.diss, y = random.diss, alternative = 'less')
    test.results[i,] = c(test$statistic, test$p.value)
  }
  
  # Correct for multiple hypothesis testing, and then report the proportion of bad tests
  test.results$FDR = p.adjust(test.results$pval, method = 'fdr')
  num.failed.tests = sum(test.results$FDR > pval)
  print(paste(paste0(num.failed.tests, '/', n_perm), ' tests with FDR > 0.05 by the Wilcoxon rank sum test.', sep = ''))
  
  #returns a vector of booleans indicating if each module was significant based on the specific pval
  return(num.failed.tests / n_perm <= pval)
}


## Test time-point and disease-state variation
## Test for differences in module scores for each group pair by MW or KS
TestModuleStateVariation = function(meta.data) {
    test.pairs = rbind(c('T1_WNA', 'T1_WNE'), c('T1_WNA', 'T1_WNF'), c('T1_WNE', 'T1_WNF'), 
                       c('T2_WNA', 'T2_WNE'), c('T2_WNA', 'T2_WNF'), c('T2_WNE', 'T2_WNF'), 
                       c('T3_WNA', 'T3_WNE'), c('T3_WNA', 'T3_WNF'), c('T3_WNE', 'T3_WNF'), 
                       c('T1_WNA', 'T2_WNA'), c('T1_WNA', 'T3_WNA'), c('T2_WNA', 'T3_WNA'), 
                       c('T1_WNE', 'T2_WNE'), c('T1_WNE', 'T3_WNE'), c('T2_WNE', 'T3_WNE'), 
                       c('T1_WNF', 'T2_WNF'), c('T1_WNF', 'T3_WNF'), c('T2_WNF', 'T3_WNF'))

    mod.pvals = apply(data.frame(meta.data[, grep('^MOD', colnames(meta.data), value = T)]), 2, function(mod) {
        # Get the scores for this module separated by group
        scores.by.grp = split(mod - min(mod), meta.data$state_timepoint)
        scores.by.grp = scores.by.grp[sapply(scores.by.grp, length) >= 10]
        
        # Group size (#cells)
        grsz = t(apply(test.pairs, 1, function(x) 
                        if ( all(x %in% names(scores.by.grp)) ) {
                            c(length(scores.by.grp[[x[1]]]), length(scores.by.grp[[x[2]]]))
                        } else c(NA, NA)
                      ))
        
        # Wilcoxon and KS tests for the predefined group pairs
        wc.p = apply(test.pairs, 1, function(x) 
                        if ( all(x %in% names(scores.by.grp)) ) {
                            wilcox.test(scores.by.grp[[x[1]]], scores.by.grp[[x[2]]])$p.value
                        } else NA
                     )

        ks.p = apply(test.pairs, 1, function(x) 
                if ( all(x %in% names(scores.by.grp)) ) {
                    ks.test(scores.by.grp[[x[1]]], scores.by.grp[[x[2]]])$p.value
                } else NA
             )

        # Absolute fold change
        fcab = t(apply(test.pairs, 1, function(x) 
                        if ( all(x %in% names(scores.by.grp)) ) {
                            tmp.med1 = median(scores.by.grp[[x[1]]])
                            tmp.med2 = median(scores.by.grp[[x[2]]])
                            tmp.fcab = 2 ^ abs(log2(tmp.med2 / tmp.med1))
                            c(tmp.med1, tmp.med2, tmp.fcab)
                        } else rep(NA, 3)
                      ))
        
        test.out = cbind(test.pairs, grsz, wc.p, ks.p, fcab)
        colnames(test.out) = c('Group1', 'Group2', 'Size1', 'Size2', 
                               'PvalWC', 'PvalKS', 'Med1', 'Med2', 'AbsFC')
        data.frame(test.out)
    })
    
    if ( length(mod.pvals) == 1 ) names(mod.pvals) = 'MOD1'
    
    return(mod.pvals)
}


# Function mainly to calculate PCs, but also gives some other outputs
prep.for.cluster = function(ob) {
    ob = FindVariableFeatures(ob, selection.method = 'vst', nfeatures = 3000)
    
    all.genes = rownames(ob) 
    ob = ScaleData(ob, features = all.genes) 
    
    ob = RunPCA(ob, features = VariableFeatures(object = ob))
    
    print(DimPlot(ob, reduction = 'pca') + NoLegend())
    print(ElbowPlot(ob, ndims = 50))
    print((((ob@reductions$pca@stdev)^2 / sum((ob@reductions$pca@stdev)^2)) * 100))
    
    return(ob)
}


# Function completely re-written to work with seurat v3 that returns top genes based on PC loadings
## Function completely re-written to work with seurat v3 that returns top genes based on PC loadings (positive and negative)
## Inputs:
# object = cell type object
# npcs = number of PCs I want to use
# num.genes = number of top genes I want back per gene

PCTop = function(object, npcs, num.genes) {
    num.genes = num.genes/2 #we only want have the num.genes for positive and negative loadings
    pcs = object@reductions$pca@feature.loadings #get matrix of gene loadings for PCs
    
    gene_vector = c()
    for (i in 1:npcs) { #iterate through desired PCs
        pc_header = colnames(pcs)[i] #get genes for one PC
        new_PC1 = pcs[order(pcs[,pc_header]), ] #order those genes by their loadings for the desired PC
        top = rownames(new_PC1)[1:num.genes] #get the highest loading genes
        bottom = rownames(new_PC1)[(3000-(num.genes-1)):3000] #get highest negative loading genes
        gene_vector = c(gene_vector, top, bottom) #combine those genes with the set I already have
    }
    
    return(gene_vector)
}


##########################################################################################

##### Load the Seurat object #####

##########################################################################################

system.time({
    wnv = get(load('wnv.seqwell.pbmcs.seurat.d211011.Robj'))
    rm(wnv_cleaned)
})
#    user  system elapsed 
# 122.030  10.191 133.958

wnv
# An object of class Seurat 
# 37919 features across 56858 samples within 1 assay 
# Active assay: RNA (37919 features, 3000 variable features)
# 2 dimensional reductions calculated: pca, umap

dim(wnv@meta.data)
# [1] 56858    21

colnames(wnv@meta.data)

wnv@meta.data$new_cell_type = factor(wnv@meta.data$new_cell_type)
levels(wnv@meta.data$new_cell_type)
#[1] "B_Cells"                 "CD14+_Monocytes"         "CD16+_Monocytes"         "CD4+_T_Cells"           
#[5] "CD8+_T_Cells"            "cDCs"                    "NK_Cells"                "pDCs"                   
#[9] "Plasma_B_Cells"          "Proliferating_T_Cells"   "Regulatory_CD4+_T_Cells"
table(wnv@meta.data$new_cell_type)

wnv@meta.data$disease_state = factor(wnv@meta.data$disease_state)
levels(wnv@meta.data$disease_state)
# [1] "WNA" "WNF" "WNE"
table(wnv@meta.data$disease_state)
#   WNA   WNE   WNF 
# 22879 11574 22405

wnv@meta.data$state_timepoint = factor(wnv@meta.data$state_timepoint)
levels(wnv@meta.data$state_timepoint)
# "T1_WNA" "T2_WNA" "T1_WNF" "T2_WNF" "T2_WNE" "T1_WNE" "T3_WNA" "T3_WNF" "T3_WNE"
table(wnv@meta.data$state_timepoint)
#T1_WNA T1_WNE T1_WNF T2_WNA T2_WNE T2_WNF T3_WNA T3_WNE T3_WNF 
# 10907   4019   7904   6167   6578   9250   5805    977   5251


# Split the object by each cell type
system.time({
    group.list = SplitObject(wnv, split.by = 'new_cell_type')
})
#   user  system elapsed 
# 21.375  41.568 113.615

length(group.list)
# [1] 11
sapply(group.list, dim)
all(sapply(group.list, ncol) > 100)
#[1] TRUE
sort(sapply(group.list, ncol), decreasing = T)
group.list

{
  par(mfrow = c(1, 1), omi = c(2.2, 0, 0, 0), mar = c(4, 5, 2.5, 1), mgp = c(3.5, 1, 0), cex = 1.5)
  barplot(sort(sapply(group.list, ncol), decreasing = T), las = 2, 
          ylab = 'Number of cells', 
          main = 'Population size of cell types', cex.main = 1.8)
}

### Run PCA for all groups ###
system.time({
    group.list = lapply(group.list, prep.for.cluster)
})
#    user  system elapsed 
# 698.385  79.145 273.426

group.list = group.list[sort(names(group.list))]
names(group.list)
# [1] "B_Cells"               "CD14+_Monocytes"       "CD16+_Monocytes"       "CD4+_T_Cells"         
# [5] "CD8+_T_Cells"          "cDCs"                  "NK_Cells"              "pDCs"                 
# [9] "Plasma_B_Cells"        "Proliferating_T_Cells" "Regulatory_CD4+_T_Cells"

system.time({
  save(group.list, file = 'cellgroups.bycelltype.seuratobj.d211015.rdata')
})
#    user  system elapsed 
# 637.444  44.070 709.554


##########################################################################################

##### WGCNA significant modules and module scores for all groups #####

##########################################################################################

names(group.list)
# [1] "B_Cells"               "CD14+_Monocytes"       "CD16+_Monocytes"       "CD4+_T_Cells"         
# [5] "CD8+_T_Cells"          "cDCs"                  "NK_Cells"              "pDCs"                 
# [9] "Plasma_B_Cells"        "Proliferating_T_Cells" "Regulatory_CD4+_T_Cells"

system.time({
  wgcna.signi.mods = list()
  
  for ( grp in names(group.list) ) {
    message(grp)
    
    S.cell_object = group.list[[grp]]
    
    # Choose the number of PCs with 75% variance explained
    pca_explained = ((S.cell_object@reductions$pca@stdev)^2 / sum((S.cell_object@reductions$pca@stdev)^2))
    pc_use = 1
    while ( sum(pca_explained[1:pc_use]) < .75 ) {
      pc_use = pc_use + 1
    }
    message('#PCs to use: ', pc_use)
    
    ######################################################
    # Get the expression matrix for the top genes contributing to significant structure in PC-space
    S.cell_object = ProjectDim(S.cell_object)
    cell_object.top.genes = unique(PCTop(S.cell_object, npcs = pc_use, num.genes = 50))
    cell_object.mat.top.genes = as.matrix(S.cell_object@assays$RNA@data[cell_object.top.genes, ])
    
    # Look at the structure of the Scale Free Topology and Mean Connectivity as a function of Soft Threshold (Power)
    options(repr.plot.width = 12, repr.plot.height = 6)
    FindPower(datExpr = t(cell_object.mat.top.genes))
    
    #######################################################
    # Choose softPower from above, here we choose softPower = 5. We recommend playing with the softPower choice
    # to see how it affects the number and size of modules that are generated by the functions below.
    
    # Calculate the TOM, cluster the genes, and cut the tree based on desired module parameters
    cell_object.ClusterTOM = ClusterTOM(datExpr = t(cell_object.mat.top.genes), softPower = 5)
    cell_object.ColorsModules = CutTOMTree(datExpr = t(cell_object.mat.top.genes), 
                                           dissTOM = cell_object.ClusterTOM$dissTOM,
                                           geneTree = cell_object.ClusterTOM$geneTree, 
                                           minModuleSize = 10)
    
    if ( length(cell_object.ColorsModules$modules) == 0 ) {
      cat('\n####################\n')
      next
    }
    
    # Merge modules with disimilarity < 0.5 -- but we can change the threshold too.
    if ( length(cell_object.ColorsModules$modules) > 1 ) {
      cell_object.MergedModules = MergeSimilarModules(datExpr = t(cell_object.mat.top.genes), 
                                                      dynamicColors = cell_object.ColorsModules$dyanmicColors,
                                                      geneTree = cell_object.ClusterTOM$geneTree, MEDissThres = 0.5)        
    } else {
      cell_object.MergedModules = list(mergedColors = cell_object.ColorsModules$dyanmicColors, 
                                       merged_modules = cell_object.ColorsModules$modules)
      
      plotDendroAndColors(cell_object.ClusterTOM$geneTree, 
                          cbind(cell_object.ColorsModules$dyanmicColors, cell_object.ColorsModules$dyanmicColors),
                          c('Dynamic Tree Cut', 'Merged dynamic'),
                          hang = 0.03, guideHang = 0.05, dendroLabels = F, addGuide = T)
      next
    }
    
    #########################################################
    # Test for module significance against randomly generated modules
    cell_object.Modules.isSig = sapply(cell_object.MergedModules$merged_modules, function(module) {
      TestModuleSignificance(mod = module, dissTOM = cell_object.ClusterTOM$dissTOM, expr.data = cell_object.mat.top.genes,
                             n_perm = 10000, pval = 0.05, n.bin = 10)
    })
    
    if ( sum(cell_object.Modules.isSig) == 0 ) {
      cat('\n####################\n')
      next
    }
    
    cell_object.Sig.Modules = cell_object.MergedModules$merged_modules[cell_object.Modules.isSig] # keep those modules that are significant
    names(cell_object.Sig.Modules) = paste0('MOD', 1:length(cell_object.Sig.Modules))
    
    #########################################################
    # Score each cell for each module using Seurat's built-in AddModuleScore
    S.cell_object = AddModuleScore(S.cell_object, features = cell_object.Sig.Modules, ctrl.size = 5, name = 'MOD')
    
    tmp.metadt = S.cell_object@meta.data[, c('state_timepoint', names(cell_object.Sig.Modules))]
    
    wgcna.signi.mods[[grp]] = list(M = cell_object.Sig.Modules, S = tmp.metadt)
  }
  
  save(wgcna.signi.mods, file = 'wgcna.signi.mods.v1.d211015.rdata')
})
#     user   system  elapsed 
# 5029.908  280.567 4994.332

length(wgcna.signi.mods)
#[1] 10

sapply(wgcna.signi.mods, function(x) length(x$M))
sum(sapply(wgcna.signi.mods, function(x) length(x$M)))
#[1] 107

sapply(wgcna.signi.mods, function(x) dim(x$S))
sapply(wgcna.signi.mods, function(x) head(x$S))


##########################################################################################

##### Identification of differential modules for all groups #####

##########################################################################################

names(wgcna.signi.mods)
# [1] "B_Cells"               "CD14+_Monocytes"       "CD16+_Monocytes"       "CD4+_T_Cells"         
# [5] "CD8+_T_Cells"          "cDCs"                  "NK_Cells"              "pDCs"                 
# [9] "Plasma_B_Cells"        "Proliferating_T_Cells" "Regulatory_CD4+_T_Cells"

system.time({
  for ( grp in names(wgcna.signi.mods) ) {
    message(grp)
    
    cell_object.Sig.Modules = wgcna.signi.mods[[grp]]$M
    tmp.metadt = wgcna.signi.mods[[grp]]$S
    
    if ( is.null(tmp.metadt) ) next
    
    # Box & whisker of module scores by module
    cell_object.gene.clusters = data.frame(tmp.metadt)[grep('MOD', colnames(tmp.metadt))]
    
    # Box & whisker of module scores by disease state for each module
    tmp.colors = rainbow(9)[levels(wnv@meta.data$state_timepoint) %in% tmp.metadt$state_timepoint]
    cell_object.gene.clusters.names = grep('MOD', colnames(tmp.metadt), value = T)
    cell_object.geneclusts.time = lapply(cell_object.gene.clusters.names, function(c_name){
      p = ggplot(tmp.metadt, aes_string(x = 'state_timepoint', y = c_name, fill = 'state_timepoint')) +
        geom_boxplot() + scale_fill_manual(values = tmp.colors) + xlab('') + 
        theme(legend.position = 'none', axis.text = element_text(size = 16), axis.title = element_text(size = 16))
      return(p)
    })
    
    ###########################################################
    # Look at how many cells are present at each time point to choose the appropriate sample size and time points 

    # Test for variation in module scores for each symptomatic state vs. the asymptomatic.
    (cell_object.clust.tests = TestModuleStateVariation(meta.data = tmp.metadt))
    
    cell_object.clust.tests.table = NULL
    for ( i in 1:length(cell_object.clust.tests) ) {
      cell_object.clust.tests.table = rbind(cell_object.clust.tests.table, 
                                            cbind(Module = names(cell_object.clust.tests)[i], cell_object.clust.tests[[i]]))
    }
    
    (cell_object.clust.tests.table = cell_object.clust.tests.table[apply(cell_object.clust.tests.table, 1, function(x) all(!is.na(x))), ])
    
    write.csv(cell_object.clust.tests.table, file = paste0(grp, '.diffmod.tests.d211015.csv'), row.names = F)
    
    ## Conditions for differential modules: mean pval < 0.05 and absFC > 1.3
    cond.diffmod = sapply(cell_object.clust.tests, function(x) 
      sum(apply(x, 1, function(y) mean(as.numeric(y[c('PvalWC', 'PvalKS')])) < 0.05 & y['AbsFC'] > 1.3), na.rm = T)
      #sum(apply(x, 1, function(y) min(y[c('PvalWC', 'PvalKS')]) < 0.05 & y['AbsFC'] > 1.3), na.rm = T)
    )
    
    if ( all(cond.diffmod == 0) ) {
      cat('\n####################\n')
      next
    }
    
    ## Keep differential modules with the mean pval < 0.05 and absFC > 1.3
    cell_object.Sig.Var.Modules = cell_object.Sig.Modules[cond.diffmod > 0]
    cat('#Differential modules =', length(cell_object.Sig.Var.Modules), '\n')
    
    ###########################################################
    # Remove the original module scores, and add scores for only those modules that were variant in time
    S.cell_object = group.list[[grp]]
    S.cell_object = AddModuleScore(S.cell_object, features = cell_object.Sig.Var.Modules, ctrl.size = 5, name = 'MOD')
    cell_object.gene.clusters.names.final = grep('MOD', colnames(S.cell_object@meta.data), value = T)
    
    # Table of genes for each module
    names(cell_object.Sig.Var.Modules) = cell_object.gene.clusters.names.final
    if ( length(cell_object.Sig.Var.Modules) > 1 ) {
      easy_mod = t(plyr::ldply(cell_object.Sig.Var.Modules, rbind))[-1, ]
    } else {
      easy_mod = data.matrix(cell_object.Sig.Var.Modules[[1]])
    }
    colnames(easy_mod) = cell_object.gene.clusters.names.final
    
    write.csv(easy_mod, file = paste0(grp, '.final.diffmods.d211015.csv'), row.names = F)
    
    #############################################################
    # Box-whisker plots of the final modules
    tmp1 = table(S.cell_object@meta.data$state_timepoint)
    tmp2 = names(which(tmp1 >= 10))
    tmp.data = S.cell_object@meta.data[S.cell_object@meta.data$state_timepoint %in% tmp2, ]
    
    tmp.colors = rainbow(9)[levels(wnv@meta.data$state_timepoint) %in% tmp.data$state_timepoint]
    cell_object.geneclusts.time.final = lapply(cell_object.gene.clusters.names.final, function(c_name) {
      p = ggplot(tmp.data, aes_string(x = 'state_timepoint', y = c_name, fill = 'state_timepoint')) +
        geom_boxplot() + scale_fill_manual(values = tmp.colors) + xlab('') + 
        theme(legend.position = 'none', axis.text = element_text(size = 16), axis.text.x = element_text(angle = 90), 
              axis.title = element_text(size = 16))
      return(p)
    })
    
    options(repr.plot.width = 6 * length(cell_object.geneclusts.time.final), repr.plot.height = 6)
    print(plot_grid(plotlist = cell_object.geneclusts.time.final, nrow = 1))
    
    ggsave(paste0(grp, '.final.diffmods.d211015.png'), limitsize = F, 
           width = 6 * length(cell_object.geneclusts.time.final), height = 6)
    
    cat('\n####################\n')
  }
})
#  user  system elapsed 
#63.274   8.713  64.879


##########################################################################################

##### Robustness analysis #####

##########################################################################################

{
  output.modtest = list()
  for ( grp in names(wgcna.signi.mods) ) {
    message(grp)
    output.modtest[[grp]] = TestModuleStateVariation(meta.data = wgcna.signi.mods[[grp]]$S)
  }

  save(output.modtest, file = 'diffmods.test.robustness.output.d211015.rdata')
}

sapply(output.modtest, function(x) sapply(x, dim))
sort(sapply(output.modtest, length), decreasing = T)
sum(sort(sapply(output.modtest, length), decreasing = T))
# [1] 107

{
  par(mfrow = c(1, 1), omi = c(2, .1, 0, 0), mar = c(5, 4, 2.5, 0), mgp = c(2.5, 1, 0), cex = 1.5)
  barplot(sort(sapply(output.modtest, length), decreasing = T), las = 2, 
          ylab = 'Number of significant modules', 
          main = 'WGCNA significant modules', cex.main = 1.8)
}


### Volcano plots ###
{
  all.pairs = NULL
  for ( i in 1:length(output.modtest) ) {
    tmp1 = names(output.modtest)[i]
    tmp2 = output.modtest[[i]]
    if ( length(tmp2) == 0 ) next
    
    for ( j in 1:length(tmp2) ) {
      tmp3 = names(tmp2)[j]
      tmp4 = tmp2[[j]]
      tmp5 = apply(tmp4, 1, function(x) paste0(tmp1, ':', tmp3, ';', x[1], '|', x[2]))
      all.pairs = c(all.pairs, tmp5)
    }
  }
  
  length(all.pairs)
  # [1] 1926
  head(all.pairs)
  
  abs.fc = as.numeric(unlist(lapply(output.modtest, function(x) unlist(lapply(x, function(y) y[, 'AbsFC'])))))
  p.wc = as.numeric(unlist(lapply(output.modtest, function(x) unlist(lapply(x, function(y) y[, 'PvalWC'])))))
  p.ks = as.numeric(unlist(lapply(output.modtest, function(x) unlist(lapply(x, function(y) y[, 'PvalKS'])))))
  
  data.volca = cbind(aFC = abs.fc, pWC = p.wc, pKS = p.ks)
  rownames(data.volca) = all.pairs
  dim(data.volca)
  # [1] 1926    3
  head(data.volca)
  
  data.volca[order(data.volca[, 'aFC'], decreasing = T), ][1:10, ]
  
  data.volca = data.volca[apply(data.volca, 1, function(x) all(!is.na(x))), ]
  dim(data.volca)
  # [1] 1277    3
  
  apply(data.volca, 2, range)
  #           aFC       pWC       pKS
  # [1,] 1.000001 0.0000000 0.0000000
  # [2,] 4.887491 0.9993719 0.9970968
  
  apply(data.volca, 2, function(x) range(x[x > 0]))
  #           aFC           pWC          pKS
  # [1,] 1.000001 4.871760e-303 1.110223e-16
  # [2,] 4.887491  9.993719e-01 9.970968e-01
  
  apply(data.volca, 2, function(x) sum(x == 0))
  # aFC pWC pKS 
  #   0   8 154

  min(apply(data.volca, 2, function(x) range(x[x > 0])))
  # [1] 4.87176e-303
  
  data.volca[, 'pWC'][data.volca[, 'pWC'] == 0] = 1e-304
  data.volca[, 'pKS'][data.volca[, 'pKS'] == 0] = 1e-304
  
  range(-log10(data.volca[, 'pWC']))
  # [1] 2.728647e-04 3.040000e+02
  range(-log10(data.volca[, 'pKS']))
  # [1] 1.262671e-03 3.040000e+02
  
  par(mfrow = c(3, 1))
  apply(data.volca, 2, hist)
  
  {
    par(mfrow = c(2, 1), omi = c(.2, .1, 0, 0), mar = c(4, 4, 2, 1), cex = 1.2)
    hist(data.volca[, 'aFC'], br = 20, xlab = 'FC', 
         main = 'Fold changes for all condition pairs', cex.main = 1.5)
    plot(sort(data.volca[, 'aFC'], decreasing = T), log = 'xy', 
         xlab = 'Ordered index of condition pairs (log scale)', ylab = 'FC (log scale)')
    abline(h = c(1.3, 1.5, 2), col = 'red', lty = 2)
  }
  
  {
    par(mfrow = c(2, 2), omi = c(.2, .1, .5, 0), mar = c(4, 4, 1.5, 1), cex = 1)
    plot(data.volca[, 'pWC'], data.volca[, 'pKS'], xlab = 'p-W', ylab = 'p-KS')
    plot(data.volca[, 'pWC'], data.volca[, 'pKS'], xlab = 'p-W', ylab = 'p-KS', 
         xlim = c(0, 0.1), ylim = c(0, 0.1))
    plot(data.volca[, 'pWC'], data.volca[, 'pKS'], log = 'xy', 
         xlab = 'p-W', ylab = 'p-KS')
    plot(data.volca[, 'pWC'] + 1e-10, data.volca[, 'pKS'] + 1e-10, log = 'xy', 
         xlab = 'p-W + 1e-10', ylab = 'p-KS + 1e-10')
    title(main = 'P-values from all Wilcoxon and K-S tests', outer = T, cex.main = 1.8)
  }
  
  {
    par(mfrow = c(2, 2), omi = c(.2, .1, 0, 0), mar = c(4, 4, 2, 1), cex = 1)
    plot(-log10(data.volca[, 'pWC']), -log10(data.volca[, 'pKS']), 
         xlab = 'p-W (-log10)', ylab = 'p-KS (-log10)')
    plot(-log10(data.volca[, 'pWC'] + 1e-3), -log10(data.volca[, 'pKS'] + 1e-3), 
         xlab = 'p-W (-log10)', ylab = 'p-KS (-log10)')
    plot(-log10(data.volca[, 'pWC'] + 1e-6), -log10(data.volca[, 'pKS'] + 1e-6), 
         xlab = 'p-W (-log10)', ylab = 'p-KS (-log10)')
    plot(-log10(data.volca[, 'pWC'] + 1e-20), -log10(data.volca[, 'pKS'] + 1e-20), 
         xlab = 'p-W (-log10)', ylab = 'p-KS (-log10)')
  }
  
  {
    par(mfrow = c(2, 2), omi = c(.2, .1, 0, 0), mar = c(4, 4, 2, 1), cex = 1)
    plot(data.volca[, 'aFC'], -log10(data.volca[, 'pWC']), log = 'xy')
    plot(data.volca[, 'aFC'], -log10(data.volca[, 'pKS']), log = 'xy')
    plot(data.volca[, 'aFC'], -log10((data.volca[, 'pWC'] + data.volca[, 'pKS']) / 2), log = 'xy')
    abline(v = 1.3, col = 'green', lty = 2)
    abline(h = c(1, 3), col = 'red', lty = 2)
    plot(data.volca[, 'aFC'], -log10((data.volca[, 'pWC'] + data.volca[, 'pKS']) / 2), log = 'y')
    abline(v = 1.3, col = 'green', lty = 2)
    abline(h = c(1, 3), col = 'red', lty = 2)
  }
  
  {
    par(mfrow = c(2, 2), omi = c(.2, .1, 0, 0), mar = c(4, 4, 2, 1), cex = 1)
    plot(data.volca[, 'aFC'], -log10(data.volca[, 'pWC'] + 1e-10), log = 'xy')
    plot(data.volca[, 'aFC'], -log10(data.volca[, 'pKS'] + 1e-10), log = 'xy')
    plot(-log10(data.volca[, 'pWC'] + 1e-10), -log10(data.volca[, 'pKS'] + 1e-10))
    plot(data.volca[, 'aFC'], -log10((data.volca[, 'pWC'] + data.volca[, 'pKS']) / 2 + 1e-10), log = 'x')
    abline(v = 1.3, col = 'green', lty = 2)
    abline(h = c(1, 3), col = 'red', lty = 2)
  }
  
  {
    par(mfrow = c(1, 1), omi = c(.2, .1, 0, 0), mar = c(4, 4, 2.5, 1), cex = 1.5)
    plot(data.volca[, 'aFC'], -log10((data.volca[, 'pWC'] + data.volca[, 'pKS']) / 2 + 1e-10), 
         log = 'x', xlab = 'FC (log scale)', ylab = '-log10(mean(p-W, p-KS) + 1e-10)', 
         main = 'Fold change vs. p-value', cex.main = 1.5)
    abline(v = 1.3, col = 'green', lty = 2)
    abline(h = c(1, 3), col = 'red', lty = 2)
  }
}

{ ### Significant condition pairs ###
  sig.pairs = rownames(data.volca)[apply(data.volca, 1, function(x) 
                                   x['aFC'] > 1.3 & (mean(x[c('pWC', 'pKS')]) < .05))]
  length(sig.pairs)
  # [1] 251
  
  ### for each cell type
  sort(table(sapply(strsplit(sig.pairs, '[:]'), function(x) x[1])), decreasing = T)
  
  par(mfrow = c(1, 1), omi = c(1, 0, 0, 0))
  barplot(sort(table(sapply(strsplit(sig.pairs, '[:]'), function(x) x[1])), decreasing = T), las = 2)
  
  ### for each time point
  sig.pairs.bytime = sapply(c('T1', 'T2', 'T3'), function(x) 
                            grep(paste0('[|]', x, '_'), 
                                 grep(paste0('[;]', x, '_'), sig.pairs, value = T), value = T))
  
  sapply(sig.pairs.bytime, length)
  # T1 T2 T3 
  # 34 20 27
  
  par(mfrow = c(1, 1), omi = c(0, 0, 0, 0))
  barplot(sapply(sig.pairs.bytime, length))
}

{ ### FDR ###
  data.volca = cbind(aFC = abs.fc, pWC = p.wc, pKS = p.ks)
  rownames(data.volca) = all.pairs
  dim(data.volca)
  # [1] 1926    3
  
  head(data.volca)
  
  data.volca = data.volca[apply(data.volca, 1, function(x) all(!is.na(x))), ]
  dim(data.volca)
  # [1] 1277    3
  
  p.wc.fdr = p.adjust(data.volca[, 'pWC'], method = 'fdr')
  p.ks.fdr = p.adjust(data.volca[, 'pKS'], method = 'fdr')
  
  plot(data.volca[, 'pWC'], p.wc.fdr)
  plot(data.volca[, 'pKS'], p.ks.fdr)
  
  cor(data.volca[, 'pWC'], p.wc.fdr, method = 'spearman')
  # [1] 0.9999979
  cor(data.volca[, 'pKS'], p.ks.fdr, method = 'spearman')
  # [1] 0.9999975
  
  range(p.wc.fdr)
  # [1] 0.0000000 0.9993719
  range(p.ks.fdr)
  # [1] 0.0000000 0.9970968
  
  range(p.wc.fdr[p.wc.fdr > 0])
  # [1] 6.912486e-301  9.993719e-01
  range(p.ks.fdr[p.ks.fdr > 0])
  # [1] 9.088172e-16 9.970968e-01
  
  sum(p.wc.fdr == 0)
  # [1] 8
  sum(p.ks.fdr == 0)
  # [1] 154
}

{
  data.volca[, 'pWC'] = p.wc.fdr
  data.volca[, 'pKS'] = p.ks.fdr
}


### Number of significant modules with different p-value thresholds and the minimum FC > 1.3
sigmods.thres = list()
for ( THRES in c(seq(0.001, 0.009, 0.001), seq(0.01, 0.1, 0.01)) ) {
  tmp.sigmods = list()
  for ( g in names(output.modtest) ) {
    tmp.sigmods[[g]] = lapply(output.modtest[[g]], function(x) {
                              z = t(apply(x, 1, function(y) 
                                          c(State = paste0(y['Group1'], '|', y['Group2']), 
                                            #Cond = (min(as.numeric(y[c('PvalWC', 'PvalKS')])) < THRES & 
                                            Cond = (mean(as.numeric(y[c('PvalWC', 'PvalKS')])) < THRES & 
                                                    y['AbsFC'] > 1.3))))
                              z = z[apply(z, 1, function(j) j[2] == 'TRUE'), 'State']
                              z[!is.na(z)]
                             })
  }
  
  tmp.sigmods = lapply(tmp.sigmods, function(x) x[sapply(x, length) > 0])
  sigmods.thres[[as.character(THRES)]] = tmp.sigmods
}

if ( !file.exists('wgcna.sigmods.by.thres.d211015.rdata') ) {
  save(sigmods.thres, file = 'wgcna.sigmods.by.thres.d211015.rdata')
}

# Number of differential modules
sapply(sigmods.thres, function(x) sapply(x, length))

# Differential condition pairs
lapply(sigmods.thres, function(x) sapply(x, function(y) unlist(y)))
sapply(sigmods.thres, function(x) sapply(x, function(y) length(unlist(y))))

# Differential modules for each time point
{
  sapply(sigmods.thres, function(x) sapply(x, function(y) 
    sum(sapply(y, function(z) length(grep('T1_', z)) > 0))))
  sapply(sigmods.thres, function(x) sapply(x, function(y) 
    sum(sapply(y, function(z) length(grep('T2_', z)) > 0))))
  sapply(sigmods.thres, function(x) sapply(x, function(y) 
    sum(sapply(y, function(z) length(grep('T3_', z)) > 0))))
}

{
  (diffmods.thres.t1 = sapply(sigmods.thres, function(x) sapply(x, function(y) 
    sum(sapply(y, function(z) 
      length(grep('[|]T1_', grep('^T1_', z, value = T))) > 0 )))))
  (diffmods.thres.t2 = sapply(sigmods.thres, function(x) sapply(x, function(y) 
    sum(sapply(y, function(z) 
      length(grep('[|]T2_', grep('^T2_', z, value = T))) > 0 )))))
  (diffmods.thres.t3 = sapply(sigmods.thres, function(x) sapply(x, function(y) 
    sum(sapply(y, function(z) 
      length(grep('[|]T3_', grep('^T3_', z, value = T))) > 0 )))))
  
  (sigmods.by.time = rbind(T1 = colSums(diffmods.thres.t1), 
                           T2 = colSums(diffmods.thres.t2), 
                           T3 = colSums(diffmods.thres.t3)))
  
  {
    par(mfrow = c(1, 1), omi = c(.1, .1, 0, 0), cex = 1.5)
    barplot(sigmods.by.time, beside = T, las = 2, col = c('red', 'blue', 'green'), 
            ylab = 'Total number of differential modules', 
            main = 'Differential modules for each time point')
    legend('topleft', bty = 'n', fill = c('red', 'blue', 'green'), ncol = 3, cex = 1.5, 
           legend = c('T1', 'T2', 'T3'))
  }
}

{
  (diffmods.thres.d1 = sapply(sigmods.thres, function(x) sapply(x, function(y) 
    sum(sapply(y, function(z) 
      length(grep('_WNA$', grep('_WNA[|]', z, value = T))) > 0 )))))
  (diffmods.thres.d2 = sapply(sigmods.thres, function(x) sapply(x, function(y) 
    sum(sapply(y, function(z) 
      length(grep('_WNE$', grep('_WNE[|]', z, value = T))) > 0 )))))
  (diffmods.thres.d3 = sapply(sigmods.thres, function(x) sapply(x, function(y) 
    sum(sapply(y, function(z) 
      length(grep('_WNF$', grep('_WNF[|]', z, value = T))) > 0 )))))
  
  (sigmods.by.disease = rbind(WNA = colSums(diffmods.thres.d1), 
                              WNE = colSums(diffmods.thres.d2), 
                              WNF = colSums(diffmods.thres.d3)))
  
  {
    par(mfrow = c(1, 1), omi = c(0, 0, 0, 0), mar = c(4, 4, 2, 1), mgp = c(2.5, 1, 0), cex = 1.5)
    barplot(sigmods.by.disease, beside = T, las = 2, col = c('grey', 'orange', 'dark red'), 
            ylab = 'Total number of differential modules', 
            main = 'Differential modules for each disease state')
    legend('topleft', bty = 'n', fill = c('grey', 'orange', 'dark red'), ncol = 3, cex = 1.5, 
           legend = c('WNA', 'WNE', 'WNF'))
  }
}


##### Differential modules at p = 0.01 #####

diffmods = sigmods.thres$`0.01`

(cond.pairs = apply(output.modtest$`CD14+_Monocytes`$MOD1[, c('Group1', 'Group2'), ], 1, 
                    function(x) paste0(x, collapse = '|')))

diffmods.list = lapply(1:length(diffmods), function(i) {
  tmp.dm = diffmods[[i]]
  
  tmp.stats = output.modtest[[names(diffmods)[i]]][names(tmp.dm)]
  tmp.stats = lapply(1:length(tmp.stats), function(x) {
                      tmp1 = data.frame(tmp.stats[[x]][match(tmp.dm[[x]], 
                                            apply(tmp.stats[[x]][, 1:2], 1, 
                                                  function(y) paste(y, collapse = '|'))), 
                                            c('Med1', 'Med2', 'AbsFC')])
                      tmp2 = apply(tmp1[, c('Med1', 'Med2')], 1, 
                                   function(z) if ( z[1] > z[2] ) 1 else -1)
                      tmp3 = tmp2 * round(as.numeric(tmp1[, 'AbsFC']), 2)
                      names(tmp3) = tmp.dm[[x]]
                      tmp3
                    })
  names(tmp.stats) = names(tmp.dm)
  
  tmp.out = sapply(tmp.stats, function(y) y[match(cond.pairs, names(y))])
  rownames(tmp.out) = cond.pairs
  colnames(tmp.out) = paste0('MOD', 1:ncol(tmp.out))
  t(tmp.out)
})

names(diffmods.list) = names(diffmods)

{
  diffmods.tab = NULL
  for ( i in 1:length(diffmods.list) ) {
    tmp1 = names(diffmods.list)[i]
    tmp2 = diffmods.list[[i]]
    rownames(tmp2) = paste0(tmp1, '|', rownames(tmp2))
    diffmods.tab = rbind(diffmods.tab, tmp2)
  }
  
  dim(diffmods.tab)
  # [1] 47 18
}

if ( !file.exists('diffmods.p0.01.d211015.rdata') ) {
  save(diffmods, diffmods.list, diffmods.tab, file = 'diffmods.p0.01.d211015.rdata')
}

{
  par(omi = c(1.1, 0, .1, 1.5), cex.main = 1.8)
  heatmap.2(diffmods.tab, trace = 'n', 
            keysize = .5, key.par = list(cex.main = 1), key.xlab = 'Score', 
            col = colorpanel(10, 'blue', 'red'), na.color = 'lightgrey', 
            Colv = F, Rowv = F, dendrogram = 'none', 
            cexRow = 1.2, cexCol = 1.5, 
            adjRow = c(.01, .5), adjCol = c(.95, .5), 
            lhei = c(1, 6), lwid = c(1, 5), 
            sepcolor = 'white', sepwidth = c(.001, .001), 
            colsep = 0:ncol(diffmods.tab), rowsep = 0:nrow(diffmods.tab), 
            main = 'Differential modules\n(p < 0.01; FC > 1.3)')
  
  par(omi = c(1.1, 0, 0, 1.5), cex.main = 1.6)
  heatmap.2(diffmods.tab, trace = 'n', key = F, 
            col = c('blue', 'red'), na.color = 'lightgrey', 
            Colv = F, Rowv = F, dendrogram = 'none', 
            cexRow = 1.2, cexCol = 1.5, 
            adjRow = c(.01, .5), adjCol = c(.95, .5), 
            lhei = c(1, 9), lwid = c(1, 15), 
            sepcolor = 'white', sepwidth = c(.001, .001), 
            colsep = 0:ncol(diffmods.tab), rowsep = 0:nrow(diffmods.tab), 
            main = 'Differential modules\n(p < 0.01; FC > 1.3)')
  
  par(omi = c(1.1, 0, 0, 1.5), cex.main = 1.6)
  heatmap.2(diffmods.tab, trace = 'n', key = F, 
            col = c('blue', 'red'), na.color = 'black', 
            Colv = F, Rowv = F, dendrogram = 'none', 
            cexRow = 1.2, cexCol = 1.5, 
            adjRow = c(.01, .5), adjCol = c(.95, .5), 
            lhei = c(1, 9), lwid = c(1, 15), 
            sepcolor = 'grey', sepwidth = c(.001, .001), 
            colsep = 0:ncol(diffmods.tab), rowsep = 0:nrow(diffmods.tab), 
            main = 'Differential modules\n(p < 0.01; FC > 1.3)')
  
  par(omi = c(1.1, 0, .1, 1.5), cex.main = 1.8)
  heatmap.2(diffmods.tab, trace = 'n', 
            keysize = .5, key.par = list(cex.main = 1), key.xlab = 'Score', 
            col = colorpanel(20, 'blue', 'white', 'red'), na.color = 'white', 
            Colv = F, Rowv = F, dendrogram = 'none', 
            cexRow = 1.2, cexCol = 1.5, 
            adjRow = c(.01, .5), adjCol = c(.95, .5), 
            lhei = c(1, 6), lwid = c(1, 5), 
            sepcolor = 'black', sepwidth = c(.001, .001), 
            colsep = 0:ncol(diffmods.tab), rowsep = 0:nrow(diffmods.tab), 
            main = 'Differential modules\n(p < 0.01; FC > 1.3)')
  
  diffmods.tab2 = diffmods.tab
  diffmods.tab2[is.na(diffmods.tab2)] = 0
  par(omi = c(1.1, 0, .1, 1.5), cex.main = 1.8)
  heatmap.2(diffmods.tab2, trace = 'n', 
            keysize = .5, key.par = list(cex.main = 1), key.xlab = 'Score', 
            col = colorpanel(20, 'blue', 'white', 'red'), 
            Colv = F, Rowv = T, dendrogram = 'row', 
            cexRow = 1.2, cexCol = 1.5, 
            adjRow = c(.01, .5), adjCol = c(.95, .5), 
            lhei = c(1, 6), lwid = c(1, 5), 
            sepcolor = 'black', sepwidth = c(.001, .001), 
            colsep = 0:ncol(diffmods.tab2), rowsep = 0:nrow(diffmods.tab2), 
            main = 'Differential modules\n(p < 0.01; FC > 1.3)')
  
  diffmods.tab3 = diffmods.tab
  diffmods.tab3[is.na(diffmods.tab3)] = 0
  diffmods.tab3[diffmods.tab3 > 0] = 1
  diffmods.tab3[diffmods.tab3 < 0] = -1
  heatmap.2(diffmods.tab3, trace = 'n', key = F, 
            col = c('blue', 'black', 'red'), 
            Colv = F, Rowv = T, dendrogram = 'row', 
            cexRow = 1.2, cexCol = 1.5, 
            adjRow = c(.01, .5), adjCol = c(.95, .5), 
            lhei = c(1, 6), lwid = c(1, 5), 
            sepcolor = 'grey', sepwidth = c(.001, .001), 
            colsep = 0:ncol(diffmods.tab3), rowsep = 0:nrow(diffmods.tab3), 
            main = 'Differential modules\n(p < 0.01; FC > 1.3)')
}

{
  pdf(file = 'diffmods.p0.01.heatmap.v1.d211015.pdf', width = 10, height = 12)
  
  par(omi = c(1.1, 0, 0, 1.5), cex.main = 1.6)
  heatmap.2(diffmods.tab, trace = 'n', key = F, 
            col = c('blue', 'red'), na.color = 'black', 
            Colv = F, Rowv = F, dendrogram = 'none', 
            cexRow = 1.2, cexCol = 1.5, 
            adjRow = c(.01, .5), adjCol = c(.95, .5), 
            lhei = c(1, 9), lwid = c(1, 15), 
            sepcolor = 'grey', sepwidth = c(.001, .001), 
            colsep = 0:ncol(diffmods.tab), rowsep = 0:nrow(diffmods.tab), 
            main = 'Differential modules\n(p < 0.01; FC > 1.3)')
  
  par(omi = c(1.1, 0, .1, 1.5), cex.main = 1.8)
  heatmap.2(diffmods.tab2, trace = 'n', 
            keysize = .5, key.par = list(cex.main = 1), key.xlab = 'Score', 
            col = colorpanel(20, 'blue', 'white', 'red'), 
            Colv = F, Rowv = T, dendrogram = 'row', 
            cexRow = 1.2, cexCol = 1.5, 
            adjRow = c(.01, .5), adjCol = c(.95, .5), 
            lhei = c(1, 6), lwid = c(1, 5), 
            sepcolor = 'black', sepwidth = c(.001, .001), 
            colsep = 0:ncol(diffmods.tab2), rowsep = 0:nrow(diffmods.tab2), 
            main = 'Differential modules\n(p < 0.01; FC > 1.3)')
  
  dev.off()
}

{ ### Figure 3 for the manuscript ###
  png(file = 'fig3b.diffmods.heatmap.v1.d211015.png', width = 1600, height = 1800, res = 150)
  
  diffmods.tab4 = diffmods.tab2
  colnames(diffmods.tab4) = sub('[|]', ' | ', gsub('WN', '', colnames(diffmods.tab4)))
  rownames(diffmods.tab4) = sapply(gsub('MOD', '', rownames(diffmods.tab4)), 
                                   function(x) paste(strsplit(x, '[|]')[[1]][2:1], collapse = ' | '))
  
  par(omi = c(.5, 0, .1, 1.8), cex.main = 1.8)
  heatmap.2(diffmods.tab4, trace = 'n', 
            keysize = .5, key.par = list(cex.main = 1), key.xlab = 'Score', 
            col = colorpanel(20, 'blue', 'white', 'red'), 
            Colv = F, Rowv = T, dendrogram = 'row', 
            cexRow = 1.2, cexCol = 1.5, 
            adjRow = c(.01, .5), adjCol = c(.95, .5), 
            lhei = c(1, 6), lwid = c(1, 5), 
            sepcolor = 'black', sepwidth = c(.001, .001), 
            colsep = 0:ncol(diffmods.tab4), rowsep = 0:nrow(diffmods.tab4), 
            main = 'Differential modules')
  
  dev.off()
}

{ ### Binary heatmap ###
  diffmods.tab.bin = diffmods.tab
  diffmods.tab.bin[!is.na(diffmods.tab.bin)] = 1
  diffmods.tab.bin[is.na(diffmods.tab.bin)] = 0
  
  par(omi = c(1.1, 0, 0, 1.5), cex.main = 1.6)
  heatmap.2(diffmods.tab.bin, trace = 'n', key = F, col = c('black', 'red'), 
            Colv = F, Rowv = F, dendrogram = 'none', 
            cexRow = 1.2, cexCol = 1.5, 
            adjRow = c(.01, .5), adjCol = c(.95, .5), 
            lhei = c(1, 9), lwid = c(1, 15), 
            sepcolor = 'white', sepwidth = c(.001, .001), 
            colsep = 1:ncol(diffmods.tab.bin), rowsep = 1:nrow(diffmods.tab.bin), 
            main = 'Differential modules\n(p < 0.01; FC > 1.3)')
  
  par(omi = c(1.1, 0, 0, 1.5), cex.main = 1.6)
  heatmap.2(diffmods.tab.bin, trace = 'n', key = F, col = c('black', 'red'), 
            Colv = F, Rowv = T, dendrogram = 'row', 
            cexRow = 1.2, cexCol = 1.5, 
            adjRow = c(.01, .5), adjCol = c(.95, .5), 
            lhei = c(1, 9), lwid = c(1, 6), 
            sepcolor = 'white', sepwidth = c(.001, .001), 
            colsep = 1:ncol(diffmods.tab.bin), rowsep = 1:nrow(diffmods.tab.bin), 
            main = 'Differential modules\n(p < 0.01; FC > 1.3)')
}


for ( grp in names(diffmods) ) {
  message(grp)
  
  tmp.diffmods = wgcna.signi.mods[[grp]]$M[names(diffmods[[grp]])]
  tmp.metadt = wgcna.signi.mods[[grp]]$S[, c('state_timepoint', names(tmp.diffmods))]
  
  names(tmp.diffmods) = paste0('MOD', 1:length(tmp.diffmods))
  colnames(tmp.metadt)[-1] = names(tmp.diffmods)

  ###########################################################
  # Table of genes for each module
  if ( length(tmp.diffmods) > 1 ) {
    easy_mod = t(plyr::ldply(tmp.diffmods, rbind))[-1, ]
  } else {
    easy_mod = data.matrix(tmp.diffmods[[1]])
  }
  colnames(easy_mod) = names(tmp.diffmods)
  
  write.csv(easy_mod, file = paste0(grp, '.final.p0.01.diffmods.d211015.csv'), row.names = F)
  
  #############################################################
  # Box-whisker plots of the final modules
  tmp1 = table(tmp.metadt$state_timepoint)
  tmp2 = names(which(tmp1 >= 10))
  tmp.data = tmp.metadt[tmp.metadt$state_timepoint %in% tmp2, ]
  
  tmp.colors = rainbow(9)[levels(wnv@meta.data$state_timepoint) %in% tmp.data$state_timepoint]
  cell_object.geneclusts.time.final = lapply(names(tmp.diffmods), function(c_name) {
    p = ggplot(tmp.data, aes_string(x = 'state_timepoint', y = c_name, fill = 'state_timepoint')) +
      geom_boxplot() + scale_fill_manual(values = tmp.colors) + xlab('') + 
      theme(legend.position = 'none', axis.text = element_text(size = 16), axis.text.x = element_text(angle = 90), 
            axis.title = element_text(size = 16))
    return(p)
  })
  
  options(repr.plot.width = 6 * length(cell_object.geneclusts.time.final), repr.plot.height = 6)
  print(plot_grid(plotlist = cell_object.geneclusts.time.final, nrow = 1))
  
  ggsave(paste0(grp, '.final.p0.01.diffmods.d211015.png'), limitsize = F, 
         width = 6 * length(cell_object.geneclusts.time.final), height = 6)
}


for ( grp in names(diffmods) ) {
  message(grp)
  
  tmp.diffmods = wgcna.signi.mods[[grp]]$M[names(diffmods[[grp]])]
  tmp.metadt = wgcna.signi.mods[[grp]]$S[, c('state_timepoint', names(tmp.diffmods))]
  tmp.metadt = cbind(patient = as.character(group.list[[grp]]@meta.data[, 'U19.ID']), tmp.metadt)
  
  names(tmp.diffmods) = paste0('MOD', 1:length(tmp.diffmods))
  colnames(tmp.metadt)[grep('MOD', colnames(tmp.metadt))] = names(tmp.diffmods)
  
  #############################################################
  # Box-whisker plots of the final modules at the patient level
  tmp1 = table(tmp.metadt$state_timepoint)
  tmp2 = names(which(tmp1 >= 10))
  tmp.data = tmp.metadt[tmp.metadt$state_timepoint %in% tmp2, ]
  
  tmp.colors = rainbow(11)[unique(wnv@meta.data$U19.ID) %in% tmp.data$patient]
  cell_object.geneclusts.time.final = lapply(names(tmp.diffmods), function(c_name) {
    p = ggplot(tmp.data, aes_string(x = 'state_timepoint', y = c_name, fill = 'patient')) +
      geom_boxplot() + xlab('') + 
      scale_fill_manual(values = tmp.colors) + 
      theme(legend.position = 'none', axis.text = element_text(size = 16), axis.text.x = element_text(angle = 90), 
            axis.title = element_text(size = 16))
    return(p)
  })
  
  options(repr.plot.width = 6 * length(cell_object.geneclusts.time.final), repr.plot.height = 6)
  print(plot_grid(plotlist = cell_object.geneclusts.time.final, nrow = 1))
  
  ggsave(paste0(grp, '.final.p0.01.diffmods.bysamples.png'), limitsize = F, 
         width = 6 * length(cell_object.geneclusts.time.final), height = 6)
}

for ( grp in names(diffmods) ) {
  message(grp)
  
  tmp.diffmods = wgcna.signi.mods[[grp]]$M[names(diffmods[[grp]])]
  tmp.metadt = wgcna.signi.mods[[grp]]$S[, c('state_timepoint', names(tmp.diffmods))]
  tmp.metadt = cbind(patient = as.character(group.list[[grp]]@meta.data[, 'U19.ID']), tmp.metadt)
  
  names(tmp.diffmods) = paste0('MOD', 1:length(tmp.diffmods))
  colnames(tmp.metadt)[grep('MOD', colnames(tmp.metadt))] = names(tmp.diffmods)
  
  #############################################################
  # Box-whisker plots of the final modules for patient-level median expression
  tmp1 = table(tmp.metadt$state_timepoint)
  tmp2 = names(which(tmp1 >= 10))
  tmp.data = tmp.metadt[tmp.metadt$state_timepoint %in% tmp2, ]
  
  tmp.medi = split(tmp.data, as.character(tmp.data$state_timepoint))
  
  tmp.median.cond = t(sapply(tmp.medi, function(x) apply(x[, -c(1, 2)], 2, median)))
  tmp.median.cond = data.frame(state_timepoint = rownames(tmp.median.cond), tmp.median.cond)
  
  tmp.medi = lapply(tmp.medi, function(x) split(x, as.character(x$patient)))
  tmp.medi = lapply(tmp.medi, function(x) t(sapply(x, 
                              function(y) c(patient = y[1, 1], 
                                           state_timepoint = as.character(y[1, 2]), 
                                           apply(data.frame(y[, -c(1,2)]), 2, median)))))
  
  tmp.median = NULL
  for ( i in 1:length(tmp.medi) ) {
    tmp.median = rbind(tmp.median, tmp.medi[[i]])
  }
  
  tmp.median = data.frame(tmp.median)
  if ( length(tmp.diffmods) == 1 ) colnames(tmp.median)[3] = 'MOD1'
  
  cell_object.geneclusts.time.final = lapply(names(tmp.diffmods), function(c_name) {
    tmp.median[[c_name]] = as.numeric(tmp.median[[c_name]])
    
    tmp.colors = rainbow(9)[levels(wnv@meta.data$state_timepoint) %in% tmp.data$state_timepoint]
    p = ggplot(tmp.data, aes_string(x = 'state_timepoint', y = c_name, fill = 'state_timepoint')) +
      geom_boxplot() + xlab('') + 
      scale_fill_manual(values = tmp.colors) + 
      theme(legend.position = 'none', axis.text = element_text(size = 16), axis.text.x = element_text(angle = 90), 
            axis.title = element_text(size = 16)) + 
      geom_point(tmp.median.cond, mapping = aes_string(x = 'state_timepoint', y = c_name), 
                 shape = 17, fill = 'black', color = 'black', size = 6) + 
      geom_point(tmp.median, mapping = aes_string(x = 'state_timepoint', y = c_name), 
                 shape = 1, fill = 'black', color = 'darkred', size = 6)
    
    return(p)
  })
  
  options(repr.plot.width = 6 * length(cell_object.geneclusts.time.final), repr.plot.height = 6)
  print(plot_grid(plotlist = cell_object.geneclusts.time.final, nrow = 1))
  
  ggsave(paste0(grp, '.final.p0.01.diffmods.sample.medians.d211015.png'), limitsize = F, 
         width = 6 * length(cell_object.geneclusts.time.final), height = 6)
}

system.time({
  plots.list = list()
  for ( grp in names(diffmods) ) {
    message(grp)
    
    tmp.diffmods = wgcna.signi.mods[[grp]]$M[names(diffmods[[grp]])]
    tmp.metadt = wgcna.signi.mods[[grp]]$S[, c('state_timepoint', names(tmp.diffmods))]
    tmp.metadt = cbind(patient = as.character(group.list[[grp]]@meta.data[, 'U19.ID']), tmp.metadt)
    
    names(tmp.diffmods) = paste0('MOD', 1:length(tmp.diffmods))
    colnames(tmp.metadt)[grep('MOD', colnames(tmp.metadt))] = names(tmp.diffmods)
    
    #############################################################
    # Box-whisker plots of the final modules for patient-level median expression
    tmp1 = table(tmp.metadt$state_timepoint)
    tmp2 = names(which(tmp1 >= 10))
    tmp.data = tmp.metadt[tmp.metadt$state_timepoint %in% tmp2, ]
    
    tmp.medi = split(tmp.data, as.character(tmp.data$state_timepoint))
    
    if ( length(tmp.diffmods) == 1 ) {
      tmp.median.cond = data.frame(state_timepoint = names(tmp.medi), 
                                   MOD1 = sapply(tmp.medi, function(x) apply(data.frame(x[, -c(1, 2)]), 2, median)))
    } else {
      tmp.median.cond = t(sapply(tmp.medi, function(x) apply(data.frame(x[, -c(1, 2)]), 2, median)))
      tmp.median.cond = data.frame(state_timepoint = rownames(tmp.median.cond), tmp.median.cond)
    }
    
    tmp.medi = lapply(tmp.medi, function(x) split(x, as.character(x$patient)))
    tmp.medi = lapply(tmp.medi, function(x) t(sapply(x, 
                                                     function(y) c(patient = y[1, 1], 
                                                                   state_timepoint = as.character(y[1, 2]), 
                                                                   apply(data.frame(y[, -c(1,2)]), 2, median)))))
    
    tmp.median = NULL
    for ( i in 1:length(tmp.medi) ) {
      tmp.median = rbind(tmp.median, tmp.medi[[i]])
    }
    
    tmp.median = data.frame(tmp.median)
    if ( length(tmp.diffmods) == 1 ) colnames(tmp.median)[3] = 'MOD1'
    
    cell_object.geneclusts.time.final = lapply(names(tmp.diffmods), function(c_name) {
      tmp.median[[c_name]] = as.numeric(tmp.median[[c_name]])
      
      tmp.colors = rainbow(9)[levels(wnv@meta.data$state_timepoint) %in% tmp.median$state_timepoint]
      p = ggplot(tmp.data, aes_string(x = 'state_timepoint', y = c_name, fill = 'state_timepoint')) +
        geom_violin() + xlab('') + ylab('Score') + 
        ggtitle(paste0(grp, '\n', c_name)) + 
        scale_fill_manual(values = tmp.colors) + 
        theme(plot.title = element_text(size = 30, hjust = 0.5), legend.position = 'none', 
              axis.text = element_text(size = 20), axis.text.x = element_text(angle = 90), 
              axis.title = element_text(size = 20)) + 
        geom_point(tmp.median.cond, mapping = aes_string(x = 'state_timepoint', y = c_name), 
                   shape = 17, fill = 'black', color = 'black', size = 6) + 
        geom_point(tmp.median, mapping = aes_string(x = 'state_timepoint', y = c_name), 
                   shape = 1, fill = 'black', color = 'darkred', size = 6)
      
      return(p)
    })
    
    plots.list = c(plots.list, cell_object.geneclusts.time.final)
  }
  
  pdf('diffmods.p0.01.with.sample.medians.violin.d211015.pdf')
  invisible(lapply(plots.list, print))
  dev.off()
})
#   user  system elapsed 
# 24.436   2.575  10.683

{
  library(RColorBrewer)
  par(mar = c(3, 4, 2, 2))
  display.brewer.all()
  
  blues9
  
  my.colors1 = c(brewer.pal(5, 'Blues')[3:1], 
                 brewer.pal(5, 'Reds')[4:2], 
                 brewer.pal(5, 'Oranges')[4:2])
  my.colors1 = my.colors1[c(1, 4, 7, 2, 5, 8, 3, 6, 9)]
  
  my.colors2 = rep(c(brewer.pal(5, 'Blues')[2], 
                     brewer.pal(5, 'Reds')[3], 
                     brewer.pal(5, 'Oranges')[3]), 3)
}

system.time({
  plots.list = list()
  for ( grp in names(diffmods) ) {
    message(grp)
    
    tmp.diffmods = wgcna.signi.mods[[grp]]$M[names(diffmods[[grp]])]
    tmp.metadt = wgcna.signi.mods[[grp]]$S[, c('state_timepoint', names(tmp.diffmods))]
    tmp.metadt = cbind(patient = as.character(group.list[[grp]]@meta.data[, 'U19.ID']), tmp.metadt)
    
    names(tmp.diffmods) = paste0('MOD', 1:length(tmp.diffmods))
    colnames(tmp.metadt)[grep('MOD', colnames(tmp.metadt))] = names(tmp.diffmods)
    
    #############################################################
    # Box-whisker plots of the final modules for patient-level median expression
    tmp1 = table(tmp.metadt$state_timepoint)
    tmp2 = names(which(tmp1 >= 10))
    tmp.data = tmp.metadt[tmp.metadt$state_timepoint %in% tmp2, ]
    
    tmp.medi = split(tmp.data, as.character(tmp.data$state_timepoint))
    
    if ( length(tmp.diffmods) == 1 ) {
      tmp.median.cond = data.frame(state_timepoint = names(tmp.medi), 
                                   MOD1 = sapply(tmp.medi, function(x) apply(data.frame(x[, -c(1, 2)]), 2, median)))
    } else {
      tmp.median.cond = t(sapply(tmp.medi, function(x) apply(data.frame(x[, -c(1, 2)]), 2, median)))
      tmp.median.cond = data.frame(state_timepoint = rownames(tmp.median.cond), tmp.median.cond)
    }
    
    tmp.medi = lapply(tmp.medi, function(x) split(x, as.character(x$patient)))
    tmp.medi = lapply(tmp.medi, function(x) t(sapply(x, 
                                                     function(y) c(patient = y[1, 1], 
                                                                   state_timepoint = as.character(y[1, 2]), 
                                                                   apply(data.frame(y[, -c(1,2)]), 2, median)))))
    
    tmp.median = NULL
    for ( i in 1:length(tmp.medi) ) {
      tmp.median = rbind(tmp.median, tmp.medi[[i]])
    }
    
    tmp.median = data.frame(tmp.median)
    if ( length(tmp.diffmods) == 1 ) colnames(tmp.median)[3] = 'MOD1'
    
    cell_object.geneclusts.time.final = lapply(names(tmp.diffmods), function(c_name) {
      tmp.median[[c_name]] = as.numeric(tmp.median[[c_name]])
      
      tmp.colors = my.colors2[sort(unique(wnv@meta.data$state_timepoint)) %in% tmp.median$state_timepoint]
      #tmp.colors = my.colors1[sort(unique(wnv@meta.data$state_timepoint)) %in% tmp.median$state_timepoint]
      p = ggplot(tmp.data, aes_string(x = 'state_timepoint', y = c_name, fill = 'state_timepoint')) +
        geom_violin() + xlab('') + ylab('Score') + 
        ggtitle(paste0(grp, '\n', c_name)) + 
        scale_fill_manual(values = tmp.colors) + 
        theme(plot.title = element_text(size = 30, hjust = 0.5), legend.position = 'none', 
              axis.text = element_text(size = 20), axis.text.x = element_text(angle = 90), 
              axis.title = element_text(size = 20)) + 
        geom_point(tmp.median.cond, mapping = aes_string(x = 'state_timepoint', y = c_name), 
                   shape = 17, fill = 'black', color = 'black', size = 6) + 
        geom_point(tmp.median, mapping = aes_string(x = 'state_timepoint', y = c_name), 
                   shape = 1, fill = 'black', color = 'darkred', size = 6)
      
      ggsave(filename = paste0('figures/diffmods.p0.01.with.sample.medians.violin.d211214/', 
                               grp, '.', c_name, '.png'), plot = p, width = 7, height = 7)
      return(p)
    })
    
    plots.list = c(plots.list, cell_object.geneclusts.time.final)
  }
  
  pdf('figures/diffmods.p0.01.with.sample.medians.violin.d211214.pdf')
  invisible(lapply(plots.list, print))
  dev.off()
})
#   user  system elapsed 
# 20.833   1.594  23.837


#########################################################################################

##### All module genes #####

#########################################################################################

library(ggplot2)

# (load('wgcna.signi.mods.v1.d211015.rdata'))
# [1] "wgcna.signi.mods"

# (load('diffmods.p0.01.d211015.rdata'))
# [1] "diffmods"      "diffmods.list" "diffmods.tab"

# (load('cellgroups.bycelltype.seuratobj.d211015.rdata'))
# [1] "group.list"

wgcna.signi.mods.genes = lapply(wgcna.signi.mods, function(x) x$M)
sapply(wgcna.signi.mods.genes, length)

system.time({
  plots.list = list()
  for ( grp in names(diffmods) ) {
    message(grp)

    tmp.diffmods = wgcna.signi.mods.genes[[grp]][names(diffmods[[grp]])]
    tmp.diffmods = tmp.diffmods[!sapply(tmp.diffmods, is.null)]
    
    tmp.metadt = wgcna.signi.mods[[grp]]$S[, c('state_timepoint', names(tmp.diffmods))]
    tmp.metadt = cbind(patient = as.character(group.list[[grp]]@meta.data[, 'U19.ID']), tmp.metadt)
    
    names(tmp.diffmods) = paste0('MOD', 1:length(tmp.diffmods))
    colnames(tmp.metadt)[grep('MOD', colnames(tmp.metadt))] = names(tmp.diffmods)
    
    tmp.expr = t(data.frame(group.list[[grp]]@assays$RNA@data)[unlist(tmp.diffmods), rownames(tmp.metadt)])
    #tmp.expr = t(data.frame(group.list[[grp]]@assays$RNA@counts)[unlist(tmp.diffmods), rownames(tmp.metadt)])
    #tmp.expr = t(data.frame(group.list[[grp]]@assays$RNA@scale.data)[unlist(tmp.diffmods), rownames(tmp.metadt)])
    
    tmp.metadt = cbind(tmp.metadt, tmp.expr)
    
    #############################################################
    # Violin plots of the final modules for patient-level median expression
    tmp1 = table(tmp.metadt$state_timepoint)
    tmp2 = names(which(tmp1 >= 10))
    tmp.data = tmp.metadt[tmp.metadt$state_timepoint %in% tmp2, ]
    
    tmp.medi = split(tmp.data, as.character(tmp.data$state_timepoint))
    
    tmp.median.cond = t(sapply(tmp.medi, function(x) apply(x[, -c(1, 2)], 2, median)))
    tmp.median.cond = data.frame(state_timepoint = rownames(tmp.median.cond), tmp.median.cond, 
                                 check.names = F)
    
    tmp.medi = lapply(tmp.medi, function(x) split(x, as.character(x$patient)))
    tmp.medi = lapply(tmp.medi, function(x) t(sapply(x, 
                                                     function(y) c(patient = y[1, 1], 
                                                                   state_timepoint = as.character(y[1, 2]), 
                                                                   apply(data.frame(y[, -c(1,2)]), 2, median)))))
    
    tmp.median = NULL
    for ( i in 1:length(tmp.medi) ) {
      tmp.median = rbind(tmp.median, tmp.medi[[i]])
    }
    
    colnames(tmp.median)[-c(1, 2)] = colnames(tmp.median.cond)[-1]
    
    tmp.median = data.frame(tmp.median, check.names = F)
    if ( length(tmp.diffmods) == 1 ) colnames(tmp.median)[3] = 'MOD1'
    
    tmp.genes = NULL
    for ( i in 1:length(tmp.diffmods) ) {
      tmp.genes = rbind(tmp.genes, cbind(M = names(tmp.diffmods)[i], G = tmp.diffmods[[i]]))
    }
    
    cell_object.geneclusts.time.final = lapply(1:nrow(tmp.genes), function(j) {
      g = as.character(tmp.genes[j, 'G'])
      tmp.median[[g]] = as.numeric(tmp.median[[g]])
      
      tmp.data.g = data.frame(state_timepoint = tmp.data$state_timepoint, gene = tmp.data[[g]])
      tmp.median.cond.g = data.frame(state_timepoint = tmp.median.cond$state_timepoint, gene = tmp.median.cond[[g]])
      tmp.median.g = data.frame(state_timepoint = tmp.median$state_timepoint, gene = tmp.median[[g]])
      
      tmp.colors = rainbow(9)[levels(wnv@meta.data$state_timepoint) %in% tmp.median.g$state_timepoint]
      p = ggplot(tmp.data.g, aes_string(x = 'state_timepoint', y = 'gene', fill = 'state_timepoint')) +
        geom_violin() + xlab('') + ylab('Expression') + 
        ggtitle(paste0(grp, '\n', tmp.genes[j, 'M'], ': ', g)) + 
        scale_fill_manual(values = tmp.colors) + 
        theme(plot.title = element_text(size = 30, hjust = 0.5), legend.position = 'none', 
              axis.text = element_text(size = 20), axis.text.x = element_text(angle = 90), 
              axis.title = element_text(size = 20)) + 
        geom_point(tmp.median.cond.g, mapping = aes_string(x = 'state_timepoint', y = 'gene'), 
                   shape = 17, fill = 'black', color = 'black', size = 6) + 
        geom_point(tmp.median.g, mapping = aes_string(x = 'state_timepoint', y = 'gene'), 
                   shape = 1, fill = 'black', color = 'darkred', size = 6)
      #http://www.sthda.com/english/wiki/ggplot2-point-shapes
      #http://www.sthda.com/english/wiki/ggplot2-box-plot-quick-start-guide-r-software-and-data-visualization
      
      return(p)
    })
    
    plots.list = c(plots.list, cell_object.geneclusts.time.final)
  }
  
  pdf('diffmods.p0.01.all.genes.with.sample.medians.v1.d211015.pdf')
  invisible(lapply(plots.list, print))
  dev.off()
})
#    user  system elapsed
# 932.183 158.932 497.330


system.time({
  expr.mtx = list()
  for ( grp in names(diffmods) ) {
    message(grp)
    
    tmp.diffmods = wgcna.signi.mods.genes[[grp]][names(diffmods[[grp]])]
    tmp.diffmods = tmp.diffmods[!sapply(tmp.diffmods, is.null)]
    
    tmp.metadt = wgcna.signi.mods[[grp]]$S[, c('state_timepoint', names(tmp.diffmods))]
    tmp.metadt = cbind(patient = as.character(group.list[[grp]]@meta.data[, 'U19.ID']), tmp.metadt)
    
    names(tmp.diffmods) = paste0('MOD', 1:length(tmp.diffmods))
    colnames(tmp.metadt)[grep('MOD', colnames(tmp.metadt))] = names(tmp.diffmods)
    
    print(sum(table(unlist(tmp.diffmods)) > 1)) # the number of genes in multiple modules
    
    tmp.expr = t(data.frame(group.list[[grp]]@assays$RNA@data)[unlist(tmp.diffmods), rownames(tmp.metadt)])
    
    tmp.metadt = cbind(tmp.metadt, tmp.expr)
    
    #############################################################
    tmp1 = table(tmp.metadt$state_timepoint)
    tmp2 = names(which(tmp1 >= 10))
    
    expr.mtx[[grp]] = tmp.metadt[tmp.metadt$state_timepoint %in% tmp2, ]
  }
  
  save(expr.mtx, file = 'diffmods.p0.01.all.genes.expr.mtx.d211015.rdata')
})
#   user  system elapsed 
# 61.113  79.796 175.252

sapply(expr.mtx, dim)

genes.all = unlist(sapply(expr.mtx, function(x) colnames(x)[-c(1, 2, grep('^MOD[0-9]{1,2}$', colnames(x)))]))
length(genes.all)
# [1] 1579
length(unique(genes.all))
# [1] 1148

table(table(genes.all))
#   1   2   3   4   5   6   7   8 
# 885 163  63  21   7   5   2   2

sum(table(genes.all) > 1)
# [1] 263

names(which(table(genes.all) == 8))
# [1] "JUN"   "ZFP36"
names(which(table(genes.all) == 7))
# [1] "FOS"  "FOSB"
names(which(table(genes.all) == 6))
# [1] "BHLHE40" "DUSP1"   "GADD45B" "JUNB"    "NR4A2"
names(which(table(genes.all) == 5))
#"DUSP2"     "FTH1"      "GABARAPL1" "IER2"      "KLF4"      "PPP1R15A"  "ZFP36L2"

{
  par(omi = c(0, .2, 0, 0), cex = 1.5)
  barplot(table(table(genes.all)), log = 'y', 
          xlab = 'Frequency in different modules', ylab = 'Number of genes', 
          main = 'Genes in differential modules')
}

{
  genes.unq = names(which(table(genes.all) == 1))
  length(genes.unq)
  # [1] 885
  
  genes.unq.bin = t(sapply(genes.unq, 
                           function(x) sapply(expr.mtx, function(y) as.numeric(x %in% colnames(y)))))
  dim(genes.unq.bin)
  # [1] 885  10
  
  genes.unq.bin = genes.unq.bin[unlist(apply(genes.unq.bin, 2, function(x) which(x == 1))), ]
  
  par(omi = c(2, 0, 0, 0), cex.main = 1.5)
  heatmap.2(genes.unq.bin, trace = 'none', key = F, col = c('black', 'red'), 
            Colv = F, Rowv = F, dendrogram = 'none', 
            labRow = F, cexCol = 1.6, 
            lhei = c(1, 9), lwid = c(1, 9), 
            adjRow = c(.01, .5), adjCol = c(.95, .5), 
            sepcolor = 'white', sepwidth = c(.001, .001), 
            colsep = 0:ncol(genes.unq.bin), #rowsep = 0:nrow(genes.unq.bin), 
            main = 'Unique module genes for each cell type')
}

{
  genes.multi = names(which(table(genes.all) > 1))
  length(genes.multi)
  # [1] 263
  
  genes.multi.bin = t(sapply(genes.multi, 
                             function(x) sapply(expr.mtx, function(y) as.numeric(x %in% colnames(y)))))
  dim(genes.multi.bin)
  # [1] 263  10
  
  par(omi = c(2, 0, 0, 0), cex.main = 1.5)
  heatmap.2(genes.multi.bin, trace = 'none', key = F, col = c('black', 'red'), 
            #Colv = F, Rowv = T, dendrogram = 'row', 
            labRow = F, cexCol = 1.6, 
            lhei = c(1, 6), lwid = c(1, 9), 
            adjRow = c(.01, .5), adjCol = c(.95, .5), 
            sepcolor = 'white', sepwidth = c(.001, .001), 
            colsep = 0:ncol(genes.multi.bin), #rowsep = 0:nrow(genes.multi.bin), 
            main = 'Common module genes in multiple cell types')
}


#########################################################################################

##### Correlation between module scores and gene expression and top 10 module genes #####

#########################################################################################

(load('wgcna.signi.mods.v1.d211015.rdata'))
# [1] "wgcna.signi.mods"

(load('diffmods.p0.01.d211015.rdata'))
# [1] "diffmods"      "diffmods.list" "diffmods.tab"

(load('diffmods.p0.01.all.genes.expr.mtx.d211015.rdata'))
# [1] "expr.mtx"

wgcna.signi.mods.genes = lapply(wgcna.signi.mods, function(x) x$M)

system.time({
  mods.genes.corr = lapply(diffmods, as.null)
  for ( grp in names(diffmods) ) {
    message(grp)
    
    tmp.diffmods = wgcna.signi.mods.genes[[grp]][names(diffmods[[grp]])]
    tmp.diffmods = tmp.diffmods[!sapply(tmp.diffmods, is.null)]
    names(tmp.diffmods) = paste0('MOD', 1:length(tmp.diffmods))
    
    tmp.corr = lapply(tmp.diffmods, as.null)
    for ( i in 1:length(tmp.diffmods) ) {
      tmp.data = expr.mtx[[grp]][, c(names(tmp.diffmods)[i], tmp.diffmods[[i]])]
      tmp.corr[[i]] = sort(apply(tmp.data[, -1], 2, function(x) cor(tmp.data[, 1], x)), decreasing = T)
    }
    
    mods.genes.corr[[grp]] = tmp.corr
  }
})
#  user  system elapsed 
# 0.761   0.242   1.118

{
  sort(sapply(mods.genes.corr, function(x) mean(sapply(x, mean))))
  
  par(omi = c(0, .2, 0, 0), cex = 1.5)
  hist(unlist(sapply(mods.genes.corr, function(x) sapply(x, mean))), 
       xlab = 'Average Pearson correlation', ylab = 'Number of differential modules', 
       main = 'Module scores vs. module gene expression')
  # "diffmods.p0.01.module.scores.vs.gene.expr.d211015.png"
}

{
  lapply(mods.genes.corr, function(x) lapply(x, function(y) y[1:10]))
  
  par(omi = c(0, .2, 0, 0), cex = 1.5)
  plot(1:10, type = 'n', ylim = c(0.3, 1), xaxt = 'n', 
       xlab = 'Ranking', ylab = 'Pearson correlation', 
       main = 'Top 10 signature genes in differential modules')
  axis(1, 1:10)
  lapply(1:length(mods.genes.corr), function(x) 
         lapply(mods.genes.corr[[x]], function(y) 
                points(y[1:10], type = 'b', lwd = 2, col = rainbow(length(mods.genes.corr))[x])))
  legend('bottomleft', bty = 'n', lty = 1, lwd = 2, pch = 21, cex = .8, ncol = 2, 
         col = rainbow(length(mods.genes.corr)), 
         legend = names(mods.genes.corr))
  # "diffmods.p0.01.top10.genes.pcc.d211015.png"
}

{
  genes.top10.pcc = lapply(mods.genes.corr, function(x) lapply(x, function(y) y[1:10]))
  top10.tab = NULL
  for ( i in 1:length(genes.top10.pcc) ) {
    tmp1 = names(genes.top10.pcc)[i]
    tmp2 = t(sapply(genes.top10.pcc[[i]], function(x) c(names(x), x)))
    top10.tab = rbind(top10.tab, cbind(tmp1, rownames(tmp2), tmp2))
  }
  
  dim(top10.tab)
  # [1] 47 22
  
  rownames(top10.tab) = NULL
  colnames(top10.tab) = NULL
  
  write.csv(top10.tab, file = 'diffmods.p0.01.top10.genes.pcc.d211015.csv', quote = F, row.names = F)
}

{
  topgenes = unlist(sapply(genes.top10.pcc, function(x) sapply(x, names)))
  length(topgenes)
  # [1] 470
  length(unique(topgenes))
  # [1] 376
  
  table(table(topgenes))
  #   1   2   3   4   5   6   7 
  # 327  25  14   4   3   1   2
  
  names(which(table(topgenes) == 7))
  # [1] "JUN"   "ZFP36"
  
  par(omi = c(0, .2, 0, 0), cex = 1.5)
  barplot(table(table(topgenes)), log = 'y', 
          xlab = 'Frequency in different modules', ylab = 'Number of modules', 
          main = 'Top 10 genes in differential modules')
  # "diffmods.p0.01.top10.genes.freq.d211015.png"
}

system.time({
  ### Violin plots of the final modules for patient-level median expression
  plots.list = list()
  for ( grp in names(expr.mtx) ) {
    message(grp)
    
    tmp.metadt = expr.mtx[[grp]]
    tmp1 = table(tmp.metadt$state_timepoint)
    tmp2 = names(which(tmp1 >= 10))
    
    tmp.data = tmp.metadt[tmp.metadt$state_timepoint %in% tmp2, ]
    
    tmp.medi = split(tmp.data, as.character(tmp.data$state_timepoint))
    
    tmp.median.cond = t(sapply(tmp.medi, function(x) apply(x[, -c(1, 2)], 2, median)))
    tmp.median.cond = data.frame(state_timepoint = rownames(tmp.median.cond), tmp.median.cond, 
                                 check.names = F)
    
    tmp.medi = lapply(tmp.medi, function(x) split(x, as.character(x$patient)))
    tmp.medi = lapply(tmp.medi, function(x) t(sapply(x, 
                                                     function(y) c(patient = y[1, 1], 
                                                                   state_timepoint = as.character(y[1, 2]), 
                                                                   apply(data.frame(y[, -c(1,2)]), 2, median)))))
    
    tmp.median = NULL
    for ( i in 1:length(tmp.medi) ) {
      tmp.median = rbind(tmp.median, tmp.medi[[i]])
    }
    
    colnames(tmp.median)[-c(1, 2)] = colnames(tmp.median.cond)[-1]
    
    tmp.median = data.frame(tmp.median, check.names = F)
    
    tmp.diffmods = lapply(genes.top10.pcc[[grp]], names)
    tmp.genes = NULL
    for ( i in 1:length(tmp.diffmods) ) {
      tmp.genes = rbind(tmp.genes, cbind(M = names(tmp.diffmods)[i], G = tmp.diffmods[[i]]))
    }
    
    cell_object.geneclusts.time.final = lapply(1:nrow(tmp.genes), function(j) {
      g = as.character(tmp.genes[j, 'G'])
      tmp.median[[g]] = as.numeric(tmp.median[[g]])
      
      tmp.data.g = data.frame(state_timepoint = tmp.data$state_timepoint, gene = tmp.data[[g]])
      tmp.median.cond.g = data.frame(state_timepoint = tmp.median.cond$state_timepoint, gene = tmp.median.cond[[g]])
      tmp.median.g = data.frame(state_timepoint = tmp.median$state_timepoint, gene = tmp.median[[g]])
      
      tmp.colors = rainbow(9)[levels(wnv@meta.data$state_timepoint) %in% tmp.median.g$state_timepoint]
      p = ggplot(tmp.data.g, aes_string(x = 'state_timepoint', y = 'gene', fill = 'state_timepoint')) +
        geom_violin() + xlab('') + ylab('Expression') + 
        ggtitle(paste0(grp, '\n', tmp.genes[j, 'M'], ': ', g)) + 
        scale_fill_manual(values = tmp.colors) + 
        theme(plot.title = element_text(size = 30, hjust = 0.5), legend.position = 'none', 
              axis.text = element_text(size = 20), axis.text.x = element_text(angle = 90), 
              axis.title = element_text(size = 20)) + 
        geom_point(tmp.median.cond.g, mapping = aes_string(x = 'state_timepoint', y = 'gene'), 
                   shape = 17, fill = 'black', color = 'black', size = 6) + 
        geom_point(tmp.median.g, mapping = aes_string(x = 'state_timepoint', y = 'gene'), 
                   shape = 1, fill = 'black', color = 'darkred', size = 6)
      #http://www.sthda.com/english/wiki/ggplot2-point-shapes
      #http://www.sthda.com/english/wiki/ggplot2-box-plot-quick-start-guide-r-software-and-data-visualization
      
      return(p)
    })
    
    plots.list = c(plots.list, cell_object.geneclusts.time.final)
  }
  
  pdf('diffmods.p0.01.top10.genes.with.sample.medians.v1.d211015.pdf')
  invisible(lapply(plots.list, print))
  dev.off()
})
#   user  system elapsed
#241.352  24.157 118.105


#########################################################################################

##### Analysis of the BCR signaling genes from MOFA factor 1 #####

#########################################################################################

bcr.mofa = read.csv('../../WNV/leying/factor1.seqwell.global.mean.bcr.signaling.genes.csv', row.names = 1)
dim(bcr.mofa)
#[1] 22  3
head(bcr.mofa)

(bcr.genes = rownames(bcr.mofa))
# [1] "RELA"    "CAMK2A"  "REL"     "PIP5K1A" "RAPGEF1" "VAV2"    "IRF4"    "LYN"     "NFKBIB"  "BCL10"   "CARD11" 
#[12] "BLNK"    "NFATC2"  "VAV1"    "PPP3CA"  "MAP3K7"  "IKBKG"   "HCLS1"   "PIP5K1C" "ETS1"    "PRKCB"   "GSK3B"

(load('diffmods.p0.01.d211015.rdata'))
#[1] "diffmods"      "diffmods.list" "diffmods.tab"

(load('wgcna.signi.mods.v1.d211015.rdata'))
# [1] "wgcna.signi.mods"

(load('diffmods.p0.01.all.genes.expr.mtx.d211015.rdata'))
# [1] "expr.mtx"

all(names(diffmods) == names(wgcna.signi.mods))
#[1] TRUE

diffmods.genes = lapply(names(diffmods), function(x) {
  tmp = wgcna.signi.mods[[x]]$M[names(diffmods[[x]])]
  names(tmp) = paste0('MOD', 1:length(tmp))
  tmp
  })
names(diffmods.genes) = names(diffmods)

bcr.genes.mod = sapply(diffmods.genes, function(x) {
  tmp = sapply(x, function(y) intersect(y, bcr.genes))
  tmp[sapply(tmp, length) > 0]
  })

bcr.genes.mod = bcr.genes.mod[sapply(bcr.genes.mod, length) > 0]

mods.genes.corr = lapply(bcr.genes.mod, as.null)
for ( grp in names(bcr.genes.mod) ) {
  message(grp)
  
  tmp.diffmods = bcr.genes.mod[[grp]][names(bcr.genes.mod[[grp]])]
  tmp.diffmods = tmp.diffmods[!sapply(tmp.diffmods, is.null)]

  tmp.corr = lapply(tmp.diffmods, as.null)
  for ( i in 1:length(tmp.diffmods) ) {
    #if ( all(!tmp.diffmods[[i]] %in% colnames(expr.mtx[[grp]])) ) next
    tmp.data = expr.mtx[[grp]][, c(names(tmp.diffmods)[i], tmp.diffmods[[i]])]
    tmp.corr[[i]] = sort(apply(data.matrix(tmp.data[, -1]), 2, 
                               function(x) cor(tmp.data[, 1], x)), decreasing = T)
    names(tmp.corr[[i]]) = colnames(tmp.data)[-1]
  }
  
  mods.genes.corr[[grp]] = tmp.corr
}

{
  bcr.genes.mod.tab = NULL
  for ( i in 1:length(bcr.genes.mod) ) {
    tmp1 = names(bcr.genes.mod)[i]
    tmp2 = bcr.genes.mod[[i]]
    
    tmp3 = NULL
    for ( j in 1:length(tmp2) ) {
      tmp3 = rbind(tmp3, cbind(names(tmp2)[j], tmp2[[j]], 
                               mods.genes.corr[[tmp1]][[names(tmp2)[j]]]))
    }
    
    bcr.genes.mod.tab = rbind(bcr.genes.mod.tab, cbind(tmp1, tmp3))
  }
  
  dim(bcr.genes.mod.tab)
  #[1] 6 4
  
  colnames(bcr.genes.mod.tab) = c('CellType', 'Module', 'Gene', 'PCC')
  rownames(bcr.genes.mod.tab) = NULL
  
  write.csv(bcr.genes.mod.tab, quote = F, row.names = F, 
            file = 'diffmods.p0.01.d211015.bcr.signaling.genes.d211221.csv')
}

{
  wnv = get(load('wnv.seqwell.pbmcs.seurat.d211011.Robj'))
  rm(wnv_cleaned)
  #   user  system elapsed 
  #145.050  13.865 164.182
}

{
  library(RColorBrewer)
  my.colors2 = rep(c(brewer.pal(5, 'Blues')[2], 
                     brewer.pal(5, 'Reds')[3], 
                     brewer.pal(5, 'Oranges')[3]), 3)
}

library(ggplot2)

system.time({
  plots.list = list()
  for ( grp in names(bcr.genes.mod) ) {
    message(grp)
    
    tmp.metadt = expr.mtx[[grp]]
    tmp1 = table(tmp.metadt$state_timepoint)
    tmp2 = names(which(tmp1 >= 10))
    
    tmp.data = tmp.metadt[tmp.metadt$state_timepoint %in% tmp2, ]
    
    tmp.medi = split(tmp.data, as.character(tmp.data$state_timepoint))
    
    tmp.median.cond = t(sapply(tmp.medi, function(x) apply(x[, -c(1, 2)], 2, median)))
    tmp.median.cond = data.frame(state_timepoint = rownames(tmp.median.cond), tmp.median.cond, 
                                 check.names = F)
    
    tmp.medi = lapply(tmp.medi, function(x) split(x, as.character(x$patient)))
    tmp.medi = lapply(tmp.medi, function(x) t(sapply(x, 
                                                     function(y) c(patient = y[1, 1], 
                                                                   state_timepoint = as.character(y[1, 2]), 
                                                                   apply(data.frame(y[, -c(1,2)]), 2, median)))))
    
    tmp.median = NULL
    for ( i in 1:length(tmp.medi) ) {
      tmp.median = rbind(tmp.median, tmp.medi[[i]])
    }
    
    colnames(tmp.median)[-c(1, 2)] = colnames(tmp.median.cond)[-1]
    
    tmp.median = data.frame(tmp.median, check.names = F)
    
    tmp.diffmods = lapply(mods.genes.corr[[grp]], names)
    tmp.genes = NULL
    for ( i in 1:length(tmp.diffmods) ) {
      tmp.genes = rbind(tmp.genes, cbind(M = names(tmp.diffmods)[i], G = tmp.diffmods[[i]]))
    }
    
    cell_object.geneclusts.time.final = lapply(1:nrow(tmp.genes), function(j) {
      g = as.character(tmp.genes[j, 'G'])
      tmp.median[[g]] = as.numeric(tmp.median[[g]])
      
      tmp.data.g = data.frame(state_timepoint = tmp.data$state_timepoint, gene = tmp.data[[g]])
      tmp.median.cond.g = data.frame(state_timepoint = tmp.median.cond$state_timepoint, gene = tmp.median.cond[[g]])
      tmp.median.g = data.frame(state_timepoint = tmp.median$state_timepoint, gene = tmp.median[[g]])
      
      tmp.colors = my.colors2[sort(unique(wnv@meta.data$state_timepoint)) %in% tmp.median.g$state_timepoint]
      p = ggplot(tmp.data.g, aes_string(x = 'state_timepoint', y = 'gene', fill = 'state_timepoint')) +
        geom_violin() + xlab('') + ylab('Expression') + 
        ggtitle(paste0(grp, '\n', tmp.genes[j, 'M'], ': ', g)) + 
        scale_fill_manual(values = tmp.colors) + 
        theme(plot.title = element_text(size = 30, hjust = 0.5), legend.position = 'none', 
              axis.text = element_text(size = 20), axis.text.x = element_text(angle = 90), 
              axis.title = element_text(size = 20)) + 
        geom_point(tmp.median.cond.g, mapping = aes_string(x = 'state_timepoint', y = 'gene'), 
                   shape = 17, fill = 'black', color = 'black', size = 6) + 
        geom_point(tmp.median.g, mapping = aes_string(x = 'state_timepoint', y = 'gene'), 
                   shape = 1, fill = 'black', color = 'darkred', size = 6)
      
      ggsave(filename = paste0('figures/bcr.signaling/diffmods.p0.01.with.sample.medians.v1.d211221.', 
                               grp, '.', tmp.genes[j, 'M'], '.', g, '.png'), plot = p, width = 7, height = 7)
      
      return(p)
    })
    
    plots.list = c(plots.list, cell_object.geneclusts.time.final)
  }
  
  pdf('figures/bcr.signaling/diffmods.p0.01.with.sample.medians.v1.d211221.pdf')
  invisible(lapply(plots.list, print))
  dev.off()
})
# user  system elapsed 
#4.363   0.259   4.729

system.time({
  (load('cellgroups.bycelltype.seuratobj.d211015.rdata'))
  # [1] "group.list"
  
  (celltypes.all = names(group.list))
})
#   user  system elapsed 
#109.807   9.523 121.008

{
  system.time({
    bcr.expr.mtx = list()
    for ( grp in celltypes.all ) {
      message(grp)
      
      tmp.metadt = cbind(group.list[[grp]]@meta.data[, c('U19.ID', 'state_timepoint')], 
                         t(data.frame(group.list[[grp]]@assays$RNA@data)[bcr.genes, 
                                                                         rownames(group.list[[grp]]@meta.data)]))
      
      tmp1 = table(tmp.metadt$state_timepoint)
      tmp2 = names(which(tmp1 >= 10))
      
      bcr.expr.mtx[[grp]] = tmp.metadt[tmp.metadt$state_timepoint %in% tmp2, ]
    }
    
    save(bcr.expr.mtx, file = 'bcr.signaling.genes.expr.mtx.d211221.rdata')
  })
  #   user  system elapsed 
  # 59.154  31.161  94.773
}

sapply(bcr.expr.mtx, dim)

system.time({
  plots.list = list()
  for ( grp in celltypes.all ) {
    message(grp)
    
    tmp.data = bcr.expr.mtx[[grp]]

    tmp.medi = split(tmp.data, as.character(tmp.data$state_timepoint))
    
    tmp.median.cond = t(sapply(tmp.medi, function(x) apply(x[, -c(1, 2)], 2, median)))
    tmp.median.cond = data.frame(state_timepoint = rownames(tmp.median.cond), tmp.median.cond, 
                                 check.names = F)
    
    tmp.medi = lapply(tmp.medi, function(x) split(x, as.character(x$U19.ID)))
    tmp.medi = lapply(tmp.medi, function(x) t(sapply(x, 
                                                     function(y) c(patient = y[1, 1], 
                                                                   state_timepoint = as.character(y[1, 2]), 
                                                                   apply(data.frame(y[, -c(1,2)]), 2, median)))))
    
    tmp.median = NULL
    for ( i in 1:length(tmp.medi) ) {
      tmp.median = rbind(tmp.median, tmp.medi[[i]])
    }
    
    tmp.median = data.frame(tmp.median, check.names = F)
    
    cell_object.geneclusts.time.final = lapply(bcr.genes, function(g) {
      tmp.median[[g]] = as.numeric(tmp.median[[g]])
      
      tmp.data.g = data.frame(state_timepoint = tmp.data$state_timepoint, gene = tmp.data[[g]])
      tmp.median.cond.g = data.frame(state_timepoint = tmp.median.cond$state_timepoint, gene = tmp.median.cond[[g]])
      tmp.median.g = data.frame(state_timepoint = tmp.median$state_timepoint, gene = tmp.median[[g]])
      
      tmp.colors = my.colors2[sort(unique(wnv@meta.data$state_timepoint)) %in% tmp.median.g$state_timepoint]
      p = ggplot(tmp.data.g, aes_string(x = 'state_timepoint', y = 'gene', fill = 'state_timepoint')) +
        geom_violin() + xlab('') + ylab('Expression') + 
        ggtitle(paste0(grp, '\n', g)) + 
        scale_fill_manual(values = tmp.colors) + 
        theme(plot.title = element_text(size = 30, hjust = 0.5), legend.position = 'none', 
              axis.text = element_text(size = 20), axis.text.x = element_text(angle = 90), 
              axis.title = element_text(size = 20)) + 
        geom_point(tmp.median.cond.g, mapping = aes_string(x = 'state_timepoint', y = 'gene'), 
                   shape = 17, fill = 'black', color = 'black', size = 6) + 
        geom_point(tmp.median.g, mapping = aes_string(x = 'state_timepoint', y = 'gene'), 
                   shape = 1, fill = 'black', color = 'darkred', size = 6)
      
      ggsave(filename = paste0('figures/bcr.signaling/', grp, '.', g, '.with.sample.medians.d211221.png'), 
                               plot = p, width = 7, height = 7)
      
      return(p)
    })
    
    plots.list = c(plots.list, cell_object.geneclusts.time.final)
  }
  
  pdf('figures/bcr.signaling/cell.types.with.sample.medians.v1.d211221.pdf')
  invisible(lapply(plots.list, print))
  dev.off()
})
#   user  system elapsed 
#100.264   6.099 112.032

{ ### DEA by Wilcoxon for B cells ###
  bcells = bcr.expr.mtx$B_Cells
  dim(bcells)
  #[1] 5654   24
  head(bcells)
  
  test.pairs = rbind(c('T1_WNA', 'T1_WNE'), c('T1_WNA', 'T1_WNF'), c('T1_WNE', 'T1_WNF'), 
                     c('T2_WNA', 'T2_WNE'), c('T2_WNA', 'T2_WNF'), c('T2_WNE', 'T2_WNF'), 
                     c('T3_WNA', 'T3_WNE'), c('T3_WNA', 'T3_WNF'), c('T3_WNE', 'T3_WNF'), 
                     c('T1_WNA', 'T2_WNA'), c('T1_WNA', 'T3_WNA'), c('T2_WNA', 'T3_WNA'), 
                     c('T1_WNE', 'T2_WNE'), c('T1_WNE', 'T3_WNE'), c('T2_WNE', 'T3_WNE'), 
                     c('T1_WNF', 'T2_WNF'), c('T1_WNF', 'T3_WNF'), c('T2_WNF', 'T3_WNF'))
  
  bc.test = NULL
  for ( i in 3:ncol(bcells) ) {
    gene = colnames(bcells)[i]
    
    # Get expression levels separated by group
    expr.by.grp = split(bcells[, i], bcells$state_timepoint)
    expr.by.grp = expr.by.grp[sapply(expr.by.grp, length) >= 10]
    
    # Group size (#cells)
    grsz = t(apply(test.pairs, 1, function(y) 
      if ( all(y %in% names(expr.by.grp)) ) {
        c(length(expr.by.grp[[y[1]]]), length(expr.by.grp[[y[2]]]))
      } else c(NA, NA)
    ))
    
    # Wilcoxon test for the predefined group pairs
    wc.p = apply(test.pairs, 1, function(y) 
      if ( all(y %in% names(expr.by.grp)) ) {
        wilcox.test(expr.by.grp[[y[1]]], expr.by.grp[[y[2]]])$p.value
      } else NA
    )
    
    # Fold change
    fcab = t(apply(test.pairs, 1, function(y) 
      if ( all(y %in% names(expr.by.grp)) ) {
        tmp.mean1 = mean(expr.by.grp[[y[1]]])
        tmp.mean2 = mean(expr.by.grp[[y[2]]])
        tmp.fcab = log2(tmp.mean1 / tmp.mean2)
        c(tmp.mean1, tmp.mean2, tmp.fcab)
      } else rep(NA, 3)
    ))
    
    test.out = cbind(gene, test.pairs, grsz, wc.p, fcab)
    colnames(test.out) = c('Gene', 'Group1', 'Group2', 'Size1', 'Size2', 
                           'PvalWC', 'Mean1', 'Mean2', 'log2FC')
    
    bc.test = rbind(bc.test, data.frame(test.out))
  }
  
  dim(bc.test)
  #[1] 396   9
  head(bc.test)
  #View(bc.test)
  
  write.csv(bc.test, file = 'bcr.signaling.genes.bcells.dea.wilcoxon.d220305.csv', quote = F, row.names = F)
  
  bc.test[is.nan(as.numeric(bc.test$PvalWC)), ]
  "###
       Gene Group1 Group2 Size1 Size2 PvalWC Mean1 Mean2 log2FC
  22 CAMK2A T2_WNA T2_WNE   661   538    NaN     0     0    NaN
  23 CAMK2A T2_WNA T2_WNF   661  1214    NaN     0     0    NaN
  24 CAMK2A T2_WNE T2_WNF   538  1214    NaN     0     0    NaN
  25 CAMK2A T3_WNA T3_WNE   524    31    NaN     0     0    NaN
  30 CAMK2A T2_WNA T3_WNA   661   524    NaN     0     0    NaN
  33 CAMK2A T2_WNE T3_WNE   538    31    NaN     0     0    NaN
  "###
  
  plot(bcr.expr.mtx$B_Cells$CAMK2A)
  
  summary(as.numeric(bc.test$PvalWC))
  #    Min.  1st Qu.   Median     Mean  3rd Qu.     Max.     NA's 
  #0.000000 0.001311 0.119253 0.266765 0.501734 0.992948        6
  
  hist(as.numeric(bc.test$PvalWC))
  plot(sort(as.numeric(bc.test$PvalWC)))
  
  (bc.test.signi1 = table(bc.test$Gene[as.numeric(bc.test$PvalWC) < 0.01]))
  length(bc.test.signi1)
  #[1] 21
  bcr.genes[!bcr.genes %in% names(bc.test.signi1)]
  #[1] "IKBKG"
  
  (bc.test.signi2 = table(bc.test$Gene[as.numeric(bc.test$PvalWC) < 0.001]))
  length(bc.test.signi2)
  #[1] 16
  bcr.genes[!bcr.genes %in% names(bc.test.signi2)]
  #[1] "PIP5K1A" "VAV2"    "BCL10"   "CARD11"  "IKBKG"   "PIP5K1C"
  
  (bc.test.signi3 = table(bc.test$Gene[as.numeric(bc.test$PvalWC) < 0.05]))
  length(bc.test.signi3)
  #[1] 22
  
  bc.test[as.numeric(bc.test$PvalWC) < 0.01, ]
}


{ ### DEA by Wilcoxon for plasma B cells ###
  plasma.b = bcr.expr.mtx$Plasma_B_Cells
  dim(plasma.b)
  #[1] 102   24
  head(plasma.b)
  
  test.pairs = rbind(c('T1_WNA', 'T1_WNE'), c('T1_WNA', 'T1_WNF'), c('T1_WNE', 'T1_WNF'), 
                     c('T2_WNA', 'T2_WNE'), c('T2_WNA', 'T2_WNF'), c('T2_WNE', 'T2_WNF'), 
                     c('T3_WNA', 'T3_WNE'), c('T3_WNA', 'T3_WNF'), c('T3_WNE', 'T3_WNF'), 
                     c('T1_WNA', 'T2_WNA'), c('T1_WNA', 'T3_WNA'), c('T2_WNA', 'T3_WNA'), 
                     c('T1_WNE', 'T2_WNE'), c('T1_WNE', 'T3_WNE'), c('T2_WNE', 'T3_WNE'), 
                     c('T1_WNF', 'T2_WNF'), c('T1_WNF', 'T3_WNF'), c('T2_WNF', 'T3_WNF'))
  
  pb.test = NULL
  for ( i in 3:ncol(plasma.b) ) {
    gene = colnames(plasma.b)[i]
    
    # Get expression levels separated by group
    expr.by.grp = split(plasma.b[, i], plasma.b$state_timepoint)
    expr.by.grp = expr.by.grp[sapply(expr.by.grp, length) >= 10]
    
    # Group size (#cells)
    grsz = t(apply(test.pairs, 1, function(y) 
      if ( all(y %in% names(expr.by.grp)) ) {
        c(length(expr.by.grp[[y[1]]]), length(expr.by.grp[[y[2]]]))
      } else c(NA, NA)
    ))
    
    # Wilcoxon test for the predefined group pairs
    wc.p = apply(test.pairs, 1, function(y) 
      if ( all(y %in% names(expr.by.grp)) ) {
        wilcox.test(expr.by.grp[[y[1]]], expr.by.grp[[y[2]]])$p.value
      } else NA
    )
    
    # Fold change
    fcab = t(apply(test.pairs, 1, function(y) 
      if ( all(y %in% names(expr.by.grp)) ) {
        tmp.mean1 = mean(expr.by.grp[[y[1]]])
        tmp.mean2 = mean(expr.by.grp[[y[2]]])
        tmp.fcab = log2(tmp.mean1 / tmp.mean2)
        c(tmp.mean1, tmp.mean2, tmp.fcab)
      } else rep(NA, 3)
    ))
    
    test.out = cbind(gene, test.pairs, grsz, wc.p, fcab)
    colnames(test.out) = c('Gene', 'Group1', 'Group2', 'Size1', 'Size2', 
                           'PvalWC', 'Mean1', 'Mean2', 'log2FC')
    
    pb.test = rbind(pb.test, data.frame(test.out))
  }
  
  dim(pb.test)
  #[1] 396   9
  head(pb.test)
  #View(pb.test)
  
  write.csv(pb.test, file = 'bcr.signaling.genes.plasmab.dea.wilcoxon.d220307.csv', quote = F, row.names = F)
  
  pb.test[is.nan(as.numeric(pb.test$PvalWC)), ]
  "###
         Gene Group1 Group2 Size1 Size2 PvalWC Mean1 Mean2 log2FC
  22   CAMK2A T2_WNA T2_WNE    11    10    NaN     0     0    NaN
  23   CAMK2A T2_WNA T2_WNF    11    10    NaN     0     0    NaN
  24   CAMK2A T2_WNE T2_WNF    10    10    NaN     0     0    NaN
  28   CAMK2A T1_WNA T2_WNA    71    11    NaN     0     0    NaN
  58  PIP5K1A T2_WNA T2_WNE    11    10    NaN     0     0    NaN
  222  NFATC2 T2_WNE T2_WNF    10    10    NaN     0     0    NaN
  292   IKBKG T2_WNA T2_WNE    11    10    NaN     0     0    NaN
  293   IKBKG T2_WNA T2_WNF    11    10    NaN     0     0    NaN
  294   IKBKG T2_WNE T2_WNF    10    10    NaN     0     0    NaN
  "###
  
  plot(bcr.expr.mtx$Plasma_B_Cells$CAMK2A)
  plot(bcr.expr.mtx$Plasma_B_Cells$PIP5K1A)
  plot(bcr.expr.mtx$Plasma_B_Cells$NFATC2)
  plot(bcr.expr.mtx$Plasma_B_Cells$IKBKG)
  
  summary(as.numeric(pb.test$PvalWC))
  #  Min. 1st Qu.  Median    Mean 3rd Qu.    Max.    NA's 
  #0.0014  0.1787  0.3681  0.4307  0.6965  1.0000     317
  
  hist(as.numeric(pb.test$PvalWC))
  plot(sort(as.numeric(pb.test$PvalWC)))
  
  (pb.test.signi1 = table(pb.test$Gene[as.numeric(pb.test$PvalWC) < 0.01]))
  #HCLS1 PPP3CA 
  #    1      1
  length(pb.test.signi1)
  #[1] 2
  bcr.genes[!bcr.genes %in% names(pb.test.signi1)]
  
  (pb.test.signi2 = table(pb.test$Gene[as.numeric(pb.test$PvalWC) < 0.001]))
  length(pb.test.signi2)
  #[1] 0

  (pb.test.signi3 = table(pb.test$Gene[as.numeric(pb.test$PvalWC) < 0.05]))
  #CARD11   ETS1  HCLS1   IRF4    LYN PPP3CA  PRKCB   VAV1 
  #     1      1      1      1      1      2      1      1
  length(pb.test.signi3)
  #[1] 8
  
  pb.test[which(as.numeric(pb.test$PvalWC) < 0.05), ]
  "###
        Gene Group1 Group2 Size1 Size2              PvalWC             Mean1             Mean2             log2FC
  118   IRF4 T1_WNA T2_WNA    71    11  0.0205845205878343 0.602232490636916   1.6471750811076  -1.45160146064545
  131    LYN T2_WNA T2_WNF    11    10  0.0133299453664207 0.996815262303272 0.346996031788971   1.52240699334321
  184 CARD11 T2_WNA T2_WNE    11    10  0.0452130990123186 0.315804243066984                 0                Inf
  239   VAV1 T2_WNA T2_WNF    11    10  0.0483514750335888 0.744666407183688 0.108793008527275   2.77500843104408
  256 PPP3CA T2_WNA T2_WNE    11    10 0.00145757257905104 0.708380049712895                 0                Inf
  257 PPP3CA T2_WNA T2_WNF    11    10  0.0364698303560265 0.708380049712895 0.243528418553327   1.54043344421933
  316  HCLS1 T1_WNA T2_WNA    71    11 0.00137220039014406 0.744537294114912  1.42485492419624 -0.936399013713983
  352   ETS1 T1_WNA T2_WNA    71    11  0.0242476769720366 0.880381100460175  1.36945713812128  -0.63740403295205
  370  PRKCB T1_WNA T2_WNA    71    11  0.0131879338257186 0.468620958357308  1.04845221262943  -1.16176772329003
  "###
}
```
