## Supplemental Software for "Early cellular and molecular signatures correlate with severity of West Nile Virus infection": README.txt

##### West Nile Virus: Seq-Well data analysis #####

1. System requirements
  1.1. All software dependencies and operating systems (including version numbers)
      Seurat: 3.1.5 or 4.0.1
      WGCNA: 1.71
      nichenetr: 1.0.0
      Platform: x86_64-apple-darwin13.4.0 (64-bit, macOS Catalina 10.15.7)

  1.2. Versions the software has been tested on
      R: 4.0.3, 4.0.5, 4.1.3
      Jupyter Notebook: 6.0.1

  1.3. Any required non-standard hardware
       : None

2. Installation guide
  2.1. Instructions
      (1) Install R and Jupyter Notebook with IRKernel:
          https://www.r-project.org/
          https://jupyter.org/
          https://irkernel.github.io/
      (2) Install the following 4 main packages:
          https://satijalab.org/seurat/
          https://horvath.genetics.ucla.edu/html/CoexpressionNetwork/Rpackages/WGCNA/
          https://github.com/saeyslab/nichenetr

  2.2. Typical install time on a "normal" desktop computer
       : 1 hour

3. Demo
  3.1. Instructions to run on data
       (1) For the analysis of WGCNA-based differential modules, run the code of "code.wgcna.groups.demo.R" using the demo data file of "cellgroups.bycelltype.seuratobj.d211015.sub1.rdata", which contains the processed data of plasma B cells from the full analysis ("code.wgcna.groups_copy.R").
       (2) For NicheNet, see the NicheNet vignette,
           https://github.com/saeyslab/nichenetr/blob/master/vignettes/ligand_activity_geneset.md
  
  3.2. Expected output
       : 7 differential modules in a list object, "diffmods", in "code.wgcna.groups.demo.R"
  
  3.3. Expected run time for demo on a "normal" desktop computer
       : 3 min

4. Instructions for use
  4.1. How to run the software on your data
       : Run the following 3 code files in sequence using Seq-Well DGE data files as input files, "WNV_scRNA-seq_Processing_HL_copy.ipynb" --> "code.wgcna.groups_copy.R" --> "wnv.nichenet.wgcna.v3_copy.R"
  
  4.2. Reproduction instructions
       : Given the full raw DGE data from Seq-Well experiments, one can run "WNV_scRNA-seq_Processing_HL_copy.ipynb" to generate the processed data with cell type identification, "wnv.seqwell.pbmcs.seurat.d211011.Robj", which is used for the downstream analyses of WGCNA-based differential module analysis ("code.wgcna.groups_copy.R") and NicheNet analysis ("wnv.nichenet.wgcna.v3_copy.R"). The following files are the main results about the Seq-Well data in the manuscript:
        (1) 47 WGCNA-based differential modules ("code.wgcna.groups_copy.R")
            : "fig3b.diffmods.heatmap.v1.d211015.png"
        (2) BCR signaling genes in B cells and plasma B cells for MOFA factor 1 ("code.wgcna.groups_copy.R")
            : "bcr.signaling.genes.bcells.dea.wilcoxon.d220305.csv" and "bcr.signaling.genes.plasmab.dea.wilcoxon.d220307.csv"
        (3) Top ligand-target pairs by Nichenet analysis ("wnv.nichenet.wgcna.v3.demo.R")
            : "wnv.nichenet.wgcna.top.lig.tg.pairs.d211025.csv"
