## Supplemental Software for "Early cellular and molecular signatures correlate with severity of West Nile Virus infection": wnv.nichenet.wgcna.v3_copy.R

```
setwd("~/Box Sync/projects/yale/smita/HIPC/analysis/WNV.SeqWell/")

date()

system('which R', intern=T)
#[1] "/usr/local/bin/R"
system('R --version', intern=T)
#[1] "R version 4.0.3 (2020-10-10) -- \"Bunny-Wunnies Freak Out\""

library(Seurat)
library(nichenetr)
library(limma)
library(multtest)
library(textshape)
library(tidyverse)
library(plyr)
library(dplyr)
library(gridExtra) # for grid.arrange()

packageVersion('Seurat')
# [1] ‘4.0.1’
packageVersion('nichenetr')
# [1] ‘1.0.0’


##### Load the Seurat object and set the identities to cell types #####
system.time({
    wnv = get(load('wnv.seqwell.pbmcs.seurat.d211011.Robj'))
    rm(wnv_cleaned)
})
#    user  system elapsed 
# 149.147  12.512 166.460

wnv = SetIdent(wnv, value = 'new_cell_types')
levels(wnv)
#[1] "CD14+_Monocytes"         "CD16+_Monocytes"         "cDCs"                    "CD4+_T_Cells"           
#[5] "Proliferating_T_Cells"   "pDCs"                    "Plasma_B_Cells"          "NK_Cells"               
#[9] "B_Cells"                 "CD8+_T_Cells"            "Regulatory_CD4+_T_Cells"

table(as.character(wnv@meta.data$new_cell_types))
#       B_Cells         CD14+_Monocytes         CD16+_Monocytes            CD4+_T_Cells 
#          5654                   19440                    1780                   15442 
#  CD8+_T_Cells                    cDCs                NK_Cells                    pDCs 
#          6996                    1534                    4424                     364 
#Plasma_B_Cells   Proliferating_T_Cells Regulatory_CD4+_T_Cells 
#           118                     510                     596

par(omi = c(0, 2.2, 0, 0), cex = 1.5)
barplot(sort(table(as.character(wnv@meta.data$new_cell_types))), horiz = T, las = 2, 
        main = 'Number of cells')
# 'cell.numbers.each.celltype.barplot.v1.png'


##### Load the data of the NicheNet networks #####
ligand_target_matrix = readRDS("nichenet.data/ligand_target_matrix.rds")
dim(ligand_target_matrix) # targets by ligands
#[1] 25345   688
ligand_target_matrix[1:3, 1:3]

lr_network = readRDS("nichenet.data/lr_network.rds")
dim(lr_network)
#[1] 12651     4
head(lr_network)

apply(lr_network, 2, function(x) length(unique(x)))
# from       to   source database 
#  688      857       14        5

table(lr_network$database)
table(lr_network$source)

receptors = lr_network %>% pull(to) %>% unique()
length(receptors)
# [1] 857

# If wanted, users can remove ligand-receptor interactions that were predicted based on 
# protein-protein interactions and only keep ligand-receptor interactions that are 
# described in curated databases. 
lr_network_ppi = lr_network %>% filter(database != "ppi_prediction_go" & database != "ppi_prediction")
dim(lr_network_ppi)
#[1] 2022    4

system.time({
    weighted_networks = readRDS("nichenet.data/weighted_networks.rds")
})
# user  system elapsed 
#3.553   0.045   3.607

sapply(weighted_networks, dim)
#      lr_sig      gr
#[1,] 2476577 2912482
#[2,]       3       3
lapply(weighted_networks, head)

weighted_networks_lr = weighted_networks$lr_sig %>% inner_join(lr_network %>% distinct(from,to), 
                                                               by = c("from", "to"))
dim(weighted_networks_lr)
#[1] 12019     3
head(weighted_networks_lr)

apply(weighted_networks_lr, 2, function(x) length(unique(x)))
# from     to weight 
#  688    857   3165

{ ### Example of APP ligand and APOBEC3A target
    ligand_target_matrix['TNFRSF21', 'APP']
    # [1] 0.0006932006
    
    ligand_target_matrix['APOBEC3A', 'APP']
    # [1] 0.009249296
    
    lr_network[lr_network$from == 'APP', ]
    apply(lr_network[lr_network$from == 'APP', ], 2, function(x) length(unique(x)))
    # from       to   source database 
    #    1      136        4        3
    
    lr_network[lr_network$from == 'APP' & lr_network$to == 'TNFRSF21', ]
    # APP   TNFRSF21 ppi_lr ppi_prediction
    
    weighted_networks$lr_sig[weighted_networks$lr_sig$from == 'APP' & 
                                 weighted_networks$lr_sig$to == 'TNFRSF21', ]
    #APP   TNFRSF21  0.964
    
    weighted_networks$lr_sig[weighted_networks$lr_sig$from == 'TNFRSF21' & 
                                 weighted_networks$lr_sig$to == 'APOBEC3A', ]
    
    (lr.tmp = weighted_networks$lr_sig[weighted_networks$lr_sig$from == 'APP', ])
    (gr.tmp = weighted_networks$gr[weighted_networks$gr$to == 'APOBEC3A', ])
    
    (tmp = intersect(lr.tmp$to, gr.tmp$from))
    
    (gr.tmp = gr.tmp[gr.tmp$from %in% tmp, ])
    (sg.tmp = weighted_networks$lr_sig[weighted_networks$lr_sig$to %in% tmp, ])
    (lr.tmp = lr_network[lr_network$from == 'APP' & lr_network$to %in% sg.tmp$from, ])
}


##### Run upstream ligand analysis #####
## Get expressed genes in sender cells (at least 10% of each population)
## : lapply to get the expressed genes of every sender cell type separately

system.time({
    list_expressed_genes_sender = unique(wnv@meta.data$new_cell_types) %>% lapply(get_expressed_genes, wnv, 0.10)
    expressed_genes_sender = unique(unlist(list_expressed_genes_sender))
})
#   user  system elapsed 
#194.868  33.601 231.422

length(expressed_genes_sender)
#[1] 9056

ligands = lr_network %>% pull(from) %>% unique()
expressed_ligands = intersect(ligands, expressed_genes_sender)
length(expressed_ligands)
# [1] 148

targets.cty.mods = rbind(c('CD14+_Monocytes', 'MOD1'), 
                         c('CD16+_Monocytes', 'MOD1'), c('CD16+_Monocytes', 'MOD2'), c('CD16+_Monocytes', 'MOD3'), 
                         c('CD4+_T_Cells', 'MOD2'), c('CD8+_T_Cells', 'MOD1'), c('CD8+_T_Cells', 'MOD3'), 
                         c('cDCs', 'MOD1'), c('cDCs', 'MOD4'), c('pDCs', 'MOD1'), 
                         c('NK_Cells', 'MOD3'), c('Plasma_B_Cells', 'MOD6'))

N.TOP.LIGANDS = 20
ACTIVITY.THRES = .1
system.time({
    niches.cty.mods = list()
    plots.list = list()
    for ( i in 1:nrow(targets.cty.mods) ) {
        tmp.receiver = targets.cty.mods[i, 1]
        tmp.module = targets.cty.mods[i, 2]
        
        message('### ', tmp.receiver, ': ', tmp.module, ' ###')
        
        ## Get background expressed genes in the receiver cells (at least 10% of population)
        rec_object = subset(wnv, idents = tmp.receiver)
        expressed_genes_receiver = get_expressed_genes(tmp.receiver, rec_object, 0.1)
        background_expressed_genes = expressed_genes_receiver %>% .[. %in% rownames(ligand_target_matrix)]
        
        message("#Background genes in the receiver cells = ", length(background_expressed_genes))
        
        ## Genes of interest in the WGCNA differential module
        diffmods = read.csv(paste0(tmp.receiver, '.final.p0.01.diffmods.d211015.csv'))[, tmp.module]
        geneset_oi = diffmods[!is.na(diffmods)]
        
        message("#Genes of interest = ", length(geneset_oi))
        
        ## Expressed receptors in the receiver cells
        expressed_receptors = intersect(receptors, expressed_genes_receiver)
        message("#expressed receptors = ", length(expressed_receptors))
        
        ## Get those ligand-receptor pairs from the ligand-receptor network where both are expressed
        lr_network_expressed = lr_network[lr_network$from %in% expressed_ligands & 
                                          lr_network$to %in% expressed_receptors, ]

        ## Get our potential ligands from those pairings
        potential_ligands = unique(lr_network_expressed$from)
        
        message("#Potential ligands = ", length(potential_ligands))
        message("#Potential receptors = ", length(unique(lr_network_expressed$to)))
        
        ## Using our geneset of interest, our background expressed genes (in the receiver), our ligand-target matrix, 
        ## and our set of potential ligands, calculate ligand activities in the receiver using the matrix weight values.
        ## The base statistic is the pearson correlation.
        ligand_activities = predict_ligand_activities(geneset = geneset_oi, 
                                                      background_expressed_genes = background_expressed_genes, 
                                                      ligand_target_matrix = ligand_target_matrix, 
                                                      potential_ligands = potential_ligands)
        
        ## Rank ligand activities in order of pearson correlation and get the N.TOP.LIGANDS upstream ligands
        best_upstream_ligands = ligand_activities %>% top_n(N.TOP.LIGANDS, pearson) %>% arrange(-pearson) %>% pull(test_ligand)
        
        message("Top 5 upstream ligands: ", paste(head(best_upstream_ligands)[1:5], collapse = ' '))
        
        ## Show a histogram of ligand activity scores for the N.TOP.LIGANDS genes
        p_hist_lig_activity = ggplot(ligand_activities, aes(x = pearson)) + 
            geom_histogram(color = "black", fill = "lightgrey", bins = 10) + 
            geom_vline(aes(xintercept = min(ligand_activities %>% top_n(N.TOP.LIGANDS, pearson) %>% pull(pearson))), 
                       color = "red", linetype = "dashed", size = 1) + 
            labs(x = "ligand activity (PCC)", y = "#ligands") +
            theme(axis.title = element_text(size = 20, family = "Arial"), 
                  axis.text.x = element_text(face = "italic", size = 20), 
                  axis.text.y = element_text(size = 20))
        #theme_classic()
        
        print(p_hist_lig_activity)
        
        ## Get weighted target links from the top upstream ligands
        active_ligand_target_links_df = best_upstream_ligands %>% lapply(get_weighted_ligand_target_links,
                                                                         geneset = geneset_oi, 
                                                                         ligand_target_matrix = ligand_target_matrix,
                                                                         n = 250) %>% bind_rows()
        active_ligand_target_links_df = active_ligand_target_links_df[!is.na(active_ligand_target_links_df$weight), ]
        
        message("#Active ligand-target links: ", nrow(active_ligand_target_links_df))
        
        if ( nrow(active_ligand_target_links_df) == 0 ) next
        
        ## Prepare the ligand target visualization
        active_ligand_target_links = prepare_ligand_target_visualization(ligand_target_df = active_ligand_target_links_df, 
                                                                         ligand_target_matrix = ligand_target_matrix,
                                                                         cutoff = ACTIVITY.THRES)
        
        ## Find an overlap of the top upstream ligands and our ligand activity score matrix
        order_ligands = rev(intersect(best_upstream_ligands, colnames(active_ligand_target_links)))
        
        ## Cluster our ligand activities
        if ( nrow(active_ligand_target_links) > 1 ) {
            active_ligand_target_links = cluster_matrix(x = active_ligand_target_links[, order_ligands], dim = 'row')        
        } else {
            active_ligand_target_links = data.frame(active_ligand_target_links)[, order_ligands]        
        }
        
        ## Order the targets of ligands by activity
        order_targets = rev(rownames(active_ligand_target_links))
        
        ## Order the matrix by ligand order
        vis_ligand_target = t(active_ligand_target_links[order_targets, order_ligands])
        
        message("#Best active ligands with activity > 0.1: ", length(order_ligands))
        message("#Best active targets with activity > 0.1 : ", length(order_targets))
        
        ## Heatmap of the top upstream ligands for target genes
        p.activity = vis_ligand_target %>% make_heatmap_ggplot("Best Upstream Ligands", "Ligand Targets", 
                                                               color = "blue", 
                                                               x_axis_position = "top",
                                                               legend_position = "right", 
                                                               legend_title = "Regulatory potential") + 
                    #scale_fill_gradient2(low = "whitesmoke", high = "blue", breaks = c(0, 0.005, 0.01)) + 
                    theme(axis.title = element_text(size = 20), 
                          axis.text.x = element_text(face = "italic", size = 12), 
                          axis.text.y = element_text(size = 12))
        
        ## Dot plot of the top upstream ligand expression across all cell types
        p.ligand = DotPlot(wnv, features = rev(order_ligands), cols = c('black', 'red')) + 
                    RotatedAxis() + 
                    labs(x = 'Best Upstream Ligands', y = 'Cell Types') + 
                    theme(axis.title = element_text(size = 20), 
                          axis.text.x = element_text(size = 12), 
                          axis.text.y = element_text(size = 15))# + NoLegend()
        
        
        ## Get the ligand-receptor network of the top-ranked ligands
        lr_network_top = lr_network %>% 
            filter(from %in% best_upstream_ligands & to %in% expressed_receptors) %>% 
            distinct(from, to)
        best_receptors = lr_network_top %>% pull(to) %>% unique()
        
        message("#Best receptors : ", length(best_receptors))
        
        active_ligand_receptor_links_df = weighted_networks_lr %>% 
            filter(from %in% best_upstream_ligands & to %in% best_receptors) %>% 
            filter(weight > quantile(weighted_networks_lr$weight, .5)) #%>% rename(ligand = from, receptor = to)
        colnames(active_ligand_receptor_links_df)[1:2] = c('ligand', 'receptor')
        
        message("#Active ligand-receptor links: ", nrow(active_ligand_receptor_links_df))
        
        ## Prepare the ligand receptor visualization
        active_ligand_receptor_links = matrix(0, 
                                              ncol = length(unique(active_ligand_receptor_links_df$ligand)), 
                                              nrow = length(unique(active_ligand_receptor_links_df$receptor)))
        colnames(active_ligand_receptor_links) = unique(active_ligand_receptor_links_df$ligand)
        rownames(active_ligand_receptor_links) = unique(active_ligand_receptor_links_df$receptor)
        for ( i in 1:nrow(active_ligand_receptor_links_df) ) {
            active_ligand_receptor_links[active_ligand_receptor_links_df$receptor[i], 
                                         active_ligand_receptor_links_df$ligand[i]] = active_ligand_receptor_links_df$weight[i]
        }
        
        ## Find an overlap of the top upstream ligands and our ligand activity score matrix
        order_ligands = intersect(order_ligands, colnames(active_ligand_receptor_links))
        #order_ligands = rev(intersect(best_upstream_ligands, colnames(active_ligand_receptor_links)))
        
        ## Cluster our ligand activities
        if ( nrow(active_ligand_receptor_links) > 1 ) {
            active_ligand_receptor_links = cluster_matrix(x = active_ligand_receptor_links[, order_ligands], dim = 'row')        
        } else {
            active_ligand_receptor_links = data.frame(active_ligand_receptor_links)[, order_ligands]        
        }
        
        ## Order the targets of ligands by activity
        order_receptors = rev(rownames(active_ligand_receptor_links))
        
        ## Order the matrix by ligand order
        vis_ligand_receptor = t(active_ligand_receptor_links[order_receptors, order_ligands])
        
        message("#Best active ligands with >50% weight: ", length(order_ligands))
        message("#Best active receptors with >50% weight : ", length(order_receptors))
        
        ## Heatmap of the top upstream ligands for receptors
        p.receptor = vis_ligand_receptor %>% make_heatmap_ggplot("Best Upstream Ligands", "Receptors", 
                                                                 color = "darkgreen", 
                                                                 x_axis_position = "top",
                                                                 legend_position = "right", 
                                                                 legend_title = "Weight") + 
            #scale_fill_gradient2(low = "whitesmoke", high = "blue", breaks = c(0, 0.005, 0.01)) + 
            theme(axis.title = element_text(size = 20), 
                  axis.text.x = element_text(face = "italic", size = 12), 
                  axis.text.y = element_text(size = 12))
        
        
        #https://stackoverflow.com/questions/17059099/saving-grid-arrange-plot-to-file
        #https://cran.r-project.org/web/packages/gridExtra/vignettes/arrangeGrob.html
        #https://cran.r-project.org/web/packages/egg/vignettes/Ecosystem.html
        ggsave(paste0('figures/nichenet.', tmp.receiver, '_', tmp.module, '.plots.v1.png'), limitsize = F, 
               plot = grid.arrange(p.activity, p.receptor, p.ligand, ncol = 1), # from gridExtra
               width = 12, height = 18)
        ggsave(paste0('figures/nichenet.', tmp.receiver, '_', tmp.module, '.plots.v2.png'), limitsize = F, 
               plot = grid.arrange(p.activity, p.receptor, p.ligand, nrow = 1), # from gridExtra
               width = 3 * max(ncol(vis_ligand_target), ncol(vis_ligand_receptor)) * .25, height = 6)
        
        ## Send back the ligand-target activity matrix
        tmp.output = list(Activity = list(LT = vis_ligand_target, LR = vis_ligand_receptor), 
                          Plot = p.activity + p.receptor + p.ligand)
        
        niches.cty.mods[[paste0(tmp.receiver, '|', tmp.module)]] = tmp.output$Activity
        plots.list = c(plots.list, tmp.output['Plot'])
    }
    
    save(niches.cty.mods, file = 'nichenet.wgcna.cty.mods.d211025.rdata')
    #save(plots.list, file = 'plots.rdata') # too big file > 100GB
})
#    user  system elapsed
# 379.828 122.506 587.528

{
    #https://stackoverflow.com/questions/17059099/saving-grid-arrange-plot-to-file
    pdf('figures/nichenet.cty.mods.plots.d211025.pdf', width = 40, height = 10)
    invisible(lapply(plots.list, print))
    dev.off()
}

#(load('nichenet.wgcna.cty.mods.d211025.rdata'))

sapply(niches.cty.mods, function(x) dim(x$LT))

lapply(niches.cty.mods, function(x) head(x$LT))


##### ligand-target activities #####
par(omi = c(1.2, 0, 0, 0), cex = 1.5, mfrow = c(1, 2))
barplot(sort(sapply(niches.cty.mods, function(x) mean(x$LT)), decreasing = T), 
        las = 2, main = 'Mean activity')
barplot(sort(sapply(niches.cty.mods, function(x) median(x$LT)), decreasing = T), 
        las = 2, main = 'Median activity')
# 'ligand.target.activities.each.module.v1.png'

par(omi = c(2, 0, 0, 0), cex = 1.5, mfrow = c(1, 2))
barplot(sort(sapply(niches.cty.mods, function(x) mean(x$LT[x$LT > 0])), decreasing = T), las = 2, 
                    main = 'Mean non-zero activity')
barplot(sort(sapply(niches.cty.mods, function(x) median(x$LT[x$LT > 0])), decreasing = T), las = 2, 
        main = 'Median non-zero activity')
# 'ligand.target.activities.each.module.v2.png'

(max.ind = sapply(niches.cty.mods, function(x) which(x$LT == max(x$LT), arr.ind = T)))

{
    top.pairs = sapply(1:length(niches.cty.mods), 
                       function(x) c(Ligand = rownames(niches.cty.mods[[x]]$LT)[max.ind[1, x]], 
                                     Target = colnames(niches.cty.mods[[x]]$LT)[max.ind[2, x]], 
                                     Activity = niches.cty.mods[[x]]$LT[max.ind[1, x], max.ind[2, x]]))
    
    top.pairs = rbind(names(niches.cty.mods), 
                      sapply(strsplit(names(niches.cty.mods), '[|]'), function(x) x), 
                      #matrix(unlist(strsplit(names(niches.cty.mods), '[|]')), byrow = T, ncol = 2)
                      top.pairs)
    
    rownames(top.pairs)[1:3] = c('ID', 'CellType', 'Module')
    
    top.pairs
    
    top.pairs = t(top.pairs[, order(as.numeric(top.pairs['Activity', ]), decreasing = T)])
    top.pairs

    write.csv(top.pairs, file = 'wnv.nichenet.wgcna.top.lig.tg.pairs.d211025.csv', 
              quote = F, row.names = F)
}


sessionInfo()

date()
```
