## Supplemental Software for "Early cellular and molecular signatures correlate with severity of West Nile Virus infection": WNV_scRNA-seq_Processing_HL_copy.ipynb

```
# %% [raw]
$ conda activate seurat3.1.5
$ unalias python
$ unalias pip
$ unset PYTHONPATH

# %% [code]
date()

# %% [markdown]
---

# %% [markdown]
# Initial U19 WNV Dataset Processing

# %% [code]
# Load important packages
library(Seurat)
library(ggplot2)
library(reshape2)
library(dplyr)
library('data.table')
library(stringr)

library(repr)
options(repr.plot.width=10, repr.plot.height=10)

# %% [code]
sessionInfo()

# %% [code]
set.seed(1996)

# %% [markdown]
## Read in data, clean up, and merge

# %% [code]
# Merge cell vs. gene matrices (three functions)
union = function(list.mat){
    rows = c()
    for(mat in list.mat){
        rows = c(rows, rownames(mat))
    }
    rows = unique(rows)
    return(rows)
}
fill.up.genes = function(rows, mat){
    new.rows = setdiff(rows, rownames(mat))
    num.rows = length(new.rows)
    add.mat = matrix(0, nrow=num.rows, ncol=ncol(mat))
    rownames(add.mat) = new.rows
    mat = rbind(mat, add.mat)
    mat = mat[order(rownames(mat)), ]
    return(mat)
}
complete.merge = function(list.mat){
    rows = union(list.mat)
    
    list.mat = lapply(list.mat, function(x){
        mat = fill.up.genes(rows, x)
        return(mat)
    })
    output = do.call(cbind, list.mat)
    return(output)
}

# Function to pick cells with 500 transcripts
filter.500 = function(mat){
  mat_filtered = mat[, which(unlist(lapply(1:(ncol(mat)), function(x) {sum(mat[, x])})) > 500)]
  return(mat_filtered)
}

# %% [code]
#list files
path.SeqWell = '/Users/hlee/Box Sync/WNV-SMART_tube_Share_folder/Seqwell from Ira Dropbox_WNV Alignments-2/DGEs_Apr-2020/'
list.files(path.SeqWell)

# %% [code]
# List of all DGEs
data.list.SeqWell = list(path.SeqWell)

# Get paths to DGEs
temp.df2 = data.frame(file.name = list.files(path.SeqWell, pattern = "dge.txt.gz"))

path.names = c()
for (i in list.files(path.SeqWell, pattern = "dge.txt.gz")){
  path.names = c(path.names, paste0(path.SeqWell, "/", i))
}

# Get filenames of DGEs
file.names = c()
for (i in list.files(path.SeqWell, pattern = "dge.txt.gz")){
  file.names = c(file.names, strsplit(as.character(i), "_dge")[[1]][1])
}

# Assemble dataframe with filenames, path names, and array names
temp.df2$path.name = path.names
temp.df2$array.name = file.names

temp.df2 = temp.df2[!grepl("unmapped", temp.df2$array.name), ]
temp.df2

# %% [markdown]
## Actually read in data

# %% [code]
system.time({
wnv.data.seqwell = lapply(split(temp.df2, seq(NROW(temp.df2))), 
                          function(x){
                              path = as.character(x$path.name)
                              message(path)
                              mat = as.matrix(read.table(path, header = T, row.names = 1))
                              rm(list = setdiff(ls(), "mat"))
                              return(mat)
                          })
})

# %% [code]
# Replace underscores with periods as required in Seurat
names(wnv.data.seqwell) = gsub("_", ".", temp.df2$array.name)

# %% [code]
# Add assay identifiers to all unique cell IDs
wnv.data.seqwell.labeled = lapply(seq_along(wnv.data.seqwell), function(i){
  colnames(wnv.data.seqwell[[i]]) = paste(colnames(wnv.data.seqwell[[i]]), "_", names(wnv.data.seqwell)[i], sep="")
  return(wnv.data.seqwell[[i]])
})

# %% [markdown]
## Merge assays

# %% [code]
# Merge all matrices
system.time({
    wnv.combined_runs.merged.mat = complete.merge(wnv.data.seqwell.labeled)
})

# %% [code]
# Dimensions of final merged dataset: rows = genes, columns = cells
dim(wnv.combined_runs.merged.mat)

# %% [code]
rm(wnv.data.seqwell.labeled)

# %% [markdown]
## Filter out cells with <500 transcripts

# %% [code]
system.time({
    wnv.combined_runs.merged.mat = filter.500(wnv.combined_runs.merged.mat)
})

# %% [code]
dim(wnv.combined_runs.merged.mat)

# %% [markdown]
## Create Seurat Object

# %% [code]
# Filter out cells with fewer than 300 genes and create a Seurat object
system.time({
    wnv = CreateSeuratObject(wnv.combined_runs.merged.mat, names.delim = "_", names.field = 2, min.features = 300)
})

# %% [code]
dim(wnv)

# %% [code]
unique(Idents(wnv))

# %% [code]
wnv@meta.data

# %% [code]
rm(wnv.combined_runs.merged.mat)

# %% [markdown]
## Visualize n-gene and n-UMI

# %% [code]
meta1 = wnv@meta.data
tran_num = meta1[, c(1,2)]
colnames(tran_num) = c('CELL_BARCODE', 'NUM')
tran_num$TYPE = 'Transcripts per Cell'

gene_num = meta1[, c(1,3)]
colnames(gene_num) = c('CELL_BARCODE','NUM')
gene_num$TYPE = 'Genes per Cell'

combined_geneumi = rbind(tran_num, gene_num)

# %% [code]
combined_plot = ggplot(combined_geneumi, aes(x=TYPE, y=NUM)) + 
  geom_violin(fill = 'lightblue3') + 
  scale_y_log10() + 
  theme(axis.text.x = element_text(angle = 90),
        legend.position = "none") + 
        ylab('Count per Cell') + xlab (NULL) + theme_minimal() 
    #ggtitle("Basic boxplot")

combined_plot

# %% [markdown]
## Add Metdata

# %% [code]
metadata = read.csv(paste0("/Users/hlee/Box Sync/WNV-SMART_tube_Share_folder/Seqwell from Ira Dropbox_WNV Alignments-2/Sample_Metadata/", 
                           "Metadata_for_Seurat_WNV_FINAL_corrected.csv"))
metadata

# %% [code]
for ( i in 1:ncol(metadata) ) {
    tmp1 = metadata[, i]
    if ( length(grep('Asymptomatic', tmp1)) > 0 ) {
        metadata[, i] = sub('Asymptomatic', 'WNA', tmp1)
    }
}

metadata

# %% [code]
metadata$orig.ident = metadata$filename

wnv@meta.data$orig.ident = paste0('Sample_', wnv@meta.data$orig.ident)

setequal(unique(wnv@meta.data$orig.ident), metadata$orig.ident)

# %% [code]
foo.metadt = merge(wnv@meta.data, metadata, by = 'orig.ident')
rownames(foo.metadt) = rownames(wnv@meta.data)

wnv@meta.data = foo.metadt
wnv@meta.data$orig.ident = sub('Sample_', '', wnv@meta.data$orig.ident)

# %% [code]
wnv@meta.data[sample(nrow(wnv@meta.data), size=5, replace=F), ]

# %% [markdown]
## Determine % Mitochondrial Reads

# %% [code]
#Find percent mitochondrial genes 
mito.genes = grep(pattern = "^MT-", x = rownames(x = wnv@assays$RNA), value = T)
percent.mito = Matrix::colSums(wnv@assays$RNA[mito.genes, ]) / Matrix::colSums(wnv@assays$RNA)
wnv = AddMetaData(object = wnv, metadata = percent.mito, col.name = "percent.mito")

# %% [markdown]
## Filter out high mitochondrial read cells

# %% [code]
plot1 = FeatureScatter(wnv, feature1 = "nCount_RNA", feature2 = "percent.mito")
plot2 = FeatureScatter(wnv, feature1 = "nCount_RNA", feature2 = "nFeature_RNA")
plot1 + NoLegend()
plot2 + NoLegend()

# %% [code]
# Filtering
wnv = subset(wnv, subset = percent.mito < .25)
wnv = subset(wnv, subset = nCount_RNA < 40000)
wnv = subset(wnv, subset = nCount_RNA > 500)

dim(wnv)

# %% [code]
VlnPlot(wnv, features = c("nFeature_RNA"), pt.size = 0, group.by = 'orig.ident') + NoLegend()
VlnPlot(wnv, features = c("nCount_RNA"), pt.size = 0, group.by = 'orig.ident') + NoLegend()
VlnPlot(wnv, features = c("percent.mito"), pt.size = 0, group.by = 'orig.ident') + NoLegend()

# %% [markdown]
## Log normalize

# %% [code]
#Log normalize and scale x10000
wnv = NormalizeData(wnv, normalization.method = "LogNormalize", scale.factor = 10000)

# %% [markdown]
## Functions for normal log normalization processing

# %% [code]
#Find variable features and scale data
prep.for.cluster = function(ob, features){
    ob = FindVariableFeatures(ob, selection.method = "vst", nfeatures = features)
    top10 = head(VariableFeatures(ob), 20)
    
    plot1 = VariableFeaturePlot(ob)
    plot2 = LabelPoints(plot = plot1, points = top10, repel = T)
    print(plot2)
    
    ob = ScaleData(ob, features = rownames(ob))
    ob = RunPCA(ob, features = VariableFeatures(object = ob))
    
    print(DimPlot(ob, reduction = "pca") + NoLegend())
    print(ElbowPlot(ob, ndims = 50))
    print((((ob@reductions$pca@stdev)^2 / sum((ob@reductions$pca@stdev)^2))*100))
    
    return(ob)
}

#find nearest neighbors and clusters --> run UMAP
cluster = function(subset, dim, res, kpar){
    subset = FindNeighbors(subset, dims = c(1:dim), k.param = kpar)
    subset = FindClusters(subset, resolution = res,random.seed = 1996)
    subset = RunUMAP(subset, dims = c(1:dim), seed.use = 1996)
    return(subset)
}

#subset markers found in differential expression
heatmarkers = function(subset.markers, n_shown){
    subset.markers %>% group_by(cluster) %>% top_n(n = 2, wt = avg_logFC)
    top10.subset = subset.markers %>% group_by(cluster) %>% top_n(n = n_shown, wt = avg_logFC)
    return(top10.subset)
}

# %% [markdown]
## Get variable genes, scale data, PCA

# %% [code]
system.time({
    wnv = prep.for.cluster(wnv, 3000)
})

# %% [markdown]
## Choose number of PCs and k.param

# %% [code]
#Get percent variance of PCA
pca_explained = ((wnv@reductions$pca@stdev)^2 / sum((wnv@reductions$pca@stdev)^2)) * 100

#Explain 75% variance in X number of PCs
i = 1
while(sum(pca_explained[1:i]) < 75) {
    i = i + 1
}
pc_use = i

#Use k-param of 1/2*sqrt(# of cells)
kparam = floor(0.5 * sqrt(dim(wnv)[2]))

message("pc_use: ", pc_use)
message("kparam: ", kparam)

# %% [markdown]
## KNN, SNN, cluster, UMAP

# %% [code]
system.time({
    wnv = cluster(wnv, dim = pc_use, res = 0.5, kpar = kparam)
})

# %% [code]
DimPlot(wnv, group.by = 'U19.ID')
DimPlot(wnv, group.by = 'timepoint')
DimPlot(wnv, group.by = 'disease_state')

# %% [code]
DimPlot(wnv, label = T, repel = T, label.size = 8) + NoLegend()

# %% [code]
#How many cells per condition?
(table_object = table(wnv@meta.data$patient_timepoint))

# %% [code]
#show proportions of cells by cluster
show_proportions = function(object){    
    table_object = table(object@meta.data$seurat_clusters, object@meta.data$patient_timepoint)
    table_object

    Col_object = colSums(table_object)
    for (col in colnames(table_object)){
        table_object[,col] = (table_object[,col]/Col_object[col])*100
    }
    df_object = as.data.frame(table_object)
    colnames(df_object) = c("State","Treatment","Percent")

    df_object[,"Percent"] = as.numeric(df_object[,"Percent"])
    head(df_object)
    g = ggplot() + geom_bar(aes(y = Percent, x = Treatment, fill = State), data = df_object, stat="identity") 
    print(g + theme(axis.text.x = element_text(angle = 90, hjust = 1)))}

# %% [code]
show_proportions(wnv)

# %% [markdown]
## Plot some canonical markers

# %% [code]
FeaturePlot(object = wnv, reduction = 'umap', features = c("MS4A1", "GNLY", "CD3E", 
                                                           "CD14", "FCER1A", "FCGR3A", 
                                                           "LYZ", "TRAC", "CD8A")) + NoLegend()

# %% [markdown]
## Differential expression test between clusters

# %% [code]
wnv.down = SubsetData(wnv, max.cells.per.ident = 200)
wnv.down

# %% [code]
#Find DE markers using Wilcoxon test
system.time({
    wnv.markers = FindAllMarkers(wnv.down, only.pos = T, min.pct = 0.25, logfc.threshold = 0.25)
})

# %% [code]
#produce heatmap 
wnv.features = heatmarkers(subset.markers = wnv.markers, n_shown = 4)

#wnv.down = SubsetData(wnv, max.cells.per.ident = 100)
DoHeatmap(object=wnv.down, features = wnv.features$gene, label = F)

# %% [markdown]
## Subset low quality clusters and high quality clusters based on the heatmap (N.B. increase cluster resolution to differentiate plasma DC from plasma B)

# %% [code]
low_qual = subset(wnv, idents = c(4)) # no specific phenotype, high mitochondrial, low nfeature, low ngene
low_qual

# %% [code]
wnv = subset(wnv, idents = c(0:3, 5:15))
wnv

# %% [code]
wnv = prep.for.cluster(wnv, 3000)

# %% [code]
#Get percent variance of PCA
pca_explained = ((wnv@reductions$pca@stdev)^2 / sum((wnv@reductions$pca@stdev)^2)) * 100

#Explain 75% variance in X number of PCs
i = 1
while(sum(pca_explained[1:i]) < 75){
    i = i + 1
}
pc_use = i

#Use k-param of 1/2*sqrt(# of cells)
kparam = floor(0.5 * sqrt(dim(wnv)[2]))

message("pc_use: ", pc_use)
message("kparam: ", kparam)

# %% [code]
wnv = cluster(wnv, dim = pc_use, res = 1.5, kpar = kparam) #increased resolution

# %% [code]
DimPlot(wnv, group.by = 'U19.ID')
DimPlot(wnv, group.by = 'timepoint')
DimPlot(wnv, group.by = 'disease_state')

# %% [code]
DimPlot(wnv, group.by = 'seurat_clusters', label = T, repel = T, label.size = 8) + NoLegend()

# %% [code]
table_object = table(wnv@meta.data$patient_timepoint)
table_object
show_proportions(wnv)

# %% [code]
wnv.down = SubsetData(wnv, max.cells.per.ident = 200)

#Find DE markers using Wilcoxon test
wnv.markers = FindAllMarkers(wnv.down, only.pos = T, min.pct = 0.25, logfc.threshold = 0.25)

# %% [code]
#produce heatmap 
wnv.features = heatmarkers(subset.markers = wnv.markers, n_shown = 4)

#wnv.down = SubsetData(wnv, max.cells.per.ident = 100)
DoHeatmap(object = wnv.down, features = wnv.features$gene, label = F)

# %% [code]
marker.set = c("MS4A1", "BANK1", "CD79A", #B Cells high
               "NKG7", "GNLY", "FCGR3A", "GZMA", #NK Cells high
               "CD3D", "CD3E", "TRAC",  #NK Cells low
               "CD8A", "CD8B", "CD6", "CD3E", "CD3D", "TCF7", "GZMA", #CD8 T Cells high
               "FCGR3A", #CD8 T Cells Low
               "CD4", "CD6", "CD3E", "CD3D", "TCF7", #CD4 T cells low
               "CD14", "LYZ", "VCAN", "S100A9", "CD68", #CD14 monocytes high
               "FCGR3A", #CD14 monocytes low
               "FCGR3A", "LYZ", "VCAN", "S100A9", "CD68", #CD16 monocytes high
               "FCER1A", "CD1C", "CD74", "HLA-DPA1", "LYZ", #Dendritic Cells high
               "PPBP", "TUBB1", "SDPR", #Platelet/Megakaryocyte high
               "MS4A1", "IGJ", "IGKC", "IGHG1", #Plasma B cells high
               "MKI67", "TOP2A", #Cycling/Proliferating cells high
               "FCGR3B", "CXCR2", "SOD2", #Granulocytes high
               "FOXP3", "TIGIT", #Tregs high within T cells subclustering)
               "GZMB", "IGJ", "SERPINF1", "ITM2C", "IRF7", "IRF8") #pDCs high

# %% [code]
#Plot canonical markers
x = DotPlot(wnv, features = unique(marker.set)) + RotatedAxis()

data = x$data
options(repr.plot.width = 10, repr.plot.height = 10)
ggplot(data, aes(x=features.plot,y=id,size=pct.exp,color=avg.exp.scaled)) + 
geom_point(alpha = 0.8) + 
theme(axis.text.x = element_text(angle = 90)) + 
scale_size(range = c(0, 5)) +
scale_color_gradient(low = "black", high = "red2") +
xlab("") +
ylab("")

# %% [markdown]
## Identify high quality cell types (generalized)

# %% [code]
wnv = SetIdent(wnv, value = 'seurat_clusters')

new.cluster.ids = c("CD4+_T_Cells", "B_Cells", "CD4+_T_Cells", "CD8+_T_Cells", "CD14+_Monocytes",
                    "CD4+_T_Cells", "CD14+_Monocytes", "NK_Cells", "CD14+_Monocytes", "CD14+_Monocytes",
                    "CD14+_Monocytes", "CD8+_T_Cells", "CD14+_Monocytes", "CD16+_Monocytes", "cDCs",
                    "B_Doublets", "Granulocytes", "Platelets/Megakaryocytes", "Proliferating_T_Cells",
                    "CD14+_Monocytes", "pDCs", "Monocytes_Doublets", "Plasma_B_Cells")
                    
names(new.cluster.ids) = levels(wnv)

wnv = RenameIdents(wnv, new.cluster.ids)

wnv$cell_types = Idents(wnv)

DimPlot(wnv, label = T, repel = T, label.size = 8) + NoLegend()

# %% [markdown]
# Subclustering to Detect Doublets and Tregs

# %% [markdown]
## T/NK Cell Labeling and Doublet Finding

# %% [code]
Ts = subset(wnv, idents = c('CD4+_T_Cells', 'CD8+_T_Cells', 'NK_Cells', 'Proliferating_T_Cells'))

# %% [code]
Ts = prep.for.cluster(Ts, features = 3000)

# %% [code]
#Get percent variance of PCA
pca_explained = ((Ts@reductions$pca@stdev)^2/sum((Ts@reductions$pca@stdev)^2))*100

#Explain 75% variance in X number of PCs
i = 1
while(sum(pca_explained[1:i]) < 75){
    i = i + 1
}
pc_use = i

#Use k-param of 1/2*sqrt(# of cells)
kparam = floor(0.5 * sqrt(dim(Ts)[2]))

message("pc_use: ", pc_use)
message("kparam: ", kparam)

# %% [code]
Ts = cluster(Ts, dim = pc_use, res = 0.5, kpar = kparam)

# %% [code]
DimPlot(Ts, label = T, repel = T, label.size = 8) + NoLegend()

# %% [code]
wnv.down = SubsetData(Ts, max.cells.per.ident = 200)

#Find DE markers using Wilcoxon test
wnv.markers = FindAllMarkers(wnv.down, only.pos = T, min.pct = 0.25, logfc.threshold = 0.25)

# %% [code]
#produce heatmap 
wnv.features = heatmarkers(subset.markers = wnv.markers, n_shown = 4)

#wnv.down = SubsetData(wnv, max.cells.per.ident = 100)
DoHeatmap(object=wnv.down, features = wnv.features$gene, label = F)

# %% [code]
x = DotPlot(Ts, features = unique(marker.set)) + RotatedAxis()

data = x$data
options(repr.plot.width = 10, repr.plot.height = 10)
ggplot(data, aes(x=features.plot,y=id,size=pct.exp,color=avg.exp.scaled)) + 
geom_point(alpha = 0.8) + 
theme(axis.text.x = element_text(angle = 90)) + 
scale_size(range = c(0, 5)) +
scale_color_gradient(low = "black", high = "red2") +
xlab("") +
ylab("")

# %% [code]
Ts = SetIdent(Ts, value = 'seurat_clusters')

new.cluster.ids = c("CD4+_T_Cells", "CD8+_T_Cells", "CD4+_T_Cells", "NK_Cells", "CD4+_T_Cells",
                    "CD4+_T_Cells", "CD8+_T_Cells", "NK_Cells", "CD4+_T_Cells", "Regulatory_CD4+_T_Cells",
                    "Proliferating_T_Cells", "CD8+_T_Cells", "T_Doublets", "CD8+_T_Cells")

names(new.cluster.ids) = levels(Ts)

Ts = RenameIdents(Ts, new.cluster.ids)

Ts$cell_types = Idents(Ts)

DimPlot(Ts, label = T, repel = T, label.size = 8) + NoLegend()

# %% [markdown]
## B Labeling and Doublet Finding

# %% [code]
Bs = subset(wnv, idents = c('B_Cells', 'Plasma_B_Cells'))

# %% [code]
Bs = prep.for.cluster(Bs, features = 3000)

# %% [code]
#Get percent variance of PCA
pca_explained = ((Bs@reductions$pca@stdev)^2/sum((Bs@reductions$pca@stdev)^2))*100

#Explain 75% variance in X number of PCs
i = 1
while(sum(pca_explained[1:i]) < 75){
    i = i + 1
}
pc_use = i

#Use k-param of 1/2*sqrt(# of cells)
kparam = floor(0.5 * sqrt(dim(Bs)[2]))

message("pc_use: ", pc_use)
message("kparam: ", kparam)

# %% [code]
Bs = cluster(Bs, dim = pc_use, res = 0.5, kpar = kparam)

# %% [code]
DimPlot(Bs, label = T, repel = T, label.size = 8) + NoLegend()

# %% [code]
wnv.down = SubsetData(Bs, max.cells.per.ident = 200)

#Find DE markers using Wilcoxon test
wnv.markers = FindAllMarkers(wnv.down, only.pos = T, min.pct = 0.25, logfc.threshold = 0.25)

# %% [code]
#produce heatmap 
wnv.features = heatmarkers(subset.markers = wnv.markers, n_shown = 4)

#wnv.down = SubsetData(wnv, max.cells.per.ident = 100)
DoHeatmap(object=wnv.down, features = wnv.features$gene, label = F)

# %% [code]
x = DotPlot(Bs, features = unique(marker.set)) + RotatedAxis()

data = x$data
options(repr.plot.width = 10, repr.plot.height = 10)
ggplot(data, aes(x=features.plot,y=id,size=pct.exp,color=avg.exp.scaled)) + 
geom_point(alpha = 0.8) + 
theme(axis.text.x = element_text(angle = 90)) + 
scale_size(range = c(0, 5)) +
scale_color_gradient(low = "black", high = "red2") +
xlab("") +
ylab("")

# %% [code]
Bs = SetIdent(Bs, value = 'seurat_clusters')

new.cluster.ids = c("B_Cells", "B_Cells", "B_Cells", "B_Cells", "B_Cells", "B_Cells", "B_Doublets", "Plasma_B_Cells")

names(new.cluster.ids) = levels(Bs)

Bs = RenameIdents(Bs, new.cluster.ids)

Bs$cell_types = Idents(Bs)

DimPlot(Bs, label = T, repel = T, label.size = 8) + NoLegend()

# %% [markdown]
## DC and Monocyte Labeling and Doublet Finding

# %% [code]
Myeloid = subset(wnv, idents = c('cDCs', 'pDCs', 'CD14+_Monocytes', 'CD16+_Monocytes'))

# %% [code]
Myeloid = prep.for.cluster(Myeloid, features = 3000)

# %% [code]
#Get percent variance of PCA
pca_explained = ((Myeloid@reductions$pca@stdev)^2/sum((Myeloid@reductions$pca@stdev)^2))*100

#Explain 75% variance in X number of PCs
i = 1
while(sum(pca_explained[1:i]) < 75){
    i = i + 1
}
pc_use = i

#Use k-param of 1/2*sqrt(# of cells)
kparam = floor(0.5 * sqrt(dim(Myeloid)[2]))

message("pc_use: ", pc_use)
message("kparam: ", kparam)

# %% [code]
Myeloid = cluster(Myeloid, dim = pc_use, res = 0.5, kpar = kparam)

# %% [code]
DimPlot(Myeloid, label = T, repel = T, label.size = 8) + NoLegend()

# %% [code]
wnv.down = SubsetData(Myeloid, max.cells.per.ident = 200)

#Find DE markers using Wilcoxon test
wnv.markers = FindAllMarkers(wnv.down, only.pos = T, min.pct = 0.25, logfc.threshold = 0.25)

# %% [code]
#produce heatmap 
wnv.features = heatmarkers(subset.markers = wnv.markers, n_shown = 4)

#wnv.down = SuMyeloidetData(wnv, max.cells.per.ident = 100)
DoHeatmap(object=wnv.down, features = wnv.features$gene, label = F)

# %% [code]
x = DotPlot(Myeloid, features = unique(marker.set)) + RotatedAxis()

data = x$data
options(repr.plot.width = 10, repr.plot.height = 10)
ggplot(data, aes(x=features.plot,y=id,size=pct.exp,color=avg.exp.scaled)) + 
    geom_point(alpha = 0.8) + 
    theme(axis.text.x = element_text(angle = 90)) + 
    scale_size(range = c(0, 5)) +
    scale_color_gradient(low = "black", high = "red2") +
    xlab("") +
    ylab("")

# %% [code]
Myeloid = SetIdent(Myeloid, value = 'seurat_clusters')

new.cluster.ids = c("CD14+_Monocytes","CD14+_Monocytes","CD14+_Monocytes","CD14+_Monocytes",
                    "CD16+_Monocytes","CD14+_Monocytes","Monocytes_Doublets","cDCs",
                    "CD14+_Monocytes","CD14+_Monocytes","CD14+_Monocytes","CD14+_Monocytes","pDCs")
                    
names(new.cluster.ids) = levels(Myeloid)

Myeloid = RenameIdents(Myeloid, new.cluster.ids)

Myeloid$cell_types = Idents(Myeloid)

DimPlot(Myeloid, label = T, repel = T, label.size = 8) + NoLegend()

# %% [markdown]
# Relabeling

# %% [markdown]
## Reconcile old and new cell types

# %% [code]
new.labels.myeloid = as.matrix(Myeloid@meta.data$cell_types)
rownames(new.labels.myeloid) = rownames(Myeloid@meta.data)

new.labels.T = as.matrix(Ts@meta.data$cell_types)
rownames(new.labels.T) = rownames(Ts@meta.data)

new.labels.B = as.matrix(Bs@meta.data$cell_types)
rownames(new.labels.B) = rownames(Bs@meta.data)

new.labels = rbind(new.labels.T, new.labels.B, new.labels.myeloid)
colnames(new.labels) = c('new_cell_types')

df = complete.merge(list(wnv@meta.data, new.labels))

# %% [code]
tester = df
colnames(tester)[length(colnames(tester))] = 'new_cell_types'
tester$new_cell_types[tester$new_cell_types == 0] = NA

df = tester
df = df %>% mutate(new_cell_types = coalesce(new_cell_types, cell_types))
df = df[length(colnames(df))]

# %% [code]
df

# %% [code]
wnv = AddMetaData(object = wnv, metadata = df, col.name = "new_cell_types")

# %% [code]
DimPlot(wnv, group.by = 'new_cell_types', label = T, repel = T, label.size = 8) + NoLegend()

# %% [code]
unique(wnv@meta.data$new_cell_types)

# %% [code]
rm(Bs)
rm(Ts)
rm(Myeloid)

wnv = SetIdent(wnv, value = 'new_cell_types')
wnv

# %% [code]
save(wnv, file = "wnv.seqwell.complete.seurat.d211011.Robj")

# %% [markdown]
## Remove non-PBMCs

# %% [code]
system.time({
wnv_cleaned = subset(wnv, idents = c('CD14+_Monocytes', 'CD16+_Monocytes', 'cDCs', 
                                     'CD4+_T_Cells', 'Proliferating_T_Cells', 'pDCs', 
                                     'Plasma_B_Cells', 'NK_Cells', 'B_Cells', 'CD8+_T_Cells',
                                     'Regulatory_CD4+_T_Cells'))
})
wnv_cleaned

# %% [code]
rm(wnv)

# %% [code]
wnv_cleaned = prep.for.cluster(wnv_cleaned, 3000)

# %% [code]
#Get percent variance of PCA
pca_explained = ((wnv_cleaned@reductions$pca@stdev)^2/sum((wnv_cleaned@reductions$pca@stdev)^2))*100

#Explain 75% variance in X number of PCs
i = 1
while(sum(pca_explained[1:i]) < 75){
    i = i + 1
}
pc_use = i

#Use k-param of 1/2*sqrt(# of cells)
kparam = floor(0.5 * sqrt(dim(wnv_cleaned)[2]))

message("pc_use: ", pc_use)
message("kparam: ", kparam)

# %% [code]
wnv_cleaned = cluster(wnv_cleaned, dim = pc_use, res = 0.5, kpar = kparam)

# %% [code]
DimPlot(wnv_cleaned, group.by = 'U19.ID')
DimPlot(wnv_cleaned, group.by = 'timepoint')
DimPlot(wnv_cleaned, group.by = 'disease_state')

# %% [code]
DimPlot(wnv_cleaned, group.by = 'new_cell_types', label = T, repel = T, label.size = 8) + NoLegend()

# %% [code]
wnv_cleaned = SetIdent(wnv_cleaned, value = 'new_cell_types')

# %% [raw]
wnv.down = SubsetData(wnv, max.cells.per.ident = 200)
wnv.down

# %% [code]
#Find DE markers using Wilcoxon test
wnv.markers = FindAllMarkers(wnv.down, only.pos = T, min.pct = 0.25, logfc.threshold = 0.25)

# %% [code]
#produce heatmap 
wnv.features = heatmarkers(subset.markers = wnv.markers, n_shown = 4)

#wnv.down = SubsetData(wnv, max.cells.per.ident = 100)
DoHeatmap(object = wnv.down, features = wnv.features$gene, label = F)

# %% [code]
save(wnv_cleaned, file = "wnv.seqwell.pbmcs.seurat.d211011.Robj")

# %% [markdown]
---

# %% [code]
date()
```
